# MUSSEL: Enhanced Bayesian Polygenic Risk Prediction Leveraging Information across Multiple Ancestry Groups

**DOI:** 10.1101/2023.04.12.536510

**Authors:** Jin Jin, Jianan Zhan, Jingning Zhang, Ruzhang Zhao, Jared O’Connell, Yunxuan Jiang, 23andMe Research Team, Steven Buyske, Christopher Gignoux, Christopher Haiman, Eimear E. Kenny, Charles Kooperberg, Kari North, Bertram L. Koelsch, Genevieve Wojcik, Haoyu Zhang, Nilanjan Chatterjee

## Abstract

Polygenic risk scores (PRS) are now showing promising predictive performance on a wide variety of complex traits and diseases, but there exists a substantial performance gap across different populations. We propose MUSSEL, a method for ancestry-specific polygenic prediction that borrows information in the summary statistics from genome-wide association studies (GWAS) across multiple ancestry groups. MUSSEL conducts Bayesian hierarchical modeling under a MUltivariate Spike-and-Slab model for effect-size distribution and incorporates an Ensemble Learning step using super learner to combine information across different tuning parameter settings and ancestry groups. In our simulation studies and data analyses of 16 traits across four distinct studies, totaling 5.7 million participants with a substantial ancestral diversity, MUSSEL shows promising performance compared to alternatives. The method, for example, has an average gain in prediction R^2^ across 11 continuous traits of 40.2% and 49.3% compared to PRS-CSx and CT-SLEB, respectively, in the African Ancestry population. The best-performing method, however, varies by GWAS sample size, target ancestry, underlying trait architecture, and the choice of reference samples for LD estimation, and thus ultimately, a combination of methods may be needed to generate the most robust PRS across diverse populations.

## Introduction

Polygenic models for predicting complex traits are widely developed utilizing summary-level association statistics from genome-wide association studies (GWAS). While being on course to translate GWAS results into clinical practice, polygenic risk scores (PRS) encounter obstacles due to the poor predictive performance on underrepresented non-European (non-EUR) ancestry populations, especially those with substantial African ancestry^1-4^. As sample sizes for GWAS in many non-EUR populations remain low for many traits, applications of PRS often rely on EUR-based models, which underperform in other populations due in part to differences in allele frequencies, SNP effect sizes, and linkage disequilibrium (LD)^1-3,5,6^.

To improve the poor performance of PRS on non-EUR populations, several multi-ancestry methods have recently been developed to combine information from available GWAS summary statistics and LD reference data across multiple ancestry groups^7^. One simple approach is the weighted PRS^8^, which trains a linear combination of the PRS developed using single-ancestry methods (e.g., LD clumping and P-value thresholding, C+T) applied separately to available GWAS data across different ancestry groups^8^. More recent methods attempt to borrow information across ancestry at the level of individual SNPs based on Bayesian methods^9,10^, penalized regressions^11,12^, or through the extension of C+T^13^. However, applications show that no single method performs uniformly the best, and their performance depends on many aspects, including the underlying genetic architecture of the trait, the absolute and relative sample sizes across populations, and the algorithm for the estimation of LD based on the underlying reference dataset^13^.

We propose MUSSEL, a novel method for developing ancestry-specific PRS by jointly modeling ancestry-specific GWAS summary data across diverse ancestries. The method conducts Bayesian modeling of SNP effect sizes across ancestries via a MUltivariate Spike and Slab prior and an Ensemble Learning (MUSSEL) step to seek an “optimal” combination of a series of PRS obtained under different tuning parameters. We evaluate MUSSEL and benchmark it against a variety of alternatives through large-scale simulation studies and analyses of 16 traits from four different studies: (1) the Population Architecture using Genomics and Epidemiology (PAGE) Study supplemented with data from the Biobank Japan (BBJ) and UK Biobank (UKBB), (2) Global Lipids Genetics Consortium (GLGC), (3) All of Us (AoU), and (4) 23andMe, Inc. These studies, with training data and additional validation samples from the UKBB study, included a total of 3.4 million European (EUR), 226K Admixed African, African, or African American (AFR), 437K Admixed Americans or Hispanic/Latino (AMR), 389K East Asian (EAS), and 56K South Asian (SAS). Results reveal the promising performance of MUSSEL for developing robust PRS in the multi-ancestry setting and identify a number of practical considerations for implementations that are crucial to the performance of the method.

## Results

### MUSSEL Overview

Considering that GWAS summary-level association statistics can be shared much more easily among research teams than individual-level genotype and phenotype data from GWAS, we will focus on PRS methods that can use summary-statistics from the GWAS training samples. The implementation of our proposed method, MUSSEL, as well as other multi-ancestry methods which we will compare MUSSEL to, requires three (ancestry-specific) datasets from each training ancestry group: (1) GWAS summary data, (2) LD reference data, and (3) a validation (tuning + testing) dataset with genotype and phenotype data for an adequate number of individuals that are independent of GWAS samples and LD reference samples.

We now introduce MUSSEL, a novel method for enhanced ancestry-specific polygenic risk prediction based on available GWAS summary-level association statistics and LD reference data across multiple ancestry groups. MUSSEL consists of two steps (Figure 1): (1) a Bayesian modeling step (“MUSS”) to model the genetic correlation structure in SNP effect sizes across ancestry groups while accounting for ancestry-specific LD across SNPs, and (2) an ensemble learning (EL) step via a super learner (SL) to construct an “optimal” linear combination of a series of PRS obtained from MUSS under different tuning parameter settings and across all ancestry groups. Additionally, a step 0 was conducted before step 1 to obtain tuned causal SNP proportion and heritability parameters for each training ancestry group from LDpred2. These parameters will be used to specify the prior causal SNP proportions and heritability parameters in MUSS.

**Figure 1:**
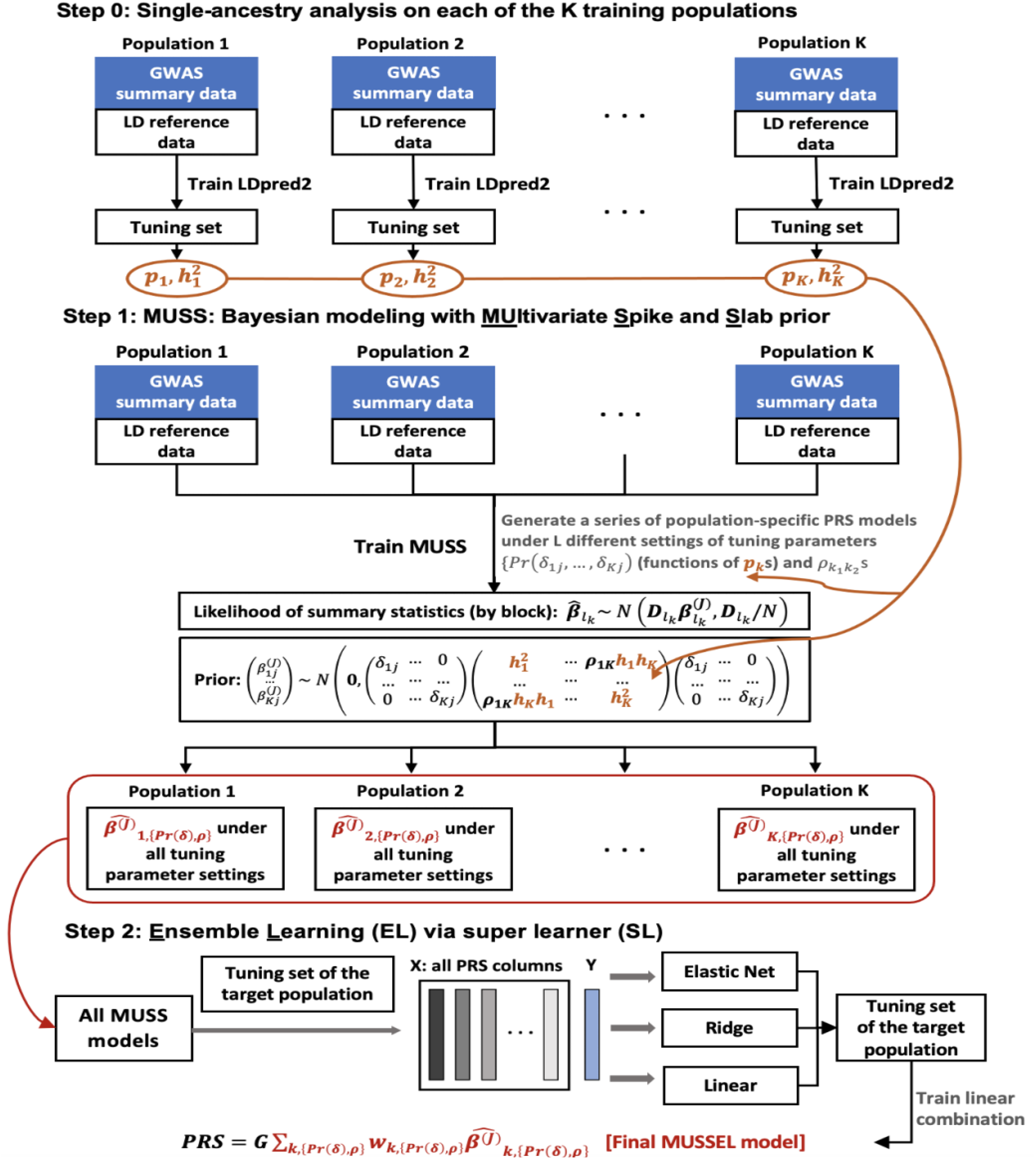
MUSSEL Workflow. [**Step 0**] apply LDpred2 to each of the K training populations (ancestry groups) to obtain estimated causal SNP proportions (*p*_*k*_, *k* = 1, …, *K*) and heritability 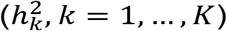 parameters based on the tuning set, these parameters will be used to specify the prior distributions and tuning parameter settings for Bayesian learning with MUSS. [**Step 1**] MUSS: jointly model across all training populations to obtain a total of (L×K) PRS models under L different tuning parameter settings for Pr (*δ*_1*j*_, …, *δ*_*Kj*_) (functions of *p*_*k*_s) and 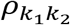 across K training populations. [**Step 2**] for the target population, conduct ensemble learning (EL) via a super learner (SL) algorithm with 3 base learners (elastic net regression, ridge regression, and linear regression) to train an “optimal” linear combination of the (L×K) PRS models from the MUSS step to obtain the final MUSSEL model. The prediction performance of the final PRS derived using MUSSEL should be evaluated on an independent testing set.

**Step 1: MUSS: Bayesian modeling with MUltivariate Spike and Slab prior**

MUSS tailors effect size estimates for each ancestry group by incorporating data from other ancestry groups via Bayesian hierarchical modeling with a multivariate spike and slab prior on SNP effect sizes across ancestry groups. For population-specific SNPs, i.e., SNPs with minor allele frequency (MAF)>0.01 in only one ancestry group, we assume a spike-and-slab prior as in LDpred2. For SNPs that are polymorphic across multiple populations, the between-SNP correlation is induced in two aspects: (1) we assume a SNP is causal in all those populations or none, and (2) the effect sizes for causal SNPs across populations are correlated (see Online Methods for details). The prior specification is distinct compared to the recent method PRS-CSx^9^ in two aspects: (1) the use of a multivariate spike-and-slab prior versus a continuous shrinkage prior to perform shrinkage estimation; and (2) flexible specification of genetic correlation structure across ancestry groups in MUSSEL compared to PRS-CSx, which assumes a single hyperparameter is shared across different ancestry groups and thus incorporates as fairly rigid specification of the correlation structure.

We infer posterior estimates of LD-adjusted SNP effect sizes across different ancestries via an efficient Markov chain Monte Carlo (MCMC) algorithm (Online Methods). Multiple PRS will be developed for each ancestry under carefully designed settings of two sets of tuning parameters, (1) the causal SNP proportion in each ancestry group, which will be used to specify the correlated prior causal probabilities across ancestry groups (Online Methods), and (2) the between-ancestry genetic correlation in SNP effect sizes. Ancestry-specific SNP effect sizes are estimated based on MCMC with an approximation strategy previously implemented in the LDpred2 algorithm^14^, which substantially reduces the number of iterations required to reach convergence with a spike-and-slab type prior on a large number of correlated SNPs. Detailed MCMC algorithm and estimation procedure are described in Online Methods.

**Step 2: Ensemble Learning (EL) via Super Learner (SL)**

Research has shown that combining multiple C+T PRS under different p-value thresholds^15^ or combining the best ancestry-specific PRS across multiple ancestry groups^8,9^ can significantly improve predictive performances. Thus, as a second step of MUSSEL, we consider combining PRS obtained from the MUSS step both across different tuning parameter settings and across ancestry groups via an SL model trained on the tuning dataset. SL is an EL method for seeking an “optimal” linear combination of various base learners for prediction^16^. In our analyses, we consider three linear base learners, including linear regression, elastic net regression^17^, and ridge regression^18^. A similar SL procedure was also implemented recently in another multi-ancestry method CT-SLEB^13^. In our simulation studies and real data examples, we will show explicitly how much improvement in predictive power can be obtained separately through the Bayesian modeling step and the EL step. Considering that both weighted PRS and PRS-CSx construct a linear combination of the best PRS for each ancestry group, we tried the same approach on our Bayesian model (MUSS) and called this alternative method “weighted MUSS”. We observe on both simulated data and real data that the gain in predictive power by this linear combination strategy is mostly lower than, and sometimes comparable to, the gain by our proposed EL strategy. (Supplementary Figures 1-10 and 13-15, “weighted MUSS” versus “MUSSEL”).

### Simulation Settings

We first investigate the performance of MUSSEL and a series of existing methods under various simulated scenarios of the genetic architecture of a continuous trait and absolute and relative GWAS sample sizes across ancestry groups. This large-scale dataset, including simulated genotype and phenotype data for a total of 600,000 individuals across EUR, AFR, AMR, EAS, and SAS, was recently released by our group^13^. Detailed simulation setup is described in Zhang et al. (2022)^13^ and briefly summarized in the Supplementary Notes.

We apply eight existing approaches for comparison, which include two single-ancestry methods applied to GWAS and LD reference data from the target population: (1) C+T, (2) LDpred2; the same single-ancestry methods applied to GWAS and LD reference data for EUR: (3) EUR C+T, (4) EUR LDpred2; and three existing multi-ancestry methods applied to ancestry-specific GWAS and LD reference data for all ancestry groups: (5) weighted C+T (weighted PRS using C+T as the base method), (6) weighted LDpred2 (weighted PRS using LDpred2 as the base method), (7) PRS-CSx^9^, and (8) CT-SLEB^13^. Results from another two recently proposed multi-ancestry methods, PolyPred+^19^ and XPASS^10^, on the same simulated dataset are reported in Zhang et al. (2022)^13^. Table 1 provides a comparison of the various methods in terms of data requirement, similarities, and differences. Taking into account both ancestral diversity and computational efficiency, throughout the text, we restrict all our analyses to the SNPs among approximately 2.0 million SNPs in HapMap 3^20^ plus Multi-Ethnic Genotyping Array (MEGA)^21^ that are also available in the discovery GWAS, LD reference panel, and validation (tuning + testing) samples. We assess the predictive performance of a PRS by prediction R^2^, i.e., the proportion of variance of the trait explained by the PRS. The corresponding 95% bootstrap CIs are calculated based on 10,000 bootstrap samples using the Bca approach^22^ implemented in the R package “boot”^23^ (Supplementary Figures 1–10, Supplementary Tables 1.1 – 1.5). Results from the various methods are compared in five simulation settings: (1) fixed common SNP heritability, strong negative selection, with a genetic correlation set to *ρ* = 0.8 between any two ancestry groups (Figure 2, Supplementary Figures 1-2), (2) fixed per-SNP heritability, strong negative selection, *ρ* = 0.8 (Supplementary Figures 3-4), (3) fixed per-SNP heritability, strong negative selection, with a weaker between-ancestry genetic correlation *ρ* = 0.6 (Supplementary Figures 5-6), (4) fixed common SNP heritability, no negative selection, *ρ* = 0.8 (Supplementary Figures 7-8), and (5) fixed common SNP heritability, mild negative selection, *ρ* = 0.8 (Supplementary Figures 9-10).

**Table 1:** An overview of the methods implemented for PRS development.

| Type | Method | Required training data source<sup>*</sup> | Features | Tuning parameters |
| --- | --- | --- | --- | --- |
| Single-ancestry method | C+T | Target ancestry | Model-free | $p_t$ (p-value threshold) |
| | LDpred2 | Target ancestry | Bayesian (spike-and-slab prior) | $p$ (causal SNP proportion), $h^2$ (heritability) |
| | EUR C+T | EUR | Model-free | $p_t$ (p-value threshold) |
| | EUR LDpred2 | EUR | Bayesian (spike-and-slab prior) | $p$ (causal SNP proportion), $h^2$ (heritability) |
| Multi-ancestry method | Weighted LDpred2 | Ancestry-specific data from each available ancestry | Bayesian (spike-and-slab prior), linear combination strategy | $p$, $H^2$, weight of each ancestry-specific PRS in the final model |
| | PRS-CSx | | Bayesian (Strawderman–Berger prior), linear combination strategy | $\phi$ (global shrinkage parameter), weight of each ancestry-specific PRS in the final model |
|  | XPASS<sup>**</sup> |  | Bayesian (bivariate normal prior), infinitesimal model |  |
|  | PolyPred+<sup>**</sup> |  | Bayesian, functional annotation, linear combination of SBayesR and PolyFun | Parameters in SBayesR and PolyFun, weight of SBayesR PRS and PolyFun PRS in the final model |
| | CT-SLEB | | Empirical Bayes, EL via SL | $p_t$ (p-value threshold), $d$ (genetic distance) for C+T step, parameters in the SL |
| | MUSSEL | | Bayesian (multi-variate spike-and-slab prior), EL via SL | $\Pr(\delta_1, \dots, \delta_K), \rho_{k_1 k_2}, 1 \leq k_1 < k_2 \leq K$, parameters in the SL |

<sup>\*</sup>Note: all methods require three datasets to train the PRS model: (1) discovery GWAS summary data, (2) LD reference data, and (3) tuning data. <sup>\*\*</sup> Results from PolyPred+ and XPASS on all simulated and real datasets (except for PAGE + UKBB + BBJ) were reported in Zhang et al. (2022)<sup>12</sup>.

**Figure 2:**
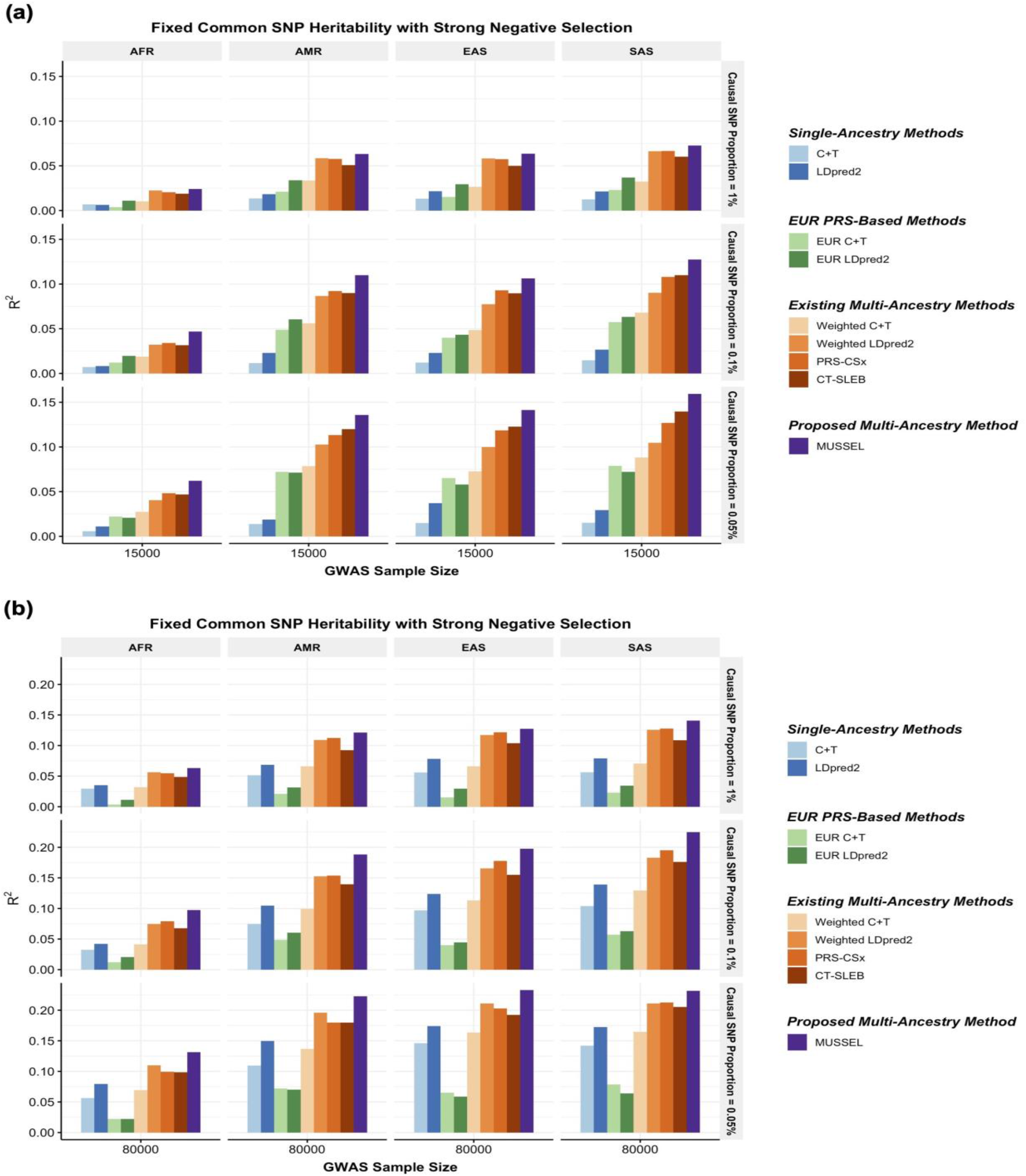
**Simulation results showing performance of the PRS constructed by MUSSEL and various existing methods, assuming a fixed common SNP heritability (0.4) across ancestries under a strong negative selection model for the relationship between SNP effect size and allele frequency**. The genetic correlation in SNP effect size is set to 0.8 across all pairs of populations. The causal SNP proportion (degree of polygenicity) is set to 1.0%, 0.1%, or 0.05% (∼192*K*, 19.2*K*, or 9.6*K* causal SNPs). We generate data for ∼19 million common SNPs (MAF≥1%) across the five ancestries but conduct analyses only on the ∼2.0 million SNPs in HapMap 3 + MEGA. The PRS-CSx software only considers approximately 1.2 million HapMap 3 SNPs and therefore we report the performance of PRS-CSx PRS only based on the HapMap 3 SNPs. The discovery GWAS sample size is set to **(a)** 15,000 or **(b)** 80,000 for each non-EUR ancestry, and 100,000 for EUR. A tuning set consisting of 10,000 individuals is used for parameter tuning and training the SL in CT-SLEB and MUSSEL or the linear combination model in weighted C+T, weighted LDpred2, and PRS-CSx. The reported *R*^2^ values are calculated on an independent testing set of 10,000 individuals for each ancestry group.

**Figure 3:**
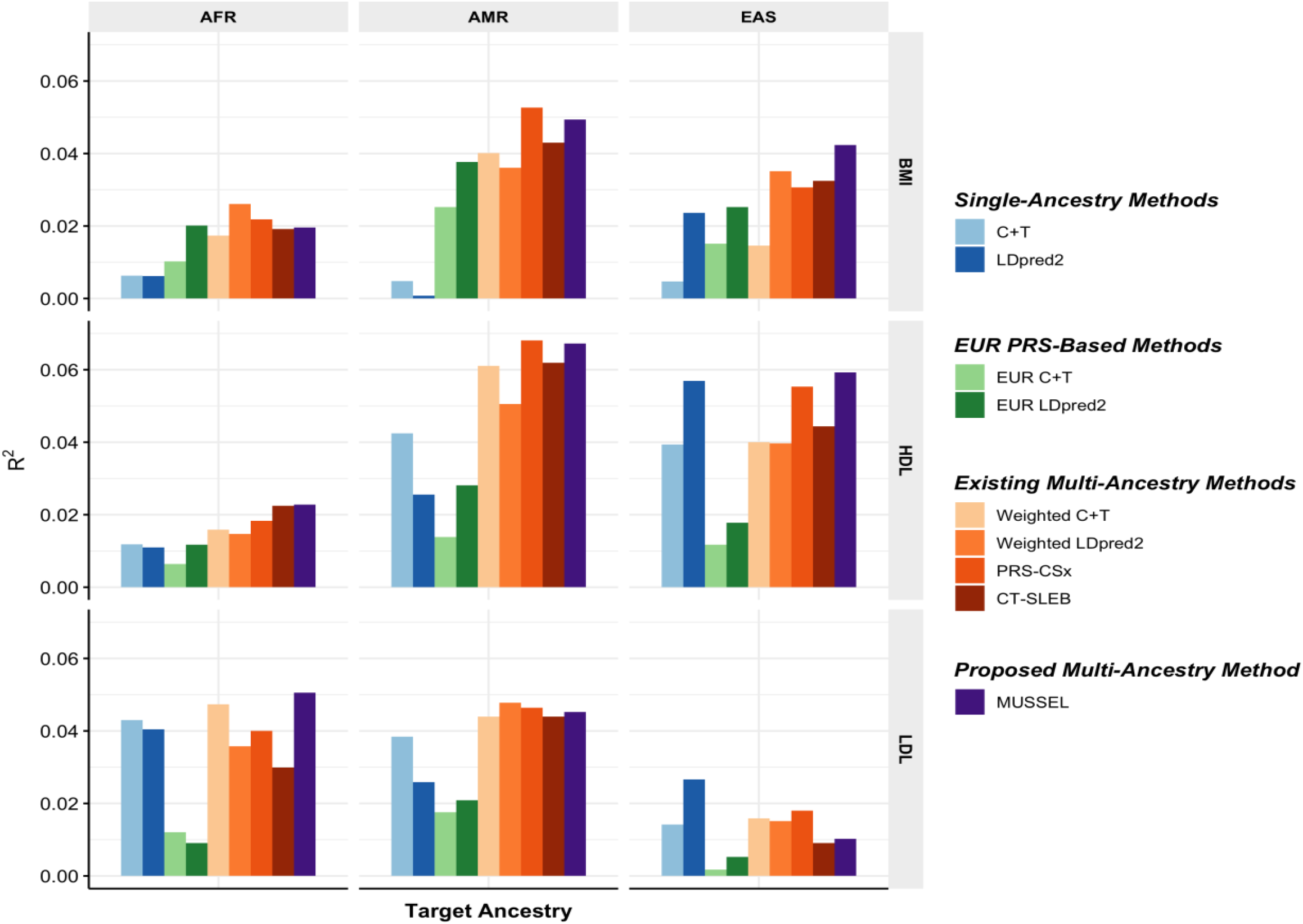
**Prediction R^2^ on validation individuals of AFR (N=2,015–3,428), EAS (N=2,316-4,647), and AMR ancestries (N=3,479-4,397) in PAGE based on discovery GWAS from PAGE (AFR N_GWAS_=7,775 – 13,699, AMR N_GWAS_=13,894 – 17,558), BBJ (EAS N_GWAS_=70,657 – 158,284), and UKBB (EUR N_GWAS_=315,133 – 355,983).** We used genotype data from 1000 Genomes Project (498 EUR, 659 AFR, 347 AMR, 503 EAS, 487 SAS) as the LD reference dataset. All methods were evaluated on the ∼2.0 million SNPs that are available in HapMap 3 + MEGA, except for PRS-CSx which is evaluated based on the HapMap 3 SNPs only, as implemented in their software. Ancestry- and trait-specific GWAS sample sizes, number of SNPs included, and validation sample sizes are summarized in Supplementary Table 3.1. A random half of the validation individuals is used as the tuning set to tune model parameters, as well as train the SL in CT-SLEB and MUSSEL or the linear combination model in weighted C+T, weighted LDpred2, and PRS-CSx. The other half of the validation set is used as the testing set to report R^2^ values for PRS on each ancestry, after adjusting for whether the sample is from BioMe and the top 10 genetic principal components for BMI, and additionally the age at lipid measurement and sex. The 95% bootstrap CIs of the estimated R^2^ are reported in Supplementary Figure 13 and Supplementary Table 9.

**Figure 4:**
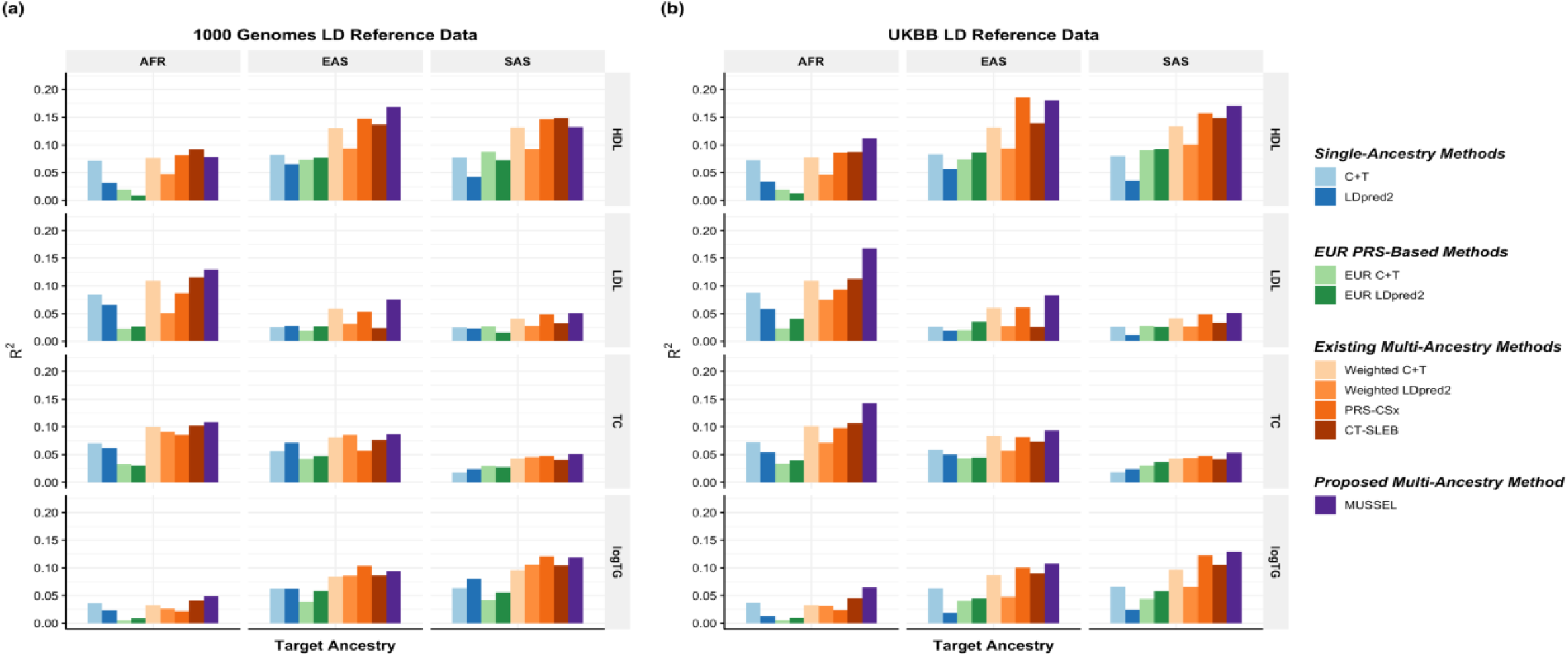
**Prediction R^2^ on UKBB validation individuals of EUR (17,457 – 19,030), AFR (7,954 – 8,598), EAS (1,752 – 1,921), and SAS (9,385 – 10,288) origin based on discovery GWAS from GLGC on EUR (N_GWAS_ =842,660 – 930,671), AFR or admixed AFR (N_GWAS_ =87,760 – 92,555), Hispanic/Latino (N_GWAS_ =46,040 – 49,582), EAS (N_GWAS_ =82,587 – 146,492), and SAS (N_GWAS_ =33,s658 – 34,135).** The LD reference data are from either **(a)** 1000 Genomes Project (498 EUR, 659 AFR, 347 AMR, 503 EAS, 487 SAS), or **(b)** UKBB data (PRS-CSx: default UKBB LD reference data which overlaps with our testing samples including 375,120 EUR, 7,507 AFR, 687 AMR, 2,181 EAS, and 8,412 SAS; all other methods: UKBB tuning samples including 10,000 EUR, 4,585 AFR, 1,010 EAS, and 5,427 SAS). The ancestry of UKBB individuals were determined by a genetic ancestry prediction approach (Supplementary Notes). Due to the low prediction accuracy of genetic component analysis and extremely small validation sample size of UKBB AMR, prediction R^2^ on UKBB AMR is unreliable and thus is not reported here. All methods were evaluated on the ∼2.0 million SNPs that are available in HapMap 3 + MEGA, except for PRS-CSx which is evaluated based on the HapMap 3 SNPs only, as implemented in their software. Ancestry- and trait-specific GWAS sample sizes, number of SNPs included, and validation sample sizes are summarized in Supplementary Table 4.1. A random half of the validation individuals is used as the tuning set to tune model parameters, as well as train the SL in CT-SLEB and MUSSEL or the linear combination model in weighted LDpred2, PRS-CSx, and weighted MUSS. The other half of the validation set is used as the testing set to report R^2^ values for each ancestry, after adjusting for age, sex, and the top 10 genetic principal components. In (b), PRS-CSx and other methods do not have a fair comparison because the UKBB LD reference data provided by the PRS-CSx software (UKBB_PRS-CSx_) is much larger than that for other methods, and thus the R^2^ of PRS-CSx PRS may be inflated due to a big overlap between UKBB_PRS-CSx_ and the UKBB testing sample. The 95% bootstrap CIs of the estimated R^2^ are reported in Supplementary Figure 14 and Supplementary Table 9.

**Figure 5:**
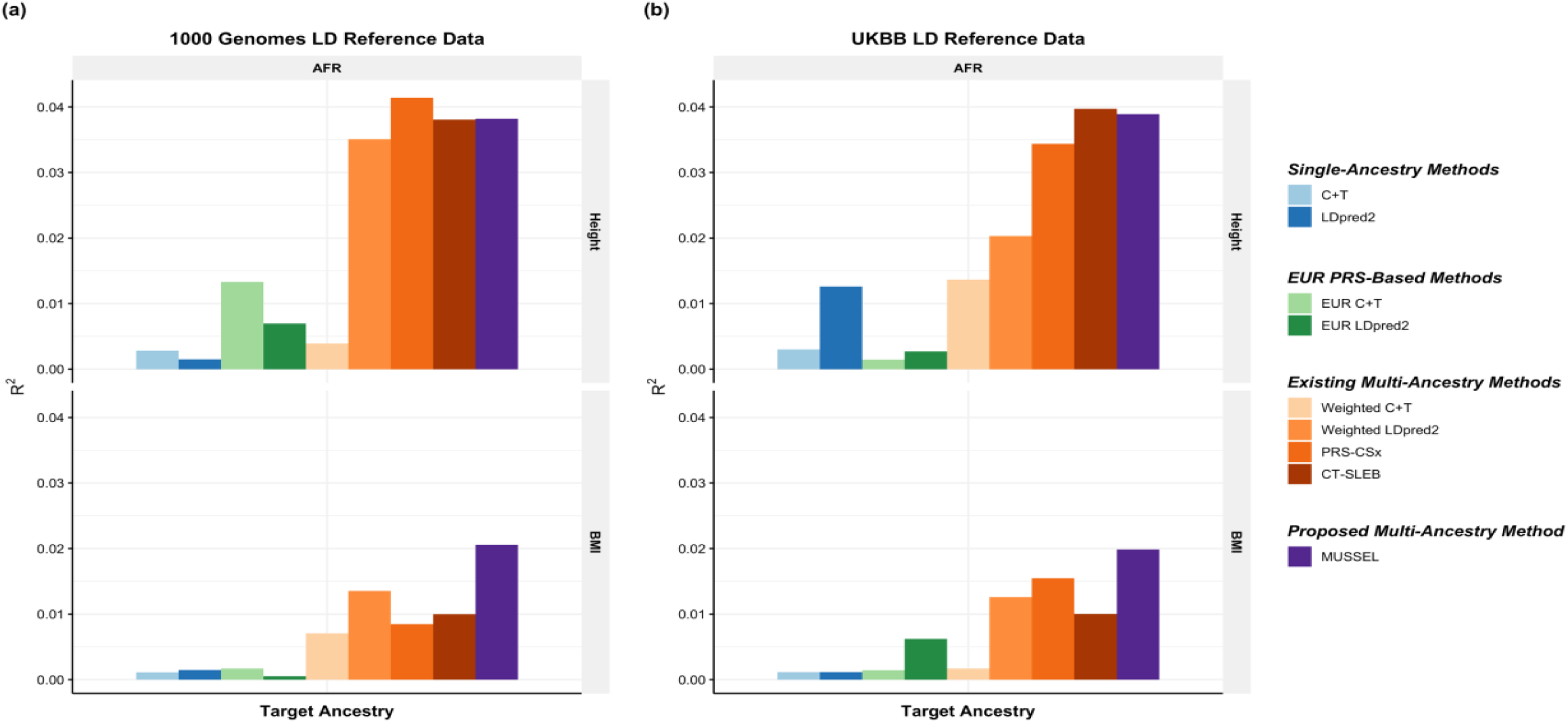
**Prediction R**^**2**^ **on UKBB validation individuals of AFR (N=9,026 – 9,042) origin based on discovery GWAS from AoU on EUR (N**_**GWAS**_ **=48,229 – 48,332), AFR (N**_**GWAS**_ **=21,514 – 21,550), and Hispanic/Latino (N**_**GWAS**_ **=15,364 – 15,413)**. The LD reference data is either **(a)** 1000 Genomes Project (498 EUR, 659 AFR, 347 AMR, 503 EAS, 487 SAS), or **(b)** UKBB data (PRS-CSx: default UKBB LD reference data which overlap with our testing samples including 375,120 EUR, 7,507 AFR, 687 AMR, 2,181 EAS, and 8,412 SAS; all other methods: UKBB tuning samples including 10,000 EUR, 4,585 AFR, 1,010 EAS, and 5,427 SAS). The ancestry of UKBB individuals were determined by a genetic ancestry prediction approach (Supplementary Notes). Due to the low prediction accuracy of genetic component analysis and extremely small validation sample size of UKBB AMR, prediction R^2^ on UKBB AMR is unreliable and thus is not reported here. All methods were evaluated on the ∼2.0 million SNPs that are available in HapMap3 + MEGA, except for PRS-CSx which is evaluated based on the HapMap 3 SNPs only, as implemented in their software. Ancestry- and trait-specific sample sizes of GWAS, number of SNPs included, and validation sample sizes are summarized in Supplementary Table 5.1. A random half of the validation individuals is used as the tuning set to tune model parameters, as well as train the SL in CT-SLEB and MUSSEL or the linear combination model in weighted LDpred2, PRS-CSx, and weighted MUSS. The other half of the validation set is used as the testing set to report R^2^ values for each ancestry, after adjusting for age, sex, and the top 10 genetic principal components. In (b), PRS-CSx and other methods do not have a fair comparison because the UKBB LD reference data provided by the PRS-CSx software (UKBB_PRS-CSx_) is much larger than that for other methods, and thus the R^2^ of PRS-CSx may be inflated due to a big overlap between UKBB_PRS-CSx_ and the UKBB testing sample. The 95% bootstrap CIs of the estimated R^2^ are reported in Supplementary Figure 15 and Supplementary Table 9.

### Simulation results

The multi-ancestry methods tend to outperform the single-ancestry methods, except for weighted C+T, which performs worse than LDpred2 when GWAS sample size of the non-EUR target population becomes adequately large (Figure 2, Supplementary Figures 1-10). When the discovery GWAS sample size of the target non-EUR population is relatively small (N=15,000) compared to EUR GWAS (N=100,000), EUR PRS tends to outperform PRS generated based on training data from the target non-EUR population; but as GWAS sample size of the target non-EUR population increases, the prediction R^2^ of LDpred2 eventually becomes substantially higher than that of EUR C+T and EUR LDpred2. Among the existing multi-ancestry methods, weighted LDpred2, PRS-CSx, and CT-SLEB perform similarly but show advantages over others in different settings: weighted LDpred2 performs well in the scenario of a large causal SNP proportion, CT-SLEB performs similarly as PRS-CSx but shows some advantages when there is a small causal SNP proportion (0.05%) and when GWAS sample size for target non-EUR population is small. Overall, the proposed method MUSSEL outperforms these existing methods in almost all settings. This is expected given that the SNP effect sizes were simulated under a multivariate spike-and-slab distribution as assumed in the MUSS model. The proposed EL step (in MUSSEL) and the alternative linear combination step (in weighted MUSS) only provide minimal improvement in R^2^ on top of MUSS (Supplementary Figures 1-10). This may be because when the specified distribution of SNP effect sizes approximates the true distribution well, the best PRS trained for each ancestry by MUSS can already provide a high predictive power, and an additional step of combining PRS across tuning parameter settings and ancestry groups is unnecessary.

We also checked computation intensity of MUSSEL in comparison with PRS-CSx. A comparison of computation time between PRS-CSx and CT-SLEB on the same simulation dataset was reported in Zhang et al. (2022)^13^. With AMD EPYC 7702 64-Core Processors running at 2.0 GHz using a single core, on chromosome 22 and with a total of 5 × (K+1) tuning parameter settings, MUSSEL has an average runtime of approximately 75.9 minutes combining *K*=3 ancestry groups with a total of 17,192 SNPs, 127.2 minutes combining *K*=4 ancestry groups with 17,721 SNPs, and 237.4 minutes across *K*=5 ancestry groups with 17,722 SNPs. Although not as fast as simpler methods such as CT-SLEB and XPASS, MUSSEL is computationally more efficient than PRS-CSx (K=3: 3.8-fold, K=4: 3.2-fold, K=5: 2.5-fold) and thus is easier to implement than PRS-CSx especially when four or more training populations are available to be combined.

To examine whether the performance of MUSSEL is sensitive to misspecification of the LD matrix, we conduct a sensitivity analysis, where we estimate LD for each ancestry group based on a slightly mis-specified LD reference sample that contains 800 individuals from the same ancestry group and 50 individuals from each of the other four ancestry groups, totaling 200 (20%) individuals with ancestry mismatch. We repeat our analysis under the setting of having fixed common SNP heritability, a strong negative selection, and a genetic correlation of 0.8 across all pairs of ancestry groups, based on the mis-specified LD reference samples. We also apply LDpred2, EUR LDpred2, and weighted LDpred2, which may also be sensitive to ancestry mismatch between the discovery GWAS samples and LD reference samples. Compared to the results assuming no ancestry mismatch between the discovery GWAS and LD reference data, the R^2^ of LDpred2, EUR LDpred2, weighted LDpred2, and MUSSEL PRS are on average 3.3%, 5.7%, 15.1%, and 10.4% lower, respectively (Supplementary Figures 11 – 12, Supplementary Table 8). The amount of power loss appears to increase as the underlying causal SNP proportion decreases.

### PAGE + UKBB + BBJ data analysis with validation on non-EUR individuals from PAGE

We evaluate the performance of the various methods on predicting the polygenic risk of inverse-rank normal transformed BMI (IRNT BMI), high-density lipoprotein (HDL), and low-density lipoprotein (LDL) separately for AFR, AMR, and EAS. We collected ancestry-specific training GWAS summary data for AFR and AMR from PAGE, GWAS summary data for EAS from BBJ, and EUR GWAS summary data from UKBB. The PRS developed by the various methods are evaluated on validation individuals of AFR, AMR, and EAS populations from PAGE. We use genotype data for 498 EUR, 659 AFR, 347 AMR, 503 EAS, and 487 SAS individuals from the 1000 Genomes project as the LD reference data^24^.

In this set of analyses, we observe that the multi-ancestry methods tend to outperform single-ancestry methods for EUR, AFR, and AMR (Figure 3, Supplementary Figure 13, Supplementary Table 3). For EAS, LDpred2 can reach an R^2^ similar to or higher than that of EUR LDpred2 and multi-ancestry methods, which is possibly because the BBJ GWAS sample sizes for EAS are relatively large (N=70,657 – 158,284). For the proposed method MUSSEL, we observe potential improvement in R^2^ from both the Bayesian modeling step (“MUSS” versus “LDpred2”) and the SL step (“MUSSEL” versus “MUSS”). The linear combination strategy (“weighted MUSS”, Supplementary Figure 13) provides a smaller or similar gain in R^2^ compared to our SL strategy (“MUSSEL”). The relative performance of the various multi-ancestry methods varies by trait and ancestry, and no method is uniformly better than others. In some settings, MUSSEL PRS gives a lower R^2^ than the PRS trained by weighted LDpred2 and PRS-CSx in some settings, such as for BMI on AFR and LDL on EAS. But in general, the MUSSEL PRS has the best overall performance, with an average increase of 3.6% and 19.6% in R^2^ compared to PRS-CSx and CT-SLEB, respectively, on non-EUR ancestries.

### GLGC data analysis with validation on UKBB individuals

We apply the various methods to develop ancestry-specific PRS for four blood lipid traits, including HDL, LDL, total cholesterol (TC), and log of triglycerides (logTG)^25^, based on ancestry-specific GWAS summary data for EUR, AFR, AMR, EAS, and SAS, from the Global Lipids Genetics Consortium (GLGC). We validate the performance of the various methods on UKBB individuals of AFR, EAS, and SAS origin separately, where the ancestry information of the UKBB validation individuals was determined based on an ancestry genetic component analysis (Supplementary Notes).

We first use genotype data of the unrelated 1000 Genomes samples as the LD reference data^24^. We observe that the MUSSEL PRS performs the best or similarly to the best PRS (Figure 4(a), Supplementary Figure 14(a), and Supplementary Table 4). We see a notable gain in R^2^ comparing MUSSEL PRS to weighted LDpred2 PRS (average increase: 50.7%). MUSSEL outperforms CT-SLEB in most cases (average increase in R^2^: 27.1%). Although the relative performance between MUSSEL and PRS-CSx varies by ancestry and trait, MUSSEL PRS has a better overall performance, with an average increase of 19.9% in R^2^ compared to PRS-CSx PRS. Similar to the results from PAGE + UKBB + BBJ analysis, MUSSEL improves on top of LDpred2 by both the Bayesian modeling step (“MUSS” versus “LDpred2”, Supplementary Figure 14(a)) and the SL step (“MUSSEL” versus “MUSS”, Supplementary Figure 14(a)). The PRS generated by the alternative linear combination strategy has a similar or lower R^2^ than the PRS generated by our proposed EL strategy (“weighted MUSS” versus “MUSSEL”, Supplementary Figure 14(a)).

It has been observed that LDpred2 sometimes has suboptimal performance based on the widely implemented 1000 Genomes LD reference data^26,27,^ which may be due to convergence issue in the presence of inadequate LD reference sample size and/or ancestry mismatch between 1000 Genomes samples and the target population^26^. Implemented by an MCMC algorithm that utilizes similar computational tricks as LDpred2, MUSSEL may likewise underperform with the 1000 Genomes reference data. We therefore conduct a sensitivity analysis where we estimate LD based on UKBB tuning samples (10,000 EUR, 4,585 AFR, 687 AMR, 1,010 EAS, 5,427 SAS) instead of the 1000 Genomes samples. We observe that the R^2^ of MUSSEL PRS improves notably compared to using 1000 Genomes LD reference (Figure 4(b), Supplementary Table 4), especially on AFR (average increase: 33.8%). The R^2^ of PRS-CSx PRS has also increased but not as much as the R^2^ of MUSSEL PRS. This is particularly noteworthy because PRS-CSx by default uses a much larger number of UKBB LD reference samples (375,120 EUR, 7,507 AFR, 687 AMR, 2,181 EAS, and 8,412 SAS), which also overlap with our UKBB testing samples and thus lead to potentially inflated R^2^ estimates. The advantage of MUSSEL now becomes more obvious: it outperforms the existing methods in all scenarios except for HDL in EAS, where it performs slightly worse than PRS-CSx PRS. MUSSEL shows the most notable advantage on AFR, for which PRS are typically not powerful and hard to improve (average R^2^ increase compared to the best existing method: 38.6%). Interestingly, the alternative weighted MUSS approach has a similar or slightly lower R^2^ than MUSSEL, but it still outperforms PRS-CSx, which utilizes the same linear combination strategy, for almost all traits and ancestry groups (Supplementary Figure 14(b)).

### AoU data analysis with validation on UKBB individuals

We also apply the various methods to develop ancestry-specific PRS for height and BMI based on the GWAS summary data we generated from the All of Us Research Program (AoU) for EUR, AFR, and AMR. The performance of the derived PRS is evaluated on UKBB validation samples of AFR ancestry. As in the GLGC data analysis, we first use genotype data of the unrelated 1000 Genomes samples as the LD reference data^24^ (Figure 5(a), Supplementary Table 5). Although no method is uniformly the best on all traits and ancestry groups, MUSSEL PRS on average has an R^2^ that is 67.5% higher than that of the PRS-CSx PRS and 53.4% higher than that of the CT-SLEB PRS. MUSSEL PRS improves on top of the single-ancestry method by both the Bayesian modeling step (MUSS versus LDpred2, Figure 5(a)) and the SL step (MUSSEL versus MUSS, Supplementary Figure 15(a)). The weighted MUSS PRS utilizing a linear combination strategy gives a lower R^2^ than the MUSSEL PRS utilizing the EL strategy (weighted MUSS versus MUSSEL, Supplementary Figure 15(a)).

Similar to the GLGC data analysis, we also conduct a sensitivity analysis where we estimate LD using the UKBB tuning samples (10,000 EUR, 4,585 AFR, 1,010 EAS, 5,427 SAS) instead of the 1000 Genomes data. Different from the results from GLGC data analysis, no PRS has noticeably improved predictive power, even though there is a better ancestry match between the LD reference population and the target population (Figure 5(b), Supplementary Figure 15(b)). Such results from the GLGC data analysis and the AoU data analysis suggest that for MUSSEL, 1000 Genomes LD reference dataset may be adequate for building PRS models with relatively small discovery GWAS, such as the AoU GWAS (N = 15,364 – 48,332), but not so with much larger discovery GWAS, such as the GLGC GWAS (N up to 0.89 million). In other words, the ratio of the sample size of the LD reference dataset to the GWAS sample size may matter more than the sample size of the LD reference data itself or the population/ancestry match between datasets.

### 23andMe data analysis

We have collaborated with 23andMe, Inc. to develop and validate PRS for seven traits for EUR, African American (AFR), Latino (AMR), EAS, and SAS based on a large-scale dataset from 23andMe, Inc. We analyze two continuous traits: (1) heart metabolic disease burden, (2) height, and five binary traits: (3) any cardiovascular disease (any CVD), (4) depression, (5) migraine diagnosis, (6) morning person, and (7) sing back musical note (SBMN). Results are summarized in Figure 6 and Supplementary Table 6. For the two continuous traits, MUSSEL shows a major advantage over the existing methods on AFR and AMR: for example, MUSSEL has a remarkable improvement over two recently proposed advanced methods that perform the best among the existing methods, PRS-CSx (average increase in R^2^: 49.8%) and CT-SLEB (average increase in R^2^: 47.5%). For EAS and SAS, MUSSEL performs better than all existing methods considered in all scenarios, except for heart metabolic disease burden in SAS, which has the smallest discovery GWAS (N = 20,062), where MUSSEL PRS has an R^2^ slightly lower than that of CT-SLEB PRS but higher than the R^2^ of all other PRS.

**Figure 6:**
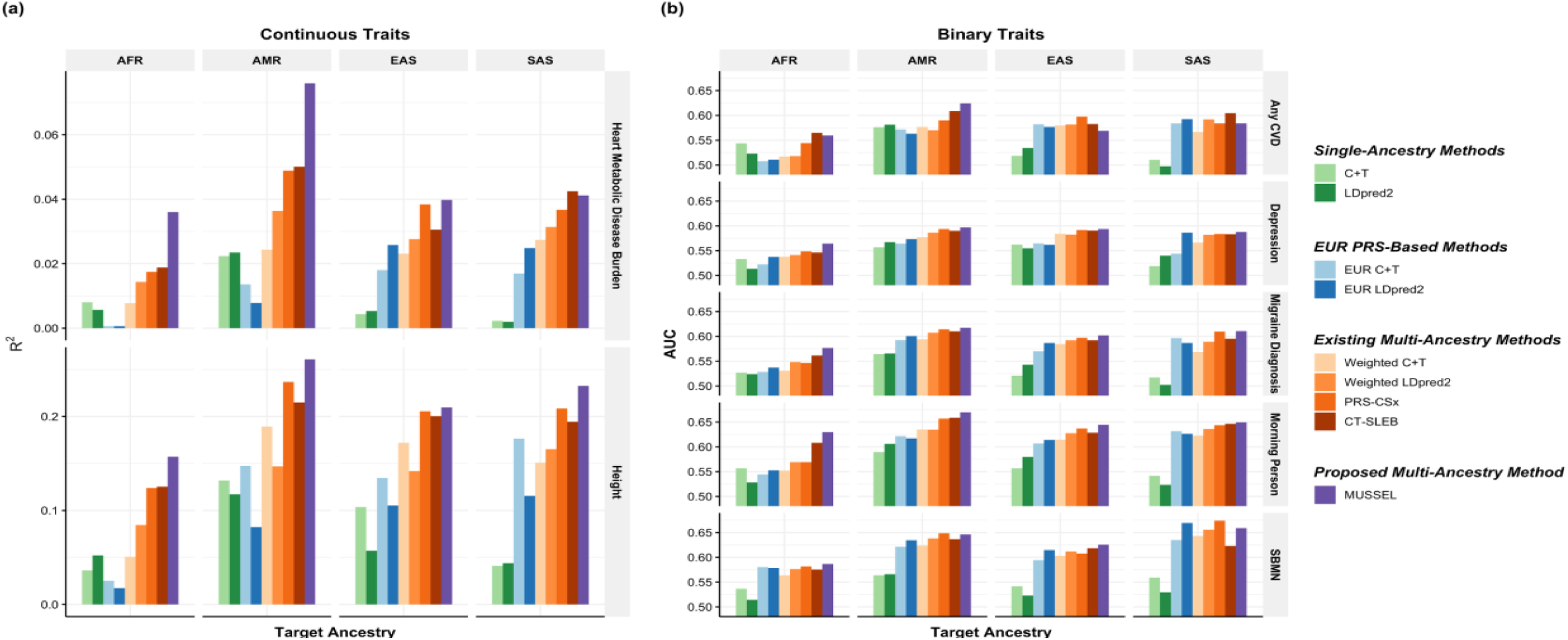
Prediction results on 23andMe validation individuals based on discovery GWAS from 23andMe on EUR, African American (AFR), Latino (AMR), EAS, and SAS. The performance of the various methods is evaluated by **(a)** residual R^2^ for two continuous traits, heart Metabolic Disease Burden and height, and **(b)** residual AUC for five binary traits, any CVD, depression, migraine diagnosis, morning person, and SBMN. The LD reference data is from the 1000 Genomes Project (498 EUR, 659 AFR, 347 AMR, 503 EAS, 487 SAS). The dataset is randomly split into 70%, 20%, 10% for training GWAS, model tuning (tuning model parameters and training the SL in CT-SLEB and MUSSEL or the linear combination model in weighted LDpred2 and PRS-CSx), and testing (to report residual R^2^ or AUC values after adjusting for the top 5 genetic principal components, sex, and age), respectively. All methods were evaluated on the ∼2.0 million SNPs that are available in HapMap3 + MEGA, except for PRS-CSx which is evaluated based on the HapMap 3 SNPs only, as implemented in their software. Ancestry- and trait-specific sample sizes of GWAS, number of SNPs included, and validation sample sizes are summarized in Supplementary Table 6.1.

For the five binary traits, we observe a similar pattern as for continuous traits, where MUSSEL generally performs better than or similarly to the best of the existing methods, and it shows the biggest improvement in (AUC – 0.5) over existing methods on AFR (average improvement: 14.4%, Figure 6(b), Supplementary Table 6). Averaged across all five traits and four non-EUR ancestry groups, MUSSEL PRS gives an (AUC – 0.5) that is 13.8% higher than that of the PRS-CSx PRS and 9.0% higher than that of the CT-SLEB PRS.

## Discussion

We propose MUSSEL, a powerful method for constructing enhanced ancestry-specific PRS integrating information from GWAS summary statistics and LD reference data across multiple ancestry groups. Built based on an extension of spike-and-slab type prior^14^, MUSSEL enhances the ancestry-specific polygenic prediction by (1) borrowing information from GWAS of other ancestries via specification of a between-ancestry covariance structure in SNP effect sizes, (2) incorporating heterogeneity in LD and MAF distribution across ancestries, and (3) an SL algorithm combining ancestry-specific PRS developed under various possible genetic architectures of the trait. We benchmark our method against a wide variety of alternatives, including multiple state-of-the-art multi-ancestry methods^8,9^,13, using extensive simulation studies and data analyses. Results show that while no method is uniformly the best, MUSSEL is generally a robust method that shows close to optimal performance across a wide range of scenarios and have the potential to notably improve PRS performance in the AFR population compared to the alternative methods.

One important observation from the data applications is that the advantage of MUSSEL over existing methods tends to be more notable with larger GWAS accompanied by larger LD reference dataset. In the GLGC and 23andMe data analyses where the discovery GWAS sample sizes are relatively large, especially for the non-EUR populations, we can clearly observe that MUSSEL performs almost uniformly better than the existing methods. In contrast, in the PAGE + UKBB + BBJ data analysis, where the GWAS sample sizes for AFR and AMR are relatively small, MUSSEL sometimes shows a suboptimal performance. Such trend of having more notable advantages with larger GWAS sample sizes and larger LD reference datasets exists not only when comparing MUSSEL to existing methods, but also when comparing the more advanced methods, such as MUSSEL and PRS-CSx, to simpler alternatives, such as the weighted PRS method.

One key factor in implementing MUSSEL is the LD reference data. The analyses of the GLGC and AoU datasets illustrates that the sample size of the LD reference data should be sufficiently large relative to the discovery GWAS sample size to give MUSSEL an optimal performance (Figure 4, Supplementary Table 4). The performance of MUSSEL depends on estimated causal SNP proportion parameters from single-ancestry LDpred2 analysis. LDpred2 has previously been shown to underperform sometimes when using 1000G LD reference data^27^ and thus could in turn affect the performance of MUSSEL. Thus, as sample sizes of the training GWAS increase, building a larger LD reference dataset than the widely used 1000 Genomes reference dataset will lead to more optimal performance.

The performance of MUSSEL is robust to modest ancestry mismatch between the discovery GWAS samples, LD reference samples, and validation samples, such as EUR in the US versus EUR in the UK, as shown in the All of Us data analysis. In our simulation study, we conducted a sensitivity analysis on the performance of MUSSEL given 20% ancestry mismatch between the discovery GWAS samples and LD reference samples. While the power loss of MUSSEL, as well as the LDpred-based methods, is within a reasonable range, an interesting finding is that the amount of power loss appears to increase as the underlying causal SNP proportion decreases. This suggests that, for MUSSEL and the LDpred-based methods, ancestry mismatch between samples may be a more severe issue for those traits that are affected by a small number of large-effect SNPs. Ideally, the populations should be matched as closely as possible between GWAS samples, LD reference samples, and validation samples to ensure an optimal performance of MUSSEL. But if there is slight LD misspecification, e.g., using samples from White population in the UK to estimate LD among White population in the US, our analyses on the simulated and real datasets suggest that the power of MUSSEL may be slightly worse but still comparable.

There are several practical considerations regarding the implementation of MUSSEL and other multi-ancestry methods. First, the SL step in MUSSEL and CT-SLEB needs to be implemented with caution. We have shown by our data examples that the SL algorithm combining PRS models across various tuning parameter settings could yield additional improvement in predictive power. With a limited tuning sample, however, the SL might be overfit in the presence of a large number of tuning parameter settings, ultimately leading to low predictive power in an independent sample. Our analysis on the simulated data suggests that the performance of SL combining 30 different PRS models is typically stable when the effective sample size of the tuning dataset is no less than 1000 for continuous traits. The required tuning sample size will also increase as the number of PRS models included in SL increases. Second, the advanced multi-ancestry methods, such as PRS-CSx, CT-SLEB and MUSSEL, may not yield higher predictive power if the training GWAS sample size is too small. We expect the advanced multi-ancestry methods to outperform simpler methods when GWAS sample size is relatively large (e.g., over 15,000 per ancestry group as in the AoU data analysis). When discovery GWAS for the target non-EUR ancestry group is relatively small (several thousand samples or fewer), then a single-ancestry PRS model trained based on the much larger EUR GWAS may outperform the multi-ancestry methods.

We have compared MUSSEL with a series of recent multi-ancestry methods including PRS-CSx and CT-SLEB, but there are other recently proposed methods that are worth investigating. In fact, we have implemented two other multi-ancestry methods named XPASS and PolyPred+ in our simulation study as well as GLGC, AoU, and 23andMe data analyses, with detailed results reported in Zhang et al. (2022)^13^. Although computationally super-fast, XPASS, which uses a bivariate normal prior under an infinitesimal model, can only combine up to two ancestry groups, and it is always outperformed by MUSSEL (Supplementary Tables 3-6). This shows the importance of including sparsity components in modeling effect-size distribution for Bayesian polygenic prediction. PolyPred+ implements a linear combination of SBayesR^28^ trained separately on EUR and the target population and a PolyFun^29^ PRS on EUR that additionally incorporates information from external functional annotations, and thus it is not directly comparable to the other methods. Even so, it performs worse than MUSSEL most of the time (Supplementary Tables 3-6).

Our study also has several limitations. First, the MUSS step requires two sets of tuning parameters: causal SNP proportion in each ancestry and between-ancestry correlation in effect sizes, the specification of which is relatively complex compared to other methods such as PRS-CSx. In the default setting of MUSS, the candidate values for genetic correlation between a pair of ancestry groups only lie between 0.7 and 0.95, while for some traits, the estimated correlation can be lower^9,25.^ But given the high computational scalability of MUSS, when the number of ancestry groups is not too large (*K* ≤ 5), prior information on genetic correlation can used to specify additional genetic correlation parameter settings to cover a wider range of potential genetic architectures of the trait. Second, all our analyses are based on a set of *approximately 2*.*0 million SNPs selected based on the combined content of the HapMap 3 and the MEGA SNP array. While this SNP set is considered very informative for multi-ancestry genetic studies*, we have previously shown that it is possible to increase PRS performance, especially in the AFR populations, by including much larger SNP contents. Future research is needed to improve scalability of the methods like PRS-CSx and MUSSEL to datasets with larger SNP contents.

The spike-and-slab type prior in MUSSEL can be sub-optimal for effect-size distribution of some traits. For example, in GLGC GWAS, we detect several top SNPs with extremely large association coefficients for all four blood lipid traits, each contributing to 0.6% -3.9% of the estimated total heritability. In this case, the Bayesian step in MUSSEL induces the same amount of shrinking on all SNPs, resulting in over-shrinkage on the few large-effect SNPs. We have considered a simple alternative approach to compensate such over-shrinkage^30,31^, where for each target ancestry group, we first construct a “top-SNP PRS” using GWAS association coefficients of the few top SNPs for the ancestry, then combine it with the MUSSEL PRS constructed based on the rest of the SNPs. This approach, however, does not provide a more powerful PRS. PRS-CSx, which allows a heavy-tail Strawderman-Berger prior, while theoretically expected to be advantageous for handling such large-effect SNPs, does not show much advantage either. In the future, other heavy-tail type priors such as the Bayesian Lasso (i.e., Laplacian)^32^, Horseshoe^33^, and Bayesian Bridge^34^, are worth investigating. Another potential limitation of the method originates in the SL step: when the tuning sample is small (e.g., <1000), the prediction algorithms utilized in SL may be overfit in the presence of a large number of tuning parameters, ultimately leading to low predictive power in an independent sample.

In our data examples, different methods show advantages in different scenarios in terms of GWAS sample size, LD reference data, the type of trait, and target ancestry. It is thus natural to consider extending our EL step from combining a series of PRS trained within a specific type of method, such as MUSS, to those generated across different methods. MUSSEL can also be modified to enhance performance of PRS by borrowing information simultaneously across traits and genetically correlated traits. Two recent studies, both using simple weighting methods, have shown significant potential for cross-trait borrowing to improve PRS performance for individual traits^35,36^. There is, however, likely to be scope for additional improvement by developing formal Bayesian methods that can utilize flexible models for effect-size distribution simultaneously across ancestries and traits.

In summary, we propose a powerful method for constructing enhanced ancestry-specific PRS combining GWAS summary data and LD reference data across multiple ancestry groups. As sample sizes of the multi-ancestry GWAS and LD reference datasets continue to increase, more advanced methods, such as MUSSEL and PRS-CSx^9^, are expected to show more and more advantages over simpler alternatives, such as the weighted methods^8^. Our large-scale simulation study and four unique data examples illustrate the relative performance of a variety of single- and multi-ancestry methods across various settings of ancestry groups, GWAS sample sizes, genetic architecture of the trait, and LD reference panel, which can serve as a guidance for method implementation in future applications.

## Competing interests

### Data and code availability

The simulated genotype data for 600K subjects of EUR, AFR, AMR, EAS, or SAS ancestry can be accessed at https://dataverse.harvard.edu/dataset.xhtml?persistentId=doi:10.7910/DVN/COXHAP. The EUR

GWAS summary data for BMI^37^, HDL^38^, and LDL^38^ based on UKBB samples (GWAS round 2) published by the Neale Laboratory can be downloaded at http://www.nealelab.is/uk-biobank. The EAS GWAS summary data from BBJ for BMI^39^, HDL^40^, and LDL^40^ were downloaded from http://jenger.riken.jp/en/result. Split GWAS summary data from PAGE for BMI, HDL, and LDL stratified for AFR and AMR, as used in the training sets in our data analysis, are available upon request (email to). Stratified GWAS summary data from PAGE for BMI, HDL and LDL for AFR and AMR (not split for training/validation sets) is available on LDHub (https://ldsc.broadinstitute.org). GWAS summary data from GLGC for HDL, LDL, TC, and logTG stratified for EUR, AFR, AMR, EAS, and SAS can be downloaded at http://csg.sph.umich.edu/willer/public/glgc-lipids2021/results/ancestry_specific/. GWAS summary data from AoU for BMI and height stratified for EUR, AFR, and AMR are available upon request (email to). GWAS summary data from 23andMe Inc. for top 10,000 genetic markers associated with height, morning person, and SBMN across five ancestry groups has been made available at https://dataverse.harvard.edu/dataset.xhtml?persistentId=doi:10.7910/DVN/3NBNCV. The full GWAS summary statistics for these three traits (height, morning person, and SBMN) are available through 23andMe to qualified researchers under an agreement with 23andMe Inc. that protects the privacy of the 23andMe participants. Please visit https://research.23andme.com/collaborate/#dataset-access/ for more information and to request data access. GWAS summary statistics for the other four traits (any CVD, heart metabolic disease burden, depression, and migraine) will not be made available because of 23andMe business requirements. Participants included in our 23andMe data analysis provided informed consent and participated in the research online, under a protocol approved by the external AAHRPP-accredited IRB, Ethical & Independent Review Services. 1000 Genomes Phase 3 reference data can be downloaded from https://mathgen.stats.ox.ac.uk/impute/1000GP_Phase3.html. Our estimated LD block matrices for EUR, AFR, AMR, EAS, and SAS for approximately 2.0 million SNPs in HapMap 3 plus MEGA that are also available in 1000 Genomes Project can be downloaded from https://github.com/Jin93/MUSSEL. LD block information, including the start and end positions of each block, are extracted from the “lassosum” R package and can be downloaded from https://github.com/tshmak/lassosum.

PLINK 1.9: https://www.cog-genomics.org/plink. PLINK 2.0: https://www.coggenomics.org/plink/2.0/. LDpred2: https://privefl.github.io/bigsnpr/articles/LDpred2.html. The R package “bigsnpr” used in the LDpred2 pipeline is available for download on Github at https://github.com/privefl/bigsnpr. PRS-CSx: https://github.com/getian107/PRScsx. CT-SLEB: https://github.com/andrewhaoyu/CTSLEB. LD score regression: https://github.com/bulik/ldsc. The MUSSEL pipeline, along with the R code for simulation studies and data analyses in this paper can be accessed at https://github.com/Jin93/MUSSEL.

## Acknowledgements

This work was supported by the following NIH grants: R00 HG012223 (J.J.), K99 CA256513 (H.Z.), R01 HG010480 (N.C., J.J., and J.Zhang), U01HG011724 (N.C.), R35 HG011944 (G.L.W.), and U01 HG007419 (G.L.W.). We thank the Neale Lab and BBJ for making the GWAS summary data from UKBB and BBJ publicly available. Individual-level genotype and phenotype data for UKBB validation samples were obtained under application 17731. The PAGE Study is supported by the following NIH grants: U01 HG007419, R01 HG010297, and R01 HL151152. The All of Us Research Program is supported by the National Institutes of Health, Office of the Director: Regional Medical Centers: 1 OT2 OD026549; 1 OT2 OD026554; 1 OT2 OD026557; 1 OT2 OD026556; 1 OT2 OD026550; 1 OT2 OD 026552; 1 OT2 OD026553; 1 OT2 OD026548; 1 OT2 OD026551; 1 OT2 OD026555; IAA #: AOD 16037; Federally Qualified Health Centers: HHSN 263201600085U; Data and Research Center: 5 U2C OD023196; Biobank: 1 U24 OD023121; The Participant Center: U24 OD023176; Participant Technology Systems Center: 1 U24 OD023163; Communications and Engagement: 3 OT2 OD023205; 3 OT2 OD023206; and Community Partners: 1 OT2 OD025277; 3 OT2 OD025315; 1 OT2 OD025337; 1 OT2 OD025276. In addition, the AoU Research Program would not be possible without the partnership of its participants. We would like to thank the research participants and employees of 23andMe, Inc. for making this work possible. We would like to thank Liz Noblin, Melissa J. Francis, and Emily Voeglein for helping with the research collaboration agreement with Harvard T.H. Chan School of Public Health, Johns Hopkins Bloomberg School of Public Health and 23andMe, Inc. We would like to thank the research participants and employees of 23andMe for making this work possible. The analyses in this paper utilized the high-performance computation Biowulf cluster at National Institutes of Health, USA, Faculty of Arts and Sciences Research Computing Cluster at Harvard University, and the Joint High Performance Computing Exchange at Johns Hopkins Bloomberg School of Public Health.

## Author Contributions

J.J. and N.C. developed all methods; J.J., J.Zhang and H.Z. conducted data analyses under the supervision of N.C; H.Z. created the simulated datasets and ran GWAS on the simulated training data with the supervision from N.C.; G.L.W. ran GWAS on the training data from the PAGE consortium; R.Z. ran GWAS on the training data from AoU under the supervision of N.C.; J.Zhan., J.O.C. and Y.J. run GWAS for training data from 23andMe, Inc. under the supervision of B.L.K; J.J. developed the software; J.J. and N.C. drafted the manuscript, and H.Z., J.Zhang, and G.L.W. provided comments; all co-authors reviewed and approved the final version of the manuscript. The following members of the 23andMe Research Team have contributed to this study: Stella Aslibekyan, Adam Auton, Elizabeth Babalola, Robert K. Bell, Jessica Bielenberg, Katarzyna Bryc, Emily Bullis, Daniella Coker, Gabriel Cuellar Partida, Devika Dhamija, Sayantan Das, Sarah L. Elson, Nicholas Eriksson, Teresa Filshtein, Alison Fitch, Kipper Fletez-Brant, Pierre Fontanillas, Will Freyman, Julie M. Granka, Karl Heilbron, Alejandro Hernandez, Barry Hicks, David A. Hinds, Ethan M. Jewett, Yunxuan Jiang, Katelyn Kukar, Alan Kwong, Keng-Han Lin, Bianca A. Llamas, Maya Lowe, Jey C. McCreight, Matthew H. McIntyre, Steven J. Micheletti, Meghan E. Moreno, Priyanka Nandakumar, Dominique T. Nguyen, Elizabeth S. Noblin, Jared O’Connell, Aaron A. Petrakovitz, G. David Poznik, Alexandra Reynoso, Morgan Schumacher, Anjali J. Shastri, Janie F. Shelton, Jingchunzi Shi, Suyash Shringarpure, Qiaojuan Jane Su, Susana A. Tat, Christophe Toukam Tchakouté, Vinh Tran, Joyce Y. Tung, Xin Wang, Wei Wang, Catherine H. Weldon, Peter Wilton, Corinna D. Wong.

## Declaration of interests

J.Zhan, Y.J., J.O., and B.L.K. are employed by and hold stock or stock options in 23andMe, Inc.

## Online Methods

### Details of MUSSEL Step 1: MUSS

MUSS conducts Bayesian modeling to generate ancestry-specific MUSS PRS models through joint modeling of GWAS summary data across all available ancestry groups. This step models the genetic correlation structure in SNP effect size across ancestry groups while accounting for ancestry-specific LD and allele frequency information.

Suppose we are interested in predicting the polygenic risk of some trait *Y* based on genotype {*G*_*j*_, *j* = 1, …, *M*_*k*_}, for an individual of ancestry *k* = 1,2, …, *K*, with *M*_*k*_ denoting the number of SNPs with a minor allele frequency (MAF) > 0.01 in ancestry *k*. For demonstration purposes, we assume the trait is continuous, but the results can be directly applied to GWAS summary-level association statistics for discrete traits in the same manner. We assume all SNPs included are biallelic, i.e., each SNP only has two alleles observed in the population. For each ancestry group *k*, we assume a true additive model for genetic variation, 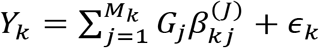, where 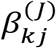 denotes the underlying joint effect size of *G*_*j*_, *j* = 1,2, …, *M*_*k*_, i.e., effect size after adjusting for the effect of other SNPs, for an individual of ancestry *k*, and ϵ_*k*_ denotes a zero-mean random error term that includes effects of risk factors other than SNPs. Suppose we have ancestry-specific GWAS summary data, 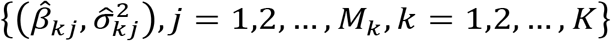, specifically, the marginal effect sizes of the SNPs 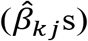 and their corresponding standard errors 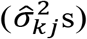 from one-SNP-at-a-time regressions, *y*_*ki*_ = *G*_*ji*_*β*_*kj*_ + *v*_*ki*_, *i* = 1, …, *N*_*k*_, for *j* = 1, …, *M*_*k*_ and *k* = 1, …, *K*. Here, *i, j* and *k* are the indices of GWAS sample, SNP, and ancestry, respectively, *v*_*ki*_ denotes a zero-mean random error term that includes effects of other risk factors and all other SNPs, *β*_*kj*_, *M*_*k*_ and *N*_*k*_ are the true marginal SNP effect sizes, total number of SNPs, and GWAS sample size, respectively, for ancestry *k*. Our goal is to obtain an estimate of the joint SNP effect sizes, 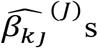, to construct polygenic risk model 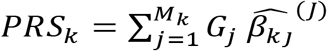 for each ancestry group *k*.

Our analysis is conducted on the standardized scale, where *G*_*kj*_s are assumed to be standardized to have a zero mean and unit variance and *Y*_*k*_ s are assumed to have a unit variance (for continuous traits). This is reflected by rescaling the GWAS summary statistics so that the variance is equal to the inverse of the GWAS sample size. For computational scalability, we divide the whole genome into a series of independent LD blocks^41^, each containing hundreds of (up to ∼2900) SNPs, and only consider the between-SNP correlation within each LD block. Such block structure for LD matrices is considered because it yields similar predictive power as the banded-structure LD matrices accounting for LD within a 3cM genetic distance suggested by LDpred2^14^, but it is computationally more efficient and requires less memory. We estimate LD matrices for SNPs within each LD block using PLINK 2.0^42^ based on LD block segmentation in Berisa and Pickrell (2015)^41^. LD block information was extracted from the R package “lassosum”^43^. Note that the LD block information is available for EUR (1747 blocks, median number of SNPs per block: 816), AFR (2626 blocks, median number of SNPs per block: 716), and EAS (1489 blocks, median number of SNPs per block: 815), but not currently available for AMR and SAS, and thus we apply the EUR LD information on AMR and SAS for now.

We denote by 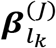 and 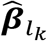 the vector of true joint effect sizes and marginal effect sizes estimated from GWAS, respectively, for SNPs within a specific LD block *l*_*k*_ in ancestry *k* = 1,2, …, *K*. To conduct analyses on the standardized scale, we first divide each raw effect size estimate 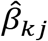 by 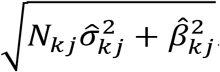. We can then write down the likelihood of the GWAS summary statistics, 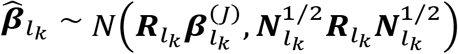, where 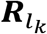 denotes the LD matrix of the SNPs within the LD block *l*_*k*_, and 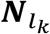 is a diagonal matrix with diagonal entries being the corresponding GWAS sample sizes for SNPs within the LD block. For population-specific SNPs, i.e., SNPs with an MAF > 0.01 in only one ancestry *k*, we assume a spike-and-slab prior as in LDpred2, 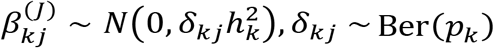, where 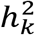 denotes the per-SNP heritability, *δ*_*kj*_ is the indicator of whether SNP *j* is causal in ancestry *k*, i.e., 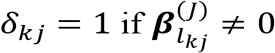 and 0 otherwise, and *p*_*k*_ is the proportion of causal SNPs in ancestry *k*. For SNPs that have MAF>0.01 in all ancestry groups, we induce a prior correlation structure between 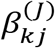 and 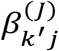 for *k, k*^′^ ∈ {1,2, …, *K*}. The prior distribution of the joint effect size 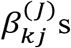 given *δ*_*kj*_ s is then specified as follows,

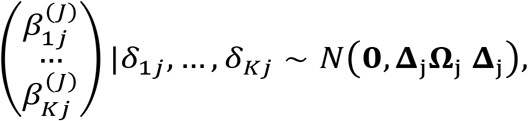

where Δ_j_ = diag(*δ*_1*j*_, …, *δ*_*Kj*_), and 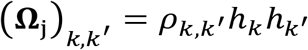, with *ρ*_*k,k*′_ denoting the genetic correlation between ancestry groups *k* and *k*^′^. For SNPs that have an MAF>0.01 in only a subset of ancestries *A* ⊂ {1, …, *K*}, similar prior distributions can be specified for SNP effect sizes within the set of ancestry groups *A*.

Recall that we introduce variables {*p*_*k*_ = Pr(*δ*_*kj*_ = 1), ∀*j, k* = 1, …, *K*} to denote ancestry-specific causal SNP proportions, and for ancestry-specific SNPs, we assume *δ*_*kj*_ ∼ Ber(*p*_*k*_). Now we generalize this Bernoulli prior to a multinomial prior on (*δ*_1*j*_, …, *δ*_*Kj*_) ^*T*^ for SNPs that exist in a subset of ancestry groups *A* ⊂ {1, …, *K*}, with probabilities 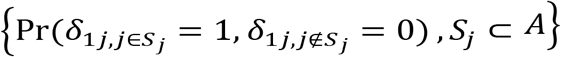 being defined as functions of *p*_*k*_, *k* = 1, …, *K*. We first focus on SNPs that only exist in two ancestry groups *A* = {*k*_1_, *k*_2_}: we set 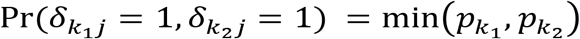, which reflects our assumption that if a SNP is causal in one ancestry group, it is also causal in another. We can then obtain 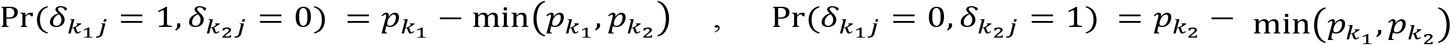, and 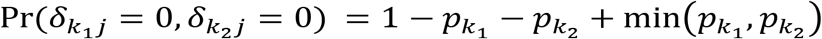. After constructing 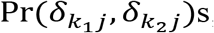, we then construct priors for SNPs that exist in three ancestry groups: by specifying 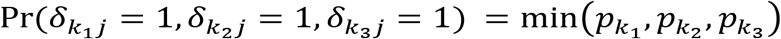, we can obtain the rest of the probabilities 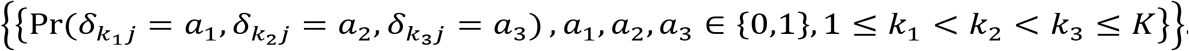. Such specifications can be easily extended to apply to SNPs that exist in four ancestry groups, five ancestry groups, etc.

We estimate 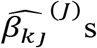 based on MCMC with an approximation strategy previously implemented in the LDpred2 algorithm^14^, which substantially reduces computation time of the algorithm. There are two sets of tuning parameters which will be estimated by grid search using a tuning dataset independent from the testing samples on which we report R^2^: (1) the ancestry-specific causal SNP proportions (*p*_1_, …, *p*_*K*_): we fix (*p*_1_, …, *p*_*K*_) to either 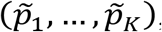, the estimated ancestry-specific causal SNP proportions obtained from LDpred2 separately on GWAS summary data of each ancestry, or 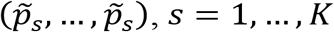, *s* = 1, …, *K*, i.e., the values of all *p*_*k*_s are set to the LDpred2 estimate of the causal SNP proportion in ancestry *s*; (2) the between-ancestry correlation parameters *ρ*_*kk*′_s: we consider two settings, i.e., either set *ρ*_*kk*’_s to all equal to *ρ* = 0.7, 0.8, 0.9, or 0.95, or set *ρ*_*kk*’_ to 0.75 for any pair of ancestry groups that include AFR and 0.9 otherwise, given that correlation with AFR tends to be weaker than that among other ancestry groups. Prior to the implementation of MCMC, we further estimate the ancestry-specific heritability 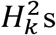 based on GWAS summary data and LD reference data using LD score regression^44^.

We now describe the detailed MCMC algorithm and estimation procedure. For SNPs that only exist (MAF>0.01) in one ancestry group, the Gibbs sampler in Vilhj álmsson et al. (2015)^45^ was implemented. For each SNP *j* that exists in all *K* ancestry groups, we sample ***δ***_*j*_ = (*δ*_1*j*_, …, *δ*_*Kj*_) ^*T*^ and ***β***_*j*_ = (*β*_1*j*_, …, *β*_*Kj*_) ^*T*^ from

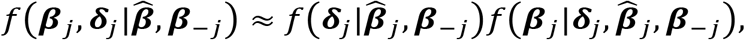

where ***β***_−*j*_ denotes the joint effect sizes for the SNPs within the LD block which SNP *j* is in, *l*_*kj*_, *k* ∈ {1, …, *K*}.

We first sample ***δ***_*j*_ from 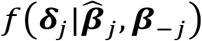. Here note that obtaining 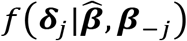 analytically is hard, and thus we approximate it by 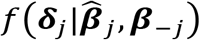. For a realization of ***δ***_*j*_, ***r*** = (*r*_1_, …, *r*_*K*_)^*T*^ where *r*_*k*_ ∈ {0,1}, ∀*k*, we first derive

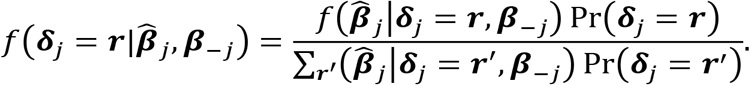

We denote the numerator by 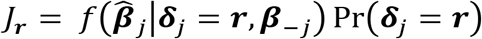), which can be derived as follows:

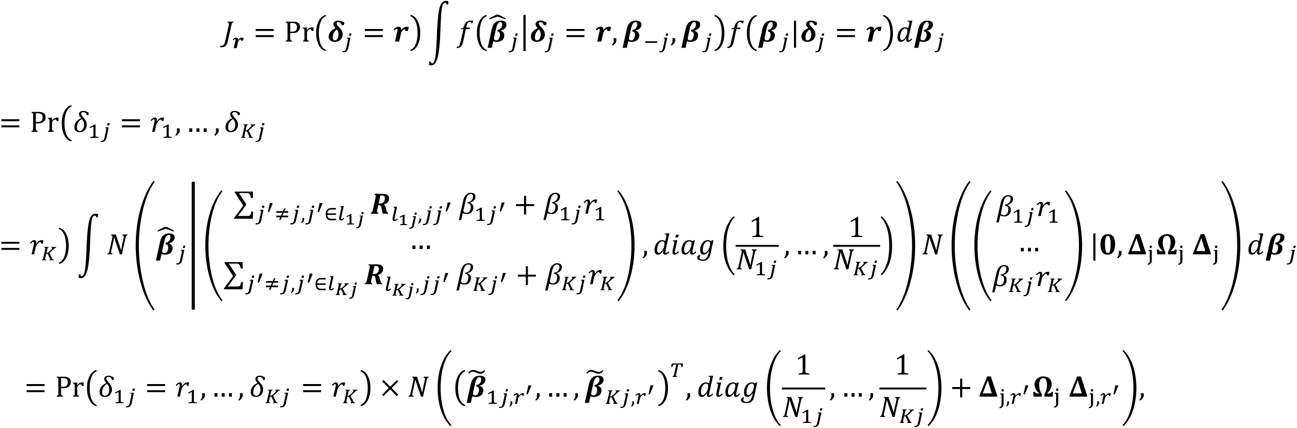

where

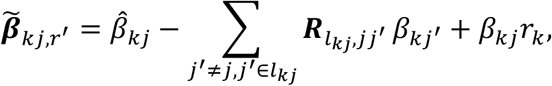

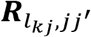 denotes the entry in 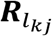 that corresponds to the correlation between SNPs *j* and *j*^′^, *I*_*a*_ = 1 if *a* ≠ 0 and 0 otherwise, 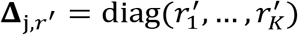, and 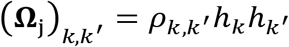. After deriving *J*_*r*_ s, we can then sample from 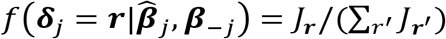.

We obtain the marginal posterior mean of ***β***_*j*_ after integrating out ***δ***_*j*_:

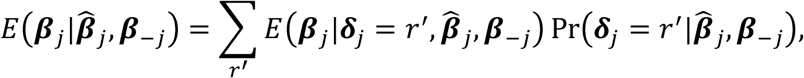

where

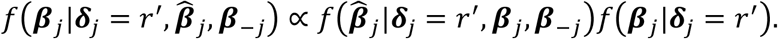

We can easily derive that 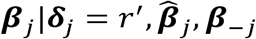 follows *N*(***μ*** _*j,r*′_, ***V***_*j,r*′_), where

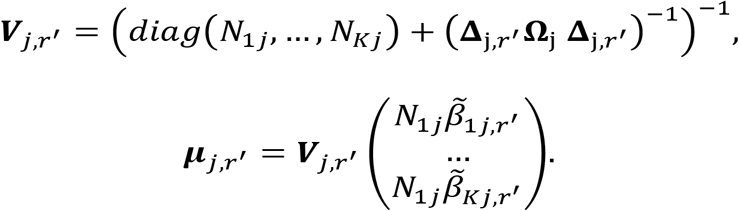

For SNPs that have an MAF>0.01 in a subset of ancestry groups *A* ⊂ {1, …, *K*}, similar sampling strategy can be conducted but only among ancestry groups *A*. In each MCMC iteration, the prior per-SNP heritability parameter is set to 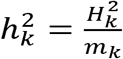, where *m*_k_ denotes the number of causal SNPs 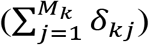 estimated from this iteration. The posterior estimate of ***β***_*j*_ is obtained by taking the average of 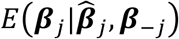) obtained from 100(*K*-1) MCMC iterations after a burn-in stage of 100 iterations.

### Existing Methods

#### Single-Ancestry Methods

##### LD Clumping and Thresholding (C+T)

C+T first constructs a series of PRS by applying an LD clumping step followed by a p-value filtering step with varying p-value cutoffs, then selects the best performing PRS on the tuning dataset. Specifically, an LD clumping step is first conducted to exclude variants that have an absolute pairwise correlation stronger than *r*^2^ =0.1 within a genetic distance (500kb) based on an LD reference dataset. The remaining variants are then filtered by excluding the ones that have a p-value larger than a significance threshold, which, in our analysis, were set to *p*_*t*_ =5 × 10^−8^, 1 ×10^−7^, 5 × 10^−7^, 1 × 10^−6^, 5 × 10^−6^, 1 × 10^−5^, 5 × 10^−5^, 1 × 10^−4^, 5 × 10^−4^, 1 × 10^−3^, 5 × 10^−3^, 1 × 10^−2^, 5 × 10^−2^, 1 × 10^−1^, 5 × 10^−1^, or 1. These 16 scores were created based on these 16 different significance thresholds *p*_*t*_s by calculating a weighted sum of the number of effect alleles of the selected SNPs, with weights being the effect size estimates from the discovery GWAS. C+T then selects the score with the “optimal” p-value thresholds via parameter tuning with respect to the residual R^2^ (for continuous traits) or residual AUC (for binary traits) on a tuning dataset that is independent of the training and testing samples. C+T was implemented using PLINK 1.90^46,47^.

##### LDpred2

LDpred2 is an LD-based Bayesian modeling approach which leverages information from GWAS summary statistics and explicitly models LD correlation structure with correlation matrices being estimated based on an external reference panel^45,48^. LDpred2 assumes a spike-and-slab prior on SNP effect sizes, i.e., each SNP has a probability *p* to have a non-zero causal effect 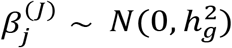, and a probability (1 − *p*) to have no contribution to the phenotypic variation 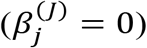. Here *p* and the total heritability, *H*^2^, are treated as tuning parameters and estimated via grid search on a tuning dataset. In each iteration of MCMC, the per-SNP heritability parameter is set to 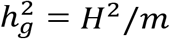, where *m* is the number of causal SNPs detected in that iteration.

We ran LDpred2 on each chromosome and GWAS of each ancestry group separately using R packages “bigstatsr” and “bigsnpr”. For our analyses on the simulated datasets, PAGE + UKBB + BBJ datasets, GLGC dataset and AoU dataset, we considered the “LDpred2 grid” model, where two tuning parameters were considered: (1) causal SNP proportion *p*, with default candidate values 1.0×10^-4^, 1.8×10^-4^, 3.2×10^-4^, 5.6×10^-4^, 1.0×10^-3^, 1.8×10^-3^, 3.2×10^-3^, 5.6×10^-3^, 1.0×10^-2^, 1.8×10^-2^, 3.2×10^-2^, 5.6×10^-2^, 1.0×10^-1^, 1.8×10^-1^, 3.2×10^-1^, 5.6×10^-1^, and 1.0; (2) total heritability *H*^2^, which is set to the heritability estimated by LD score regression^44^ multiplied by 0.7, 1, or 1.4. The “sparse” option was not considered. In our 23andMe data analysis, we considered the “LDpred2 auto” model, which estimates *p* and 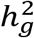 along with the other model parameters instead of treating them as tuning parameters and estimating them based on a grid search. The reason we considered the “auto” option instead of the “grid” option is that the “grid” option gave nonconvergent estimates under all considered tuning parameter settings. This convergence issue of the LDpred2 grid algorithm may be due to the low ratio between the 1000 Genomes reference sample size and the large discovery sample size of the 23andMe GWAS. We have discussed this issue in our GLGC data analysis as well.

### Multi-Ancestry Methods

#### Weighted PRS

A simple multi-ancestry method is weighted PRS, which trains an “optimal” linear combination of the effect size estimates obtained based on training data from each single ancestry. Weighted PRS was first proposed in Marquez-Luna et al. (2017)^8^ to improve the performance of single ancestry C+T PRS. Suppose we have constructed C+T PRS, *PRS*_*EUR*_, *PRS*_*AFR*_, *PRS*_*AMR*_, *PRS*_*EAS*_, and *PRS*_*SAS*_, separately based on GWAS and LD reference panel of each corresponding ancestry group. The weighted C+T PRS is then constructed as *PRS*_*wP*+*T*_ = *α*_1_ *PRS*_*EUR*_ + *α*_2_ *PRS*_*AFR*_ + *α*_3_ *PRS*_*AMR*_ + *α*_4_ *PRS*_*EAS*_ + *α*_5_ *PRS*_*SAS*_ where *α*_*k*_s are obtained by fitting a regression model on the tuning dataset. Here we apply the weighted PRS approach on either C+T (“weighted C+T”) or LDpred2 (“weighted LDpred2”).

#### PRS-CSx

“PRS-CSx”^9^ is proposed as the multi-ancestry version of PRS-CS^27^ which conducts Bayesian modeling followed by an additional step of constructing a linear combination of the best performing PRS trained for each ancestry. PRS-CSx assumes a continuous shrinkage prior named Strawderman-Berger prior on the ancestry-specific effect sizes. For SNPs available in more than one population, this prior induces information sharing across ancestry groups. After the Bayesian modeling step, PRS-CSx further trains a linear combination of the ancestry-specific PRS obtained from the previous step based on the tuning dataset. In all our analyses, we ran PRS-CSx with the default candidate values for the tuning parameter *ϕ* (1.0, 10^-2^, 10^-4^, and 10^-6^), which is the global shrinkage parameter shared by all SNPs and all ancestries that controls the overall causal SNP proportion. The PRS-CSx software only considers approximately 1.2 million HapMap 3 SNPs and therefore we only report the performance of PRS-CSx PRS based on the HapMap 3 SNPs. We have also tried to apply PRS-CSx to HapMap 3 SNPs plus an additional 0.8 million MEGA SNPs that are also available in the 1000 Genomes reference data. But we found that, on our simulated dataset, the performance of PRS-CSx PRS using the extended HapMap 3 + MEGA SNP set is significantly worse than PRS-CSx using the HapMap 3 SNPs, and in our real data analyses, results from PRS-CSx on the two SNP sets are similar. We therefore stick to the default setting with 1.2 million HapMap 3 SNPs provided by the PRS-CSx software.

#### CT-SLEB

CT-SLEB is a recently proposed method for multi-ancestry PRS construction^13^. It first conducts a two-dimensional C+T between EUR GWAS and GWAS of the target population to select SNPs to be included in the target population PRS, then uses an Empirical Bayesian approach to account for genetic correlation across populations, and finally implements an SL algorithm to combine PRS generated under different p-value thresholds in the C+T step. In our analyses, we implemented CT-SLEB with the default setting for p-value threshold, *p*_*t*_ =5 × 10^−8^, 5 × 10^−7^, 5 × 10^−6^, 5 × 10^−5^, 5 × 10^−4^, 5 × 10^−3^, 5 × 10^−2^, 5 × 10^−1^, or 1, and a genetic distance *d* =50/*r*^2^ or 100/*r*^2^, where *r*^2^ =0.01, 0.05, 0.1, 0.2, 0.5, or 0.8.

#### Runtimes and memory usage

We compare the computation time and memory usage of MUSSEL and PRS-CSx on chromosome 22 based on the simulated dataset (comparison between PRS-CSx and CT-SLEB on the same dataset has been reported in Zhang et al., 2022^13^). Results from MUSSEL and PRS-CSx combining three ancestry groups (EUR, AFR, and AMR), four ancestry groups (EUR, AFR, AMR, and EAS), and five ancestry groups (EUR, AFR, AMR, EAS, and SAS) are summarized in Supplementary Table 2. The training GWAS sample size is 15,000 for each non-EUR population and 100,000 for EUR population. The tuning and validation dataset each contains 10,000 individuals. All analyses were performed with AMD EPYC 7702 64-Core Processors running at 2.0 GHz. Other than the LDpred2 step which uses parallel computing with 17 cores, all other analyses were conducted using a single core. The reported computation time and memory usage are averaged over 10 replicates.

### PAGE + UKBB + BBJ data analysis with validation on non-EUR individuals from PAGE

Three traits, including IRNT BMI, HDL, and LDL, that were available across PAGE, UKBB, and BBJ GWAS for EUR, AFR, AMR (Hispanic), and EAS are analyzed. Ancestry- and trait-specific GWAS sample sizes, validation sample sizes, and number of SNPs analyzed are reported in Supplementary Table 3.1. The training GWAS datasets consist of PAGE, contributing data for AFR and AMR, UKBB, contributing data for EUR, and BBJ, contributing data for EAS. The validation datasets consist of PAGE, contributing data for the three non-EUR ancestry groups, and UKBB, contributing data for EUR. Specifically, we first collect data for a total of 43,769 PAGE individuals of AFR (N=17,127), AMR (N=21,995), or EAS (N=4,647) ancestry that have data available for at least one of the three traits. For AFR and AMR that have relatively large sample sizes in PAGE, we randomly divide the samples within each ancestry group into a training dataset (80%) for conducting GWAS, a tuning dataset (10%) for tuning model parameters, and training SL in CT-SLEB and MUSSEL or the linear combination model in weighted PRS and PRS-CSx, and a testing dataset (10%) for evaluating PRS performance. For EAS which has a limited sample size in PAGE, we use all PAGE samples for external validation (tuning + testing) and obtain GWAS summary data from BBJ, which has a much larger sample size. To borrow information from large EUR GWAS, we further collect EUR GWAS summary data from UKBB^49^ (N= 315,133 – 360,388) released by the Neale Lab. Finally, to tune the causal SNP proportion for EUR, which is required for specifying the prior causal probabilities for non-EUR ancestry groups, we further randomly select a sample of 20,000 random individuals from UKBB that do not overlap with samples in the EUR UKBB GWAS. Here the ancestry information for individuals from PAGE and UKBB is determined based on self-identified race/ethnicity.

For AFR and AMR, we conduct GWAS on individuals from the PAGE study to obtain the GWAS summary data. Specifically, we first collect a total of 17,127 AFR and 21,995 AMR from PAGE, then randomly divide the samples in each ancestry into a training set (80%) to conduct GWAS and a validation set (20%), of which 10% is used for selecting tuning parameters and training SL (tuning set), and the other 10% is used for reporting PRS performance (testing set). There was no significant difference between training and validation datasets in the distribution of the covariates adjusted for in GWAS. **PAGE GWAS: (1) IRNT BMI**. For ancestry-specific GWAS analysis on AFR and AMR, measurements of BMI outside of 6 standard deviations from the mean (based on sex and race) were removed. We first created sex-specific residuals for BMI adjusted for age, then inverse normally transformed these residuals. These inverse-normally-transformed residuals were then used in the final analysis where they were further adjusted for self-identified race/ethnicity, study, study center (for MEC and SOL only), and the top 10 genetic principal components (PCs). **(2) HDL**. For ancestry-specific GWAS analysis on AFR and AMR, untransformed HDL measurements were reported in mg/dL, and were adjusted for each individual’s medication use by adding a constant based on the type of medication used. Details of the adjustment are described in the Supplementary Information in Wojcik et al. (2019)^3^. Finally, models were adjusted by age at lipid measurement, sex, study, study center (for MEC and SOL only), self-identified race/ethnicity, and top 10 genetic PCs. **(3) LDL**. For ancestry-specific GWAS analysis on AFR and AMR, untransformed HDL measurements were calculated using the Friedewald Equation^50^ and reported in mg/dL. The measurements were adjusted for individuals’ medication use by adding a constant based on the type of medication used. Details of the calculation and adjustment are described in the Supplementary Information in Wojcik et al. (2019)^3^. Participants who were pregnant at blood draw or had fasted less than 8 hours prior to lipid blood draw were excluded. Finally, models were adjusted by age at lipid measurement, sex, study, study center (for MEC and SOL only), self-identified race/ethnicity, and top 10 genetic PCs.

The PAGE individuals included in our analyses are part of the PAGE participant cohort, which were collected from Hispanic Community Health Study/Study of Latinos (HCHS/SOL), Women’s Health Initiative (WHI), Multiethnic Cohort (MEC), and the Icahn School of Medicine at Mount Sinai BioMe biobank in New York City (BioMe)^3^. Due to the extensive degree of admixture within and between PAGE self-identified racial/ethnic groups, individuals were not reassigned based on their genetic ancestry but remained categorized by their self-identified race/ethnicity. However,we have assigned them to ancestry groupings based on an approximation of mappings to continental-level regions for consistency with other external studies in this manuscript. Written informed consent was obtained for all participants in this study at the relevant recruitment sites. Due to the extensive degree of admixture within and between PAGE self-identified racial/ethnic groups, individuals were not reassigned based on their genetic ancestry but remained categorized by their self-identified race/ethnicity. Detailed information about genotyping, data quality control and imputation, selection of unrelated individuals, genetic principal component analysis, and phenotype harmonization are provided in the Supplementary Information in Wojcik et al. (2019)^3^.

Since PAGE has a limited sample size for EAS (4,647), and thus we further collect publicly available GWAS summary data from BBJ (data availability) and use all PAGE individuals for validation on EAS. For BMI, the GWAS analysis included age, age^2^, sex, status of a series of diseases, and the top 10 genetic PCs as covariates^39^. For HDL and LDL, the GWAS analyses included age, sex, status of a series of diseases, and the top 10 genetic PCs as covariates^40^.

PAGE does not have individuals of EUR ancestry. To borrow information from the much larger EUR GWAS, we further download publicly available EUR GWAS summary data from UKBB (Data and code availability). For all three traits, the UKBB GWAS analyses include age, age^2^, inferred sex, an interaction term between age and inferred sex, an interaction term between age^2^ and inferred sex, and the top 20 genetic PCs as covariates. One thing to note is that for HDL and LDL, measurements are untransformed and reported in mmol/L in UKBB, untransformed and reported in mg/dL in PAGE, and reported in mg/dL then standardized to Z-score in BBJ. Although not on the same scale, the correlation in SNP effect size estimates remain the same, allowing the various GWAS summary data to be analyzed jointly. For EUR, we construct a validation dataset of 20,000 independent samples from UKBB that do not overlap with the UKBB GWAS samples. Specifically, we use the genotyping plate and well codes, which are published in the file ukb_sqc_v2.txt by UKBB and are consistent across different project applications, to identify and exclude the individuals included in the UKBB GWAS analysis by Neale Lab, and then randomly select 20,000 independent individuals from the remaining UKBB samples to conduct parameter tuning (10,000) and testing (10,000). For each ancestry group, we use unrelated samples of the same ancestry from 1000 Genomes Project as the LD reference data. For EUR, the reported prediction R^2^ are adjusted for age, sex, and top 10 genetic PCs. For AFR, AMR and EAS, the R^2^ for BMI are adjusted for age, sex, top 10 genetic PCs, and whether the individual is from the BioMe Biobank, and for HDL and LDL the R^2^ are adjusted for age at lipid measurement, sex, top 10 genetic PCs, and whether the individual is from the BioMe Biobank.

We conduct the following quality control steps for the GWAS summary-level association statistics: (1) consistent with the procedure in our simulation study and other data analyses, we restrict our analysis to approximately 1.6 million SNPs in HapMap 3 plus MEGA that are also available in LD reference panel and validation sample; (2) we remove SNPs that have duplicated positions in GWAS or LD reference panel; (3) for EUR, we remove SNPs that have alleles “AT”, “TA”, “CG”, or “GC” to avoid undetectable flipping strands when matching with UKBB validation data; (4) for the implementation of single-ancestry methods, we only keep common SNPs, i.e., SNPs that have ancestry-specific MAF > 0.01 in that ancestry group, and for the implementation of multi-ancestry methods we keep all SNPs that have ancestry-specific MAF > 0.01 in at least one ancestry group. The Manhattan plots and QQ plots for GWAS are reported in Supplementary Figures 16-18. No inflation is observed based on the genomic inflation factor. We estimate heritability of the three traits for EUR using LD score regression^44^ based on the 1000 Genomes LD reference data for EUR (Supplementary Table 7).

### GLGC data analysis with validation on UKBB individuals

We obtain GWAS summary data from the Global Lipids Genetics Consortium (GLGC) for four blood lipid traits including HDL, LDL, TC, and logTG^25^ on five ancestry groups including EUR (N_GWAS_ =840,018 – 927,975), AFR or admixed AFR (N_GWAS_ =87,759 – 92,554), Hispanic (N_GWAS_ =33,989 – 48,056), EAS (N_GWAS_ =80,676 – 145,512), and SAS (N_GWAS_ =33,658 – 34,135). Details of the study design, genotyping, quality control and GWAS are previously described^25^. We validate performance of the various methods on UKBB individuals. Specifically, we select a random set of 20,000 individuals that are of EUR origin and extracted all individuals that are of AFR (N = 9,169), EAS (N=2,019), SAS (N=10,853), or Hispanic/Latino (N=785) origin. The origin of the UKBB individuals were determined by a genetic component analysis (Supplementary Notes). We used 50% of the UKBB samples to tune model parameters and train the SL in CT-SLEB and MUSSEL or the linear combination model in weighted PRS and PRS-CS (tuning set), and the remaining 50% to evaluate PRS performance (testing set). The prediction of the genetic component has a low accuracy for AMR, and given the small number of identified AMR individuals (N=785), we do not report prediction R^2^ on UKBB AMR. We use genotype data of unrelated individuals from 1000 Genomes project or tuning samples from UKBB as the LD reference data^51^. Ancestry- and trait-specific GWAS sample sizes, validation sample sizes, and number of SNPs analyzed are reported in Supplementary Table 4.1. Based on the genomic inflation factor, no inflation is observed for the various ancestry-specific GWAS. The Manhattan plots and QQ plots are reported in Zhang et al. (2022)^13^. No inflation is observed given the genomic inflation factor. Heritability of the four traits in EUR is estimated using LD score regression (Supplementary Table 7). All GWAS summary statistics went through the same quality control steps as in PAGE + UKBB + BBJ data analysis as well as one more step, where we further remove SNPs with a GWAS sample size less than 90% of the total GWAS sample size. The GWAS summary data from GLGC does not have information on ancestry-specific MAF, and thus we use the 1000G LD reference genotype data to calculate ancestry-specific MAF for the step where we filter out all SNPs that have MAF < 0.01 in all ancestry groups. The R^2^ are adjusted for age, sex, and top 10 genetic PCs.

### AoU data analysis with validation on UKBB individuals

The individuals included in our analyses are part of the All of Us participant cohort with information collected according to the All of Us Research Program Operational Protocol (https://allofus.nih.gov/sites/default/files/aou_operational_protocol_v1.7_mar_2018.pdf). Detailed information on genotyping, ancestry determination, quality control, removal of related individuals is provided in the All Of Us Research Program Genomic Research Data Quality Report (https://www.researchallofus.org/wp-content/themes/research-hub-wordpress-theme/media/2022/06/All%20Of%20Us%20Q2%202022%20Release%20Genomic%20Quality%20Report.pdf).

On the All of Us platform, we conduct GWAS for BMI and height separately on unrelated individuals of three ancestry groups including EUR (N_GWAS_ =48,229 – 48,332), admixed AFR or AFR (N_GWAS_ =21,514 – 21,550), and Hispanic/Latino (N_GWAS_ =15,364 – 15,413). The GWAS are adjusted for age, sex, and top 16 genetic PCs. There are only about 0.9 million SNPs in HapMap 3 + MEGA that are included in our analyses, which is due to the small number of the overlapping samples across the filtered WGS data, array data, and phenotype data. Similar to the GLGC data analysis, we validate performance of the various methods on UKBB individuals, i.e., 20,000 EUR individuals and individuals of AFR (N = 9,169) origin that are identified based on a genetic component analysis (Supplementary Notes). Again, the genetic ancestry prediction accuracy for AMR is low, and considering the small number of identified AMR (N=785), we do not report validation results on UKBB AMR. We use genotype data of unrelated individuals from 1000 Genomes project or tuning samples from UKBB as the LD reference data. Ancestry- and trait-specific GWAS sample sizes, validation sample sizes, and number of SNPs analyzed are reported in Supplementary Table 5.1. Based on the genomic inflation factor, no inflation is observed other than height for Hispanic/Latino. The Manhattan plots and QQ plots are reported in Zhang et al. (2022)^13^. Heritability of the two traits in EUR was estimated using LD score regression^44^ (Supplementary Table 7). All GWAS summary statistics went through the same quality control steps as in the GLGC data analysis. The R^2^ are adjusted for age, sex, and top 10 genetic PCs.

### 23andMe Data Analysis

We develop and validate PRS for seven traits, including (1) heart metabolic disease burden, (2) height, (3) any cardiovascular disease (any CVD), (4) depression, (5) migraine diagnosis, (6) morning person, and (7) sing back musical note (SBMN) for EUR, African American (AFR), Latino (AMR), EAS, and SAS based on a large-scale dataset from 23andMe, Inc. We first conduct GWAS separately on the training dataset (70% samples) for each of the five ancestry groups, then apply the various methods to the generated GWAS summary-level association statistics and LD reference data from the 1000 Genomes Project. Within the remaining 30% of the samples, we use 20% to tune model parameters, train the SL in CT-SLEB and MUSSEL, and the linear combination model in weighted PRS and PRS-CSx, then validate the predictive performance of the constructed PRS on the remaining 10% samples. We observe from our analyses on the other three datasets that MUSSEL almost always outperforms the two alternative methods, MUSS and weighted MUSS, and thus for 23andMe data analysis, we only implement MUSSEL but not the two alternative methods.

All GWAS analyses on the training data from 23andMe, Inc. were performed adjusting for age, sex, and the top 5 genetic PCs. Genotype data of unrelated individuals from 1000 Genomes project was used to estimate LD matrices. Detailed information on participant inclusion, genotyping, phenotyping, data imputation and quality control, removing related individuals, ancestry determination, and GWAS analysis is provided in Zhang et al. (2022)^13^. Ancestry- and trait-specific GWAS sample sizes, validation (tuning + testing) sample sizes, and the number of SNPs analyzed are reported in Supplementary Table 6.1. Based on the genomic inflation factor, no inflation is observed for the various ancestry-specific GWAS. The Manhattan plots and QQ plots are reported in Zhang et al. (2022)^13^. No inflation is observed given the genomic inflation factor. Heritability of the four traits in EUR is estimated using LD score regression (Zhang et al., 2022)^13^. All GWAS summary statistics went through the same quality control steps as in PAGE + UKBB + BBJ data analysis as well as one more step where we further remove SNPs with a GWAS sample size less than 90% of the total GWAS sample size. The residual R^2^ for the two continuous traits were calculated by first regressing each trait on covariates including age, sex, and the top 5 genetic PCs, and then calculating the proportion of variation of the residual explained by the PRS. The residual AUC for the five binary traits were calculated using the “roc.binary” function in the R package RISCA version 1.0171 adjusting for the same set of covariates adjusted for the continuous traits.

## Notes

### Summary of Updates

The major updates included: (1) adding 95% bootstrap confidence intervals to the reported R2 in our simulation studies and data analyses; (2) a sensitivity analysis on the simulation dataset to show the robustness of the methods to ancestry mismatch between the discovery GWAS and LD reference dataset; and (3) we have updated the name of our method from ME-Bayes SL to MUSSEL: MUltivariate Spike and Slab and Ensemble Learning, and thus have updated the title of the manuscript to "MUSSEL: Enhanced Bayesian Polygenic Risk Prediction Leveraging Information across Multiple Ancestry Groups".

## References

1. Duncan, L. et al. Analysis of polygenic risk score usage and performance in diverse human populations. Nat Commun 10, 3328 (2019).

2. Liu, C. et al. Generalizability of Polygenic Risk Scores for Breast Cancer Among Women With European, African, and Latinx Ancestry. JAMA Network Open 4, e2119084–e2119084 (2021).

3. Wojcik, G.L. et al. Genetic analyses of diverse populations improves discovery for complex traits. Nature 570, 514–518 (2019).

4. Yu, Z. et al. Polygenic Risk Scores for Kidney Function and Their Associations with Circulating Proteome, and Incident Kidney Diseases. J Am Soc Nephrol (2021).

5. Rabinowitz, J.A. et al. Genetic propensity for risky behavior and depression and risk of lifetime suicide attempt among urban African Americans in adolescence and young adulthood. Am J Med Genet B Neuropsychiatr Genet 186, 456–468 (2021).

6. Perkins, D.O. et al. Polygenic Risk Score Contribution to Psychosis Prediction in a Target Population of Persons at Clinical High Risk. Am J Psychiatry 177, 155–163 (2020).

7. Marquez-Luna, C., Loh, P.R., South Asian Type 2 Diabetes, C., Consortium, S.T.D. & Price, A.L. Multiethnic polygenic risk scores improve risk prediction in diverse populations. Genet Epidemiol 41, 811–823 (2017).

8. Ruan, Y. et al. Improving polygenic prediction in ancestrally diverse populations. Nature Genetics 54, 573–580 (2022).

9. Cai, M. et al. A unified framework for cross-population trait prediction by leveraging the genetic correlation of polygenic traits. Am J Hum Genet 108, 632–655 (2021).

10. Tian, P. et al. Multiethnic polygenic risk prediction in diverse populations through transfer learning. Front Genet 13, 906965 (2022).

11. Sun, Q. et al. Improving polygenic risk prediction in admixed populations by explicitly modeling ancestral-specific effects via GAUDI. bioRxiv, 2022.10.06.511219 (2022).

12. Zhang, H. et al. Novel Methods for Multi-ancestry Polygenic Prediction and their Evaluations in 3.7 Million Individuals of Diverse Ancestry. bioRxiv, 2022.03.24.485519 (2022).

13. Privé, F., Arbel, J. & Vilhjálmsson, B.J. LDpred2: better, faster, stronger. Bioinformatics 36, 5424–5431 (2020).

14. Prive, F., Vilhjalmsson, B.J., Aschard, H. & Blum, M.G.B. Making the Most of Clumping and Thresholding for Polygenic Scores. Am J Hum Genet 105, 1213–1221 (2019).

15. van der Laan, M.J., Polley, E.C. & Hubbard, A.E. Super learner. Stat Appl Genet Mol Biol 6, Article25 (2007).

16. Zou, H. & Hastie, T. Regularization and Variable Selection via the Elastic Net. Journal of the Royal Statistical Society. Series B (Statistical Methodology) 67, 301–320 (2005).

17. Friedman, J., Hastie, T. & Tibshirani, R. Regularization Paths for Generalized Linear Models via Coordinate Descent. J Stat Softw 33, 1–22 (2010).

18. Weissbrod, O. et al. Leveraging fine-mapping and multipopulation training data to improve cross-population polygenic risk scores. Nat Genet 54, 450–458 (2022).

19. International HapMap, C. et al. Integrating common and rare genetic variation in diverse human populations. Nature 467, 52–8 (2010).

20. Bien, S.A. et al. Strategies for Enriching Variant Coverage in Candidate Disease Loci on a Multiethnic Genotyping Array. PLoS One 11, e0167758 (2016).

21. Siva, N. 1000 Genomes project. Nature Biotechnology 26, 256–256 (2008).

22. Graham, S.E. et al. The power of genetic diversity in genome-wide association studies of lipids. Nature 600, 675–679 (2021).

23. Privé, F., Arbel, J., Aschard, H. & Vilhjálmsson, B.J. Identifying and correcting for misspecifications in GWAS summary statistics and polygenic scores. Human Genetics and Genomics Advances 3, 100136 (2022).

24. Ge, T., Chen, C.Y., Ni, Y., Feng, Y.A. & Smoller, J.W. Polygenic prediction via Bayesian regression and continuous shrinkage priors. Nat Commun 10, 1776 (2019).

25. Lloyd-Jones, L.R. et al. Improved polygenic prediction by Bayesian multiple regression on summary statistics. Nat Commun 10, 5086 (2019).

26. Weissbrod, O. et al. Functionally informed fine-mapping and polygenic localization of complex trait heritability. Nat Genet 52, 1355–1363 (2020).

27. Yang, S. & Zhou, X. Accurate and Scalable Construction of Polygenic Scores in Large Biobank Data Sets. Am J Hum Genet 106, 679–693 (2020).

28. Khan, A. et al. Genome-wide polygenic score to predict chronic kidney disease across ancestries. Nat Med 28, 1412–1420 (2022).

29. Park, T. & Casella, G. The Bayesian Lasso. Journal of the American Statistical Association 103, 681–686 (2008).

30. Carvalho, C.M., Polson, N.G. & Scott, J.G. Handling Sparsity via the Horseshoe. in Proceedings of the Twelth International Conference on Artificial Intelligence and Statistics Vol. 5 (eds David van, D. & Max, W.) 73--80 (PMLR, Proceedings of Machine Learning Research, 2009).

31. Polson, N.G., Scott, J.G. & Windle, J. The Bayesian bridge. Journal of the Royal Statistical Society: Series B (Statistical Methodology) 76, 713–733 (2014).

32. Truong, B. et al. Integrative polygenic risk score improves the prediction accuracy of complex traits and diseases. medRxiv, 2023.02.21.23286110 (2023).

33. Albiñana, C. et al. Multi-PGS enhances polygenic prediction: weighting 937 polygenic scores. medRxiv, 2022.09.14.22279940 (2022).

34. Locke, A.E. et al. Genetic studies of body mass index yield new insights for obesity biology. Nature 518, 197–206 (2015).

35. Willer, C.J. et al. Discovery and refinement of loci associated with lipid levels. Nat Genet 45, 1274–1283 (2013).

36. Akiyama, M. et al. Genome-wide association study identifies 112 new loci for body mass index in the Japanese population. Nat Genet 49, 1458–1467 (2017).

37. Kanai, M. et al. Genetic analysis of quantitative traits in the Japanese population links cell types to complex human diseases. Nat Genet 50, 390–400 (2018).

## References

7. Kachuri, L. et al. Principles and methods for transferring polygenic risk scores across global populations. Nat Rev Genet (2023).

8. Marquez-Luna, C., Loh, P.R., South Asian Type 2 Diabetes, C., Consortium, S.T.D. & Price, A.L. Multiethnic polygenic risk scores improve risk prediction in diverse populations. Genet Epidemiol 41, 811–823 (2017).

9. Ruan, Y. et al. Improving polygenic prediction in ancestrally diverse populations. Nature Genetics 54, 573–580 (2022).

10. Cai, M. et al. A unified framework for cross-population trait prediction by leveraging the genetic correlation of polygenic traits. Am J Hum Genet 108, 632–655 (2021).

11. Tian, P. et al. Multiethnic polygenic risk prediction in diverse populations through transfer learning. Front Genet 13, 906965 (2022).

12. Sun, Q. et al. Improving polygenic risk prediction in admixed populations by explicitly modeling ancestral-specific effects via GAUDI. bioRxiv, 2022.10.06.511219 (2022).

13. Zhang, H. et al. A new Method for Multi-ancestry Polygenic Prediction Improves Performance across Diverse Populations. bioRxiv, 2022.03.24.485519 (2022).

14. Privé, F., Arbel, J. & Vilhjálmsson, B.J. LDpred2: better, faster, stronger. Bioinformatics 36, 5424–5431 (2020).

15. Prive, F., Vilhjalmsson, B.J., Aschard, H. & Blum, M.G.B. Making the Most of Clumping and Thresholding for Polygenic Scores. Am J Hum Genet 105, 1213–1221 (2019).

16. van der Laan, M.J., Polley, E.C. & Hubbard, A.E. Super learner. Stat Appl Genet Mol Biol 6, Article25 (2007).

17. Zou, H. & Hastie, T. Regularization and Variable Selection via the Elastic Net. Journal of the Royal Statistical Society. Series B (Statistical Methodology) 67, 301–320 (2005).

18. Friedman, J., Hastie, T. & Tibshirani, R. Regularization Paths for Generalized Linear Models via Coordinate Descent. J Stat Softw 33, 1–22 (2010).

19. Weissbrod, O. et al. Leveraging fine-mapping and multipopulation training data to improve cross-population polygenic risk scores. Nat Genet 54, 450–458 (2022).

20. International HapMap, C. et al. Integrating common and rare genetic variation in diverse human populations. Nature 467, 52–8 (2010).

21. Bien, S.A. et al. Strategies for Enriching Variant Coverage in Candidate Disease Loci on a Multiethnic Genotyping Array. PLoS One 11, e0167758 (2016).

22. DiCiccio, T.J. & Efron, B. Bootstrap Confidence Intervals. Statistical Science 11, 189–212 (1996).

23. Canty, A.J.Resampling methods in R: the boot package. The Newsletter of the R Project Volume 2, 2–7 (2002).

24. Siva, N. 1000 Genomes project. Nature Biotechnology 26, 256–256 (2008).

25. Graham, S.E. et al. The power of genetic diversity in genome-wide association studies of lipids. Nature 600, 675–679 (2021).

26. Privé, F., Arbel, J., Aschard, H. & Vilhjálmsson, B.J. Identifying and correcting for misspecifications in GWAS summary statistics and polygenic scores. Human Genetics and Genomics Advances 3, 100136 (2022).

27. Ge, T., Chen, C.Y., Ni, Y., Feng, Y.A. & Smoller, J.W. Polygenic prediction via Bayesian regression and continuous shrinkage priors. Nat Commun 10, 1776 (2019).

28. Lloyd-Jones, L.R. et al. Improved polygenic prediction by Bayesian multiple regression on summary statistics. Nat Commun 10, 5086 (2019).

29. Weissbrod, O. et al. Functionally informed fine-mapping and polygenic localization of complex trait heritability. Nat Genet 52, 1355–1363 (2020).

30. Yang, S. & Zhou, X. Accurate and Scalable Construction of Polygenic Scores in Large Biobank Data Sets. Am J Hum Genet 106, 679–693 (2020).

31. Khan, A. et al. Genome-wide polygenic score to predict chronic kidney disease across ancestries. Nat Med 28, 1412–1420 (2022).

32. Park, T. & Casella, G. The Bayesian Lasso. Journal of the American Statistical Association 103, 681–686 (2008).

33. Carvalho, C.M., Polson, N.G. & Scott, J.G. Handling Sparsity via the Horseshoe. in Proceedings of the Twelth International Conference on Artificial Intelligence and Statistics Vol. 5 (eds David van, D. & Max, W.) 73--80 (PMLR, Proceedings of Machine Learning Research, 2009).

34. Polson, N.G., Scott, J.G. & Windle, J. The Bayesian bridge. Journal of the Royal Statistical Society: Series B (Statistical Methodology) 76, 713–733 (2014).

35. Truong, B. et al. Integrative polygenic risk score improves the prediction accuracy of complex traits and diseases. medRxiv, 2023.02.21.23286110 (2023).

36. Albiñana, C. et al. Multi-PGS enhances polygenic prediction: weighting 937 polygenic scores. medRxiv, 2022.09.14.22279940 (2022).

37. Locke, A.E. et al. Genetic studies of body mass index yield new insights for obesity biology. Nature 518, 197–206 (2015).

38. Willer, C.J. et al. Discovery and refinement of loci associated with lipid levels. Nat Genet 45, 1274–1283 (2013).

39. Akiyama, M. et al. Genome-wide association study identifies 112 new loci for body mass index in the Japanese population. Nat Genet 49, 1458–1467 (2017).

40. Kanai, M. et al. Genetic analysis of quantitative traits in the Japanese population links cell types to complex human diseases. Nat Genet 50, 390–400 (2018).

41. Berisa, T. & Pickrell, J.K. Approximately independent linkage disequilibrium blocks in human populations. Bioinformatics 32, 283–5 (2016).

42. Shaun Purcell, C.C. PLINK 2.0. URL: http://www.cog-genomics.org/plink/2.0/.

43. Mak, T.S.H., Porsch, R.M., Choi, S.W., Zhou, X. & Sham, P.C. Polygenic scores via penalized regression on summary statistics. Genet Epidemiol 41, 469–480 (2017).

44. Bulik-Sullivan, B.K. et al. LD Score regression distinguishes confounding from polygenicity in genome-wide association studies. Nat Genet 47, 291–5 (2015).

45. Vilhjalmsson, B.J. et al. Modeling Linkage Disequilibrium Increases Accuracy of Polygenic Risk Scores. Am J Hum Genet 97, 576–92 (2015).

46. Chang, C.C. et al. Second-generation PLINK: rising to the challenge of larger and richer datasets. Gigascience 4, 7 (2015).

47. Shaun Purcell and Christopher Chang. PLINK 1.90. Vol. 2022.

48. Prive, F., Arbel, J. & Vilhjalmsson, B.J. LDpred2: better, faster, stronger. Bioinformatics (2020).

49. Sudlow, C. et al. UK biobank: an open access resource for identifying the causes of a wide range of complex diseases of middle and old age. PLoS Med 12, e1001779 (2015).

50. Friedewald, W.T., Levy, R.I. & Fredrickson, D.S. Estimation of the concentration of low-density lipoprotein cholesterol in plasma, without use of the preparative ultracentrifuge. Clin Chem 18, 499–502 (1972).

51. Siva, N. 1000 Genomes project. 26, 256 (2008).

