## Supplementary Figures and Notes for "MUSSEL: Enhanced Bayesian Polygenic Risk Prediction Leveraging Information across Multiple Ancestry Groups"

**Supplementary Figure 1: Simulation results showing performance of the PRS constructed by MUSSEL and various existing methods, assuming a fixed common SNP heritability (0.4) across ancestries under a strong negative selection model for the relationship between SNP effect size and allele frequency.** The genetic correlation in SNP effect size is set to 0.8 across all pairs of populations. The causal SNP proportion (degree of polygenicity) is set to 1.0%, 0.1%, or 0.05% (~192$K$, 19.2$K$, or 9.6$K$ causal SNPs). We generate data for ~19 million common SNPs (MAF≥1%) across the five ancestry groups but conduct analyses only on the ~2.0 million SNPs in HapMap 3 + MEGA. The discovery GWAS sample size is set to **(a) 15,000 or (b) 45,000** for each non-EUR ancestry, and 100,000 for EUR. A tuning set consisting of 10,000 individuals is used for parameter tuning, as well as training the SL in CT-SLEB and MUSSEL or the linear combination model in weighted C+T, weighted LDpred2, PRS-CSx, and weighted MUSS. The reported $R^2$ values are calculated on an independent testing set of 10,000 individuals for each ancestry group. The corresponding 95% bootstrap CIs are obtained from the same testing set based on 10,000 bootstrap samples using the Bca approach[1] implemented in the R package "boot".

(a)

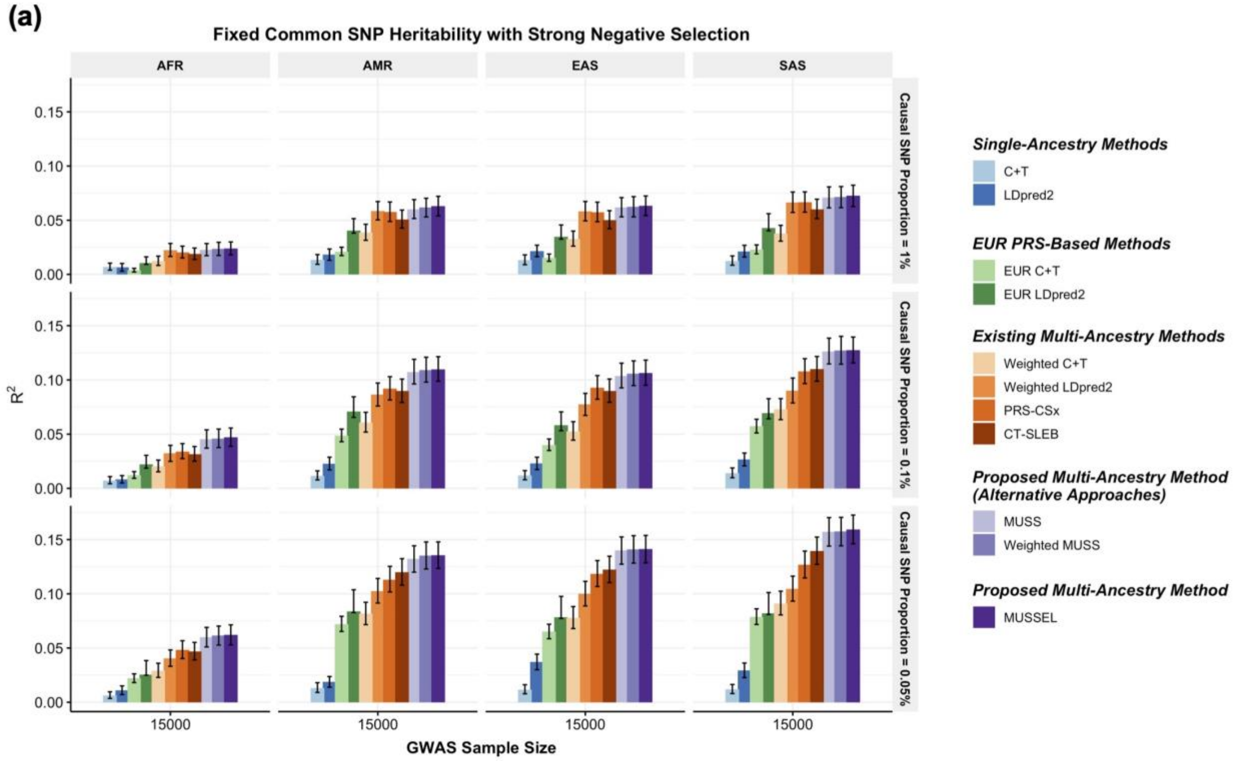
Fixed Common SNP Heritability with Strong Negative Selection
AFR
AMR
EAS
SAS
Causal SNP Proportion = 1%
Causal SNP Proportion = 0.1%
Causal SNP Proportion = 0.05%
R2
0.15
0.10
0.05
0.00
15000
GWAS Sample Size
Single-Ancestry Methods
C+T
LDpred2
EUR PRS-Based Methods
EUR C+T
EUR LDpred2
Existing Multi-Ancestry Methods
Weighted C+T
Weighted LDpred2
PRS-CSx
CT-SLEB
Proposed Multi-Ancestry Method (Alternative Approaches)
MUSS
Weighted MUSS
Proposed Multi-Ancestry Method
MUSSEL

(b)

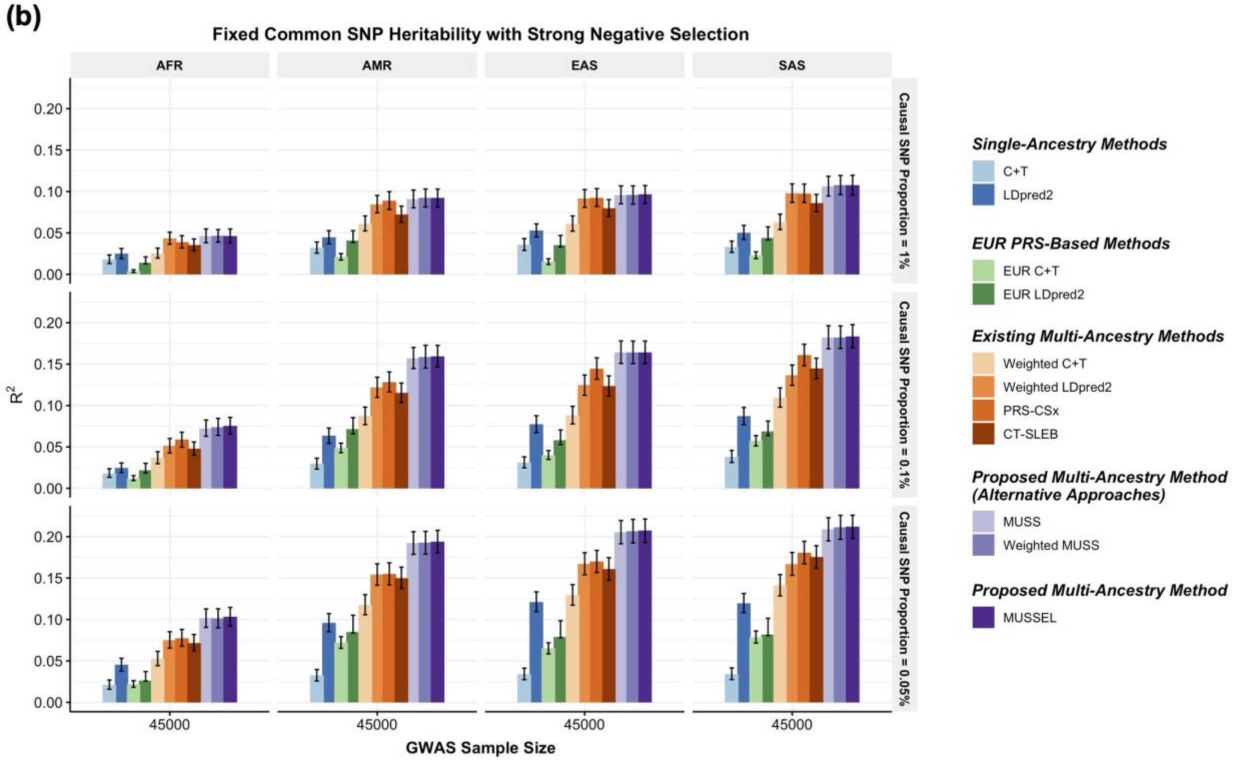
Fixed Common SNP Heritability with Strong Negative Selection
AFR
AMR
EAS
SAS
Causal SNP Proportion = 1%
Causal SNP Proportion = 0.1%
Causal SNP Proportion = 0.05%
R2
0.20
0.15
0.10
0.05
0.00
45000
GWAS Sample Size
Single-Ancestry Methods
C+T
LDpred2
EUR PRS-Based Methods
EUR C+T
EUR LDpred2
Existing Multi-Ancestry Methods
Weighted C+T
Weighted LDpred2
PRS-CSx
CT-SLEB
Proposed Multi-Ancestry Method (Alternative Approaches)
MUSS
Weighted MUSS
Proposed Multi-Ancestry Method
MUSSEL

**Supplementary Figure 2: Simulation results showing performance of the PRS constructed by MUSSEL and various existing methods, assuming a fixed common SNP heritability (0.4) across ancestries under a strong negative selection model for the relationship between SNP effect size and allele frequency.** The genetic correlation in SNP effect size is set to 0.8 across all pairs of populations. The causal SNP proportion (degree of polygenicity) is set to 1.0%, 0.1%, or 0.05% (~192$K$, 19.2$K$, or 9.6$K$ causal SNPs). We generate data for ~19 million common SNPs (MAF≥1%) across the five ancestries but conduct analyses only on the ~2.0 million SNPs in HapMap 3 + MEGA. The discovery GWAS sample size is set to **(a) 80,000 or (b) 100,000** for each non-EUR ancestry, and 100,000 for EUR. A tuning set consisting of 10,000 individuals is used for parameter tuning, as well as training the SL in CT-SLEB and MUSSEL or the linear combination model in weighted C+T, weighted LDpred2, PRS-CSx, and weighted MUSS. The reported $R^2$ values are calculated on an independent testing set of 10,000 individuals for each ancestry group. The corresponding 95% bootstrap CIs are obtained from the same testing set based on 10,000 bootstrap samples using the Bca approach[1] implemented in the R package "boot".

(a)

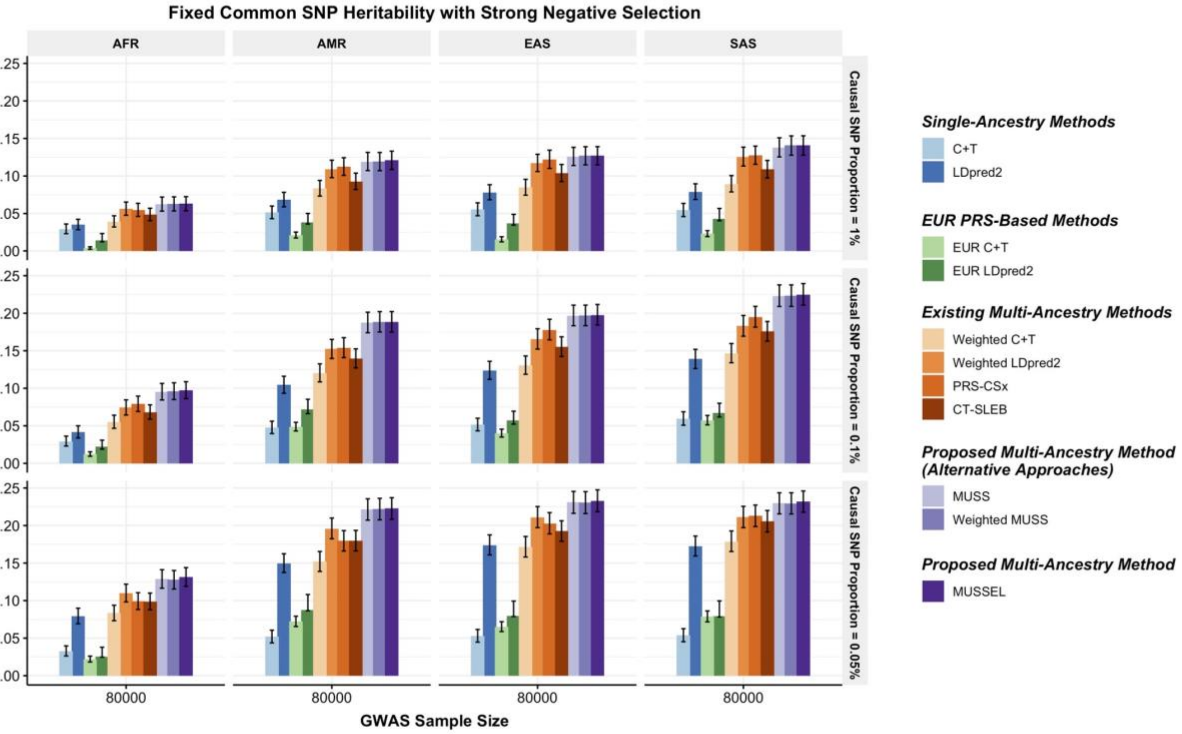
Fixed Common SNP Heritability with Strong Negative Selection
AFR
AMR
EAS
SAS
Causal SNP Proportion = 1%
Causal SNP Proportion = 0.1%
Causal SNP Proportion = 0.05%
$R^2$
0.00
0.05
0.10
0.15
0.20
0.25
80000
GWAS Sample Size
Single-Ancestry Methods
C+T
LDpred2
EUR PRS-Based Methods
EUR C+T
EUR LDpred2
Existing Multi-Ancestry Methods
Weighted C+T
Weighted LDpred2
PRS-CSx
CT-SLEB
Proposed Multi-Ancestry Method (Alternative Approaches)
MUSS
Weighted MUSS
Proposed Multi-Ancestry Method
MUSSEL

(b)

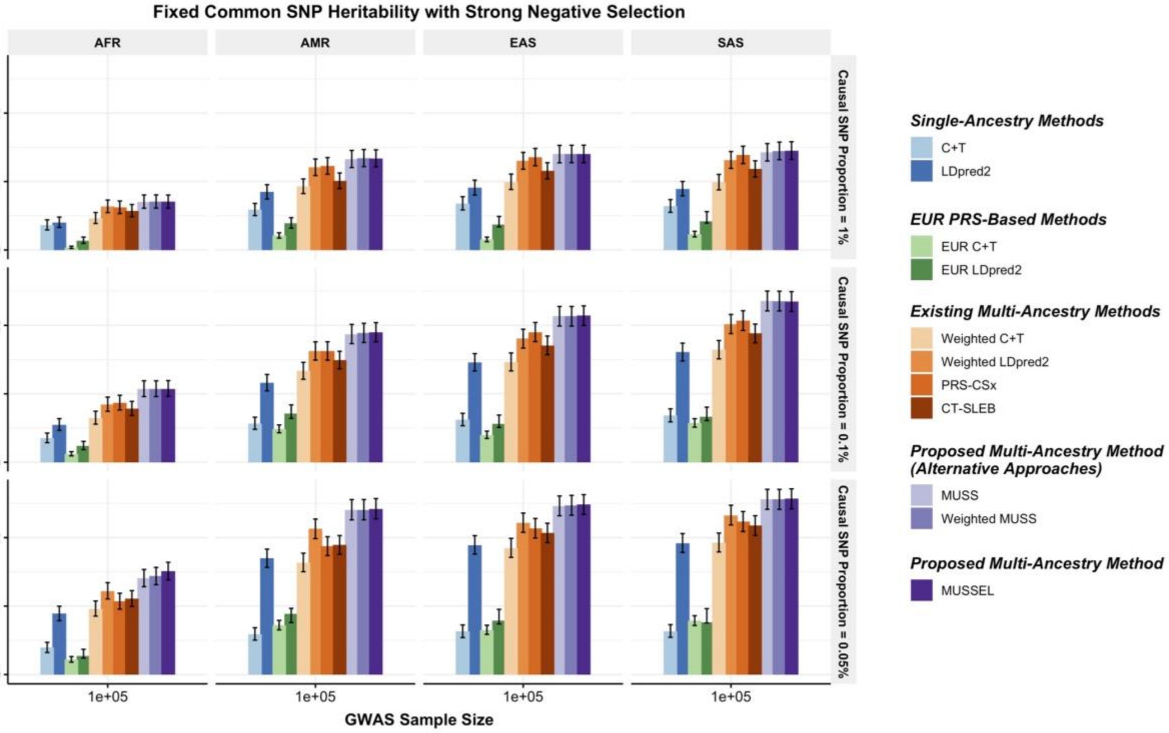
Fixed Common SNP Heritability with Strong Negative Selection
AFR
AMR
EAS
SAS
Causal SNP Proportion = 1%
Causal SNP Proportion = 0.1%
Causal SNP Proportion = 0.05%
$R^2$
0.0
0.1
0.2
1e+05
GWAS Sample Size
Single-Ancestry Methods
C+T
LDpred2
EUR PRS-Based Methods
EUR C+T
EUR LDpred2
Existing Multi-Ancestry Methods
Weighted C+T
Weighted LDpred2
PRS-CSx
CT-SLEB
Proposed Multi-Ancestry Method (Alternative Approaches)
MUSS
Weighted MUSS
Proposed Multi-Ancestry Method
MUSSEL

**Supplementary Figure 3: Simulation results showing performance of the PRS constructed by MUSSEL and various existing methods, assuming a fixed per-SNP heritability (0.4) across ancestries under a strong negative selection model for the relationship between SNP effect size and allele frequency.** The genetic correlation in SNP effect size is set to 0.8 across all pairs of populations. The causal SNP proportion (degree of polygenicity) is set to 1.0%, 0.1%, or 0.05% (~192$K$, 19.2$K$, or 9.6$K$ causal SNPs). We generate data for ~19 million common SNPs (MAF≥1%) across the five ancestries but conduct analyses only on the ~2.0 million SNPs in HapMap 3 + MEGA. The discovery GWAS sample size is set to **(a) 15,000 or (b) 45,000** for each non-EUR ancestry, and 100,000 for EUR. A tuning set consisting of 10,000 individuals is used for parameter tuning, as well as training the SL in CT-SLEB and MUSSEL or the linear combination model in weighted C+T, weighted LDpred2, PRS-CSx, and weighted MUSS. The reported $R^2$ values are calculated on an independent testing set of 10,000 individuals for each ancestry group. The corresponding 95% bootstrap CIs are obtained from the same testing set based on 10,000 bootstrap samples using the Bca approach[1] implemented in the R package “boot”.

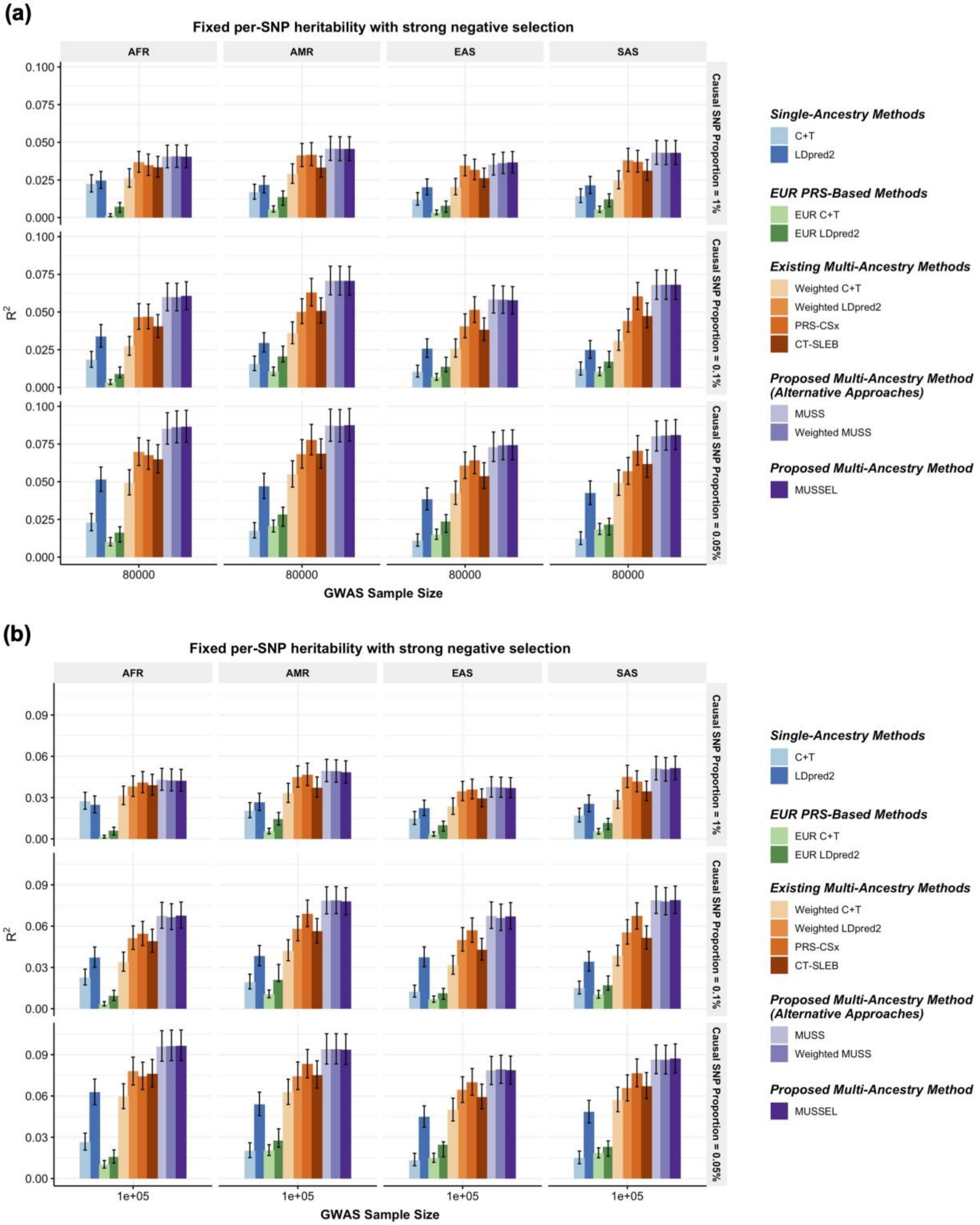
(a)
Fixed per-SNP heritability with strong negative selection
AFR
AMR
EAS
SAS
Causal SNP Proportion = 1%
Causal SNP Proportion = 0.1%
Causal SNP Proportion = 0.05%
R2
GWAS Sample Size
80000
Single-Ancestry Methods
C+T
LDpred2
EUR PRS-Based Methods
EUR C+T
EUR LDpred2
Existing Multi-Ancestry Methods
Weighted C+T
Weighted LDpred2
PRS-CSx
CT-SLEB
Proposed Multi-Ancestry Method (Alternative Approaches)
MUSS
Weighted MUSS
Proposed Multi-Ancestry Method
MUSSEL
(b)
Fixed per-SNP heritability with strong negative selection
1e+05
GWAS Sample Size

**Supplementary Figure 4: Simulation results showing performance of the PRS constructed by MUSSEL and various existing methods, assuming a fixed per-SNP heritability (0.4) across ancestries under a strong negative selection model for the relationship between SNP effect size and allele frequency.** The genetic correlation in SNP effect size is set to 0.8 across all pairs of populations. The causal SNP proportion (degree of polygenicity) is set to 1.0%, 0.1%, or 0.05% (~192$K$, 19.2$K$, or 9.6$K$ causal SNPs). We generate data for ~19 million common SNPs (MAF≥1%) across the five ancestries but conduct analyses only on the ~2.0 million SNPs in HapMap 3 + MEGA. The discovery GWAS sample size is set to **(a) 80,000 or (b) 100,000** for each non-EUR ancestry, and 100,000 for EUR. A tuning set consisting of 10,000 individuals is used for parameter tuning, as well as training the SL in CT-SLEB and MUSSEL or the linear combination model in weighted C+T, weighted LDpred2, PRS-CSx, and weighted MUSS. The reported $R^2$ values are calculated on an independent testing set of 10,000 individuals for each ancestry group. The corresponding 95% bootstrap CIs are obtained from the same testing set based on 10,000 bootstrap samples using the Bca approach[1] implemented in the R package “boot”.

(a)

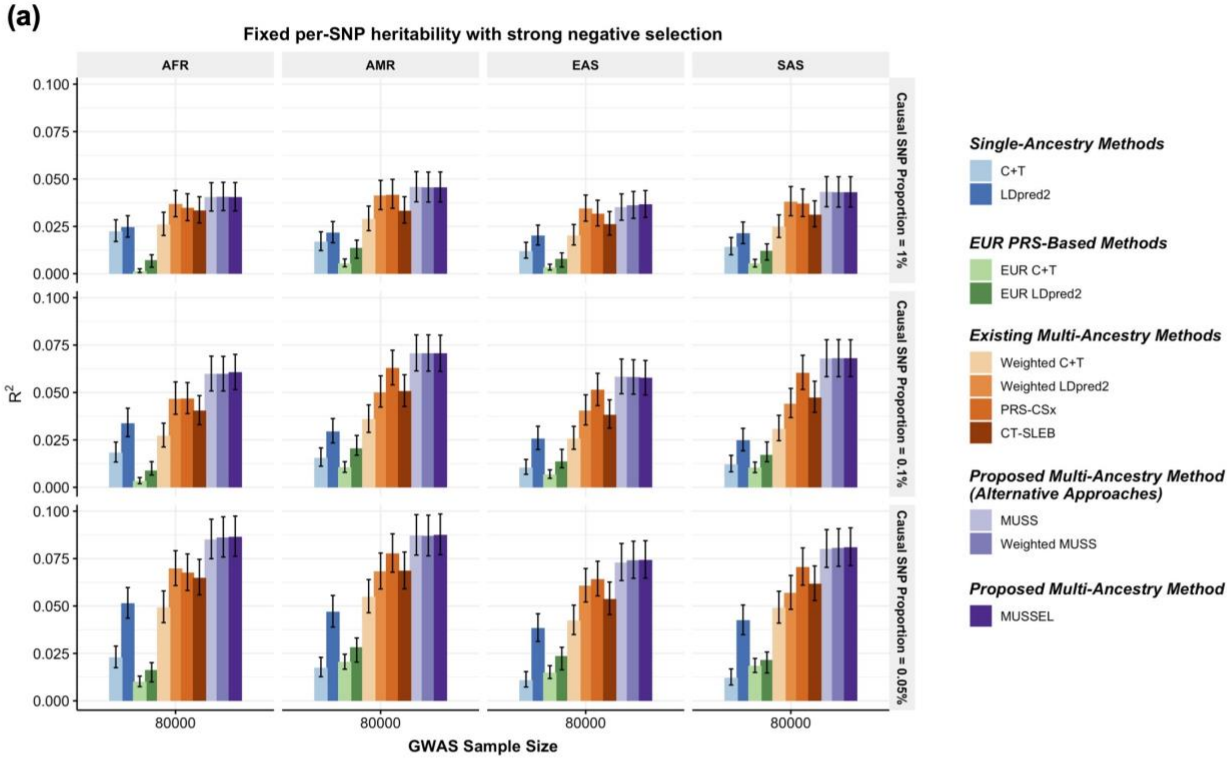
Fixed per-SNP heritability with strong negative selection
AFR
AMR
EAS
SAS
Causal SNP Proportion = 1%
Causal SNP Proportion = 0.1%
Causal SNP Proportion = 0.05%
0.100
0.075
0.050
0.025
0.000
R²
80000
GWAS Sample Size
Single-Ancestry Methods
C+T
LDpred2
EUR PRS-Based Methods
EUR C+T
EUR LDpred2
Existing Multi-Ancestry Methods
Weighted C+T
Weighted LDpred2
PRS-CSx
CT-SLEB
Proposed Multi-Ancestry Method (Alternative Approaches)
MUSS
Weighted MUSS
Proposed Multi-Ancestry Method
MUSSEL

(b)

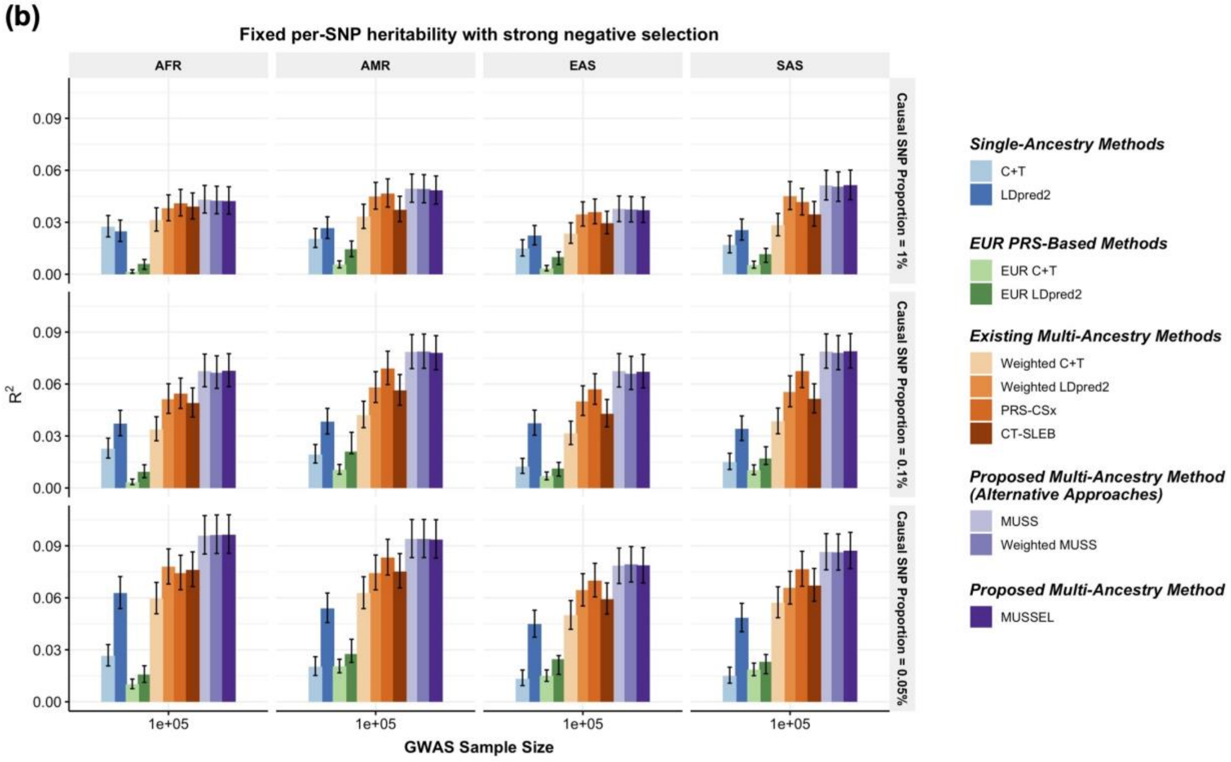
Fixed per-SNP heritability with strong negative selection
AFR
AMR
EAS
SAS
Causal SNP Proportion = 1%
Causal SNP Proportion = 0.1%
Causal SNP Proportion = 0.05%
0.09
0.06
0.03
0.00
R²
1e+05
GWAS Sample Size
Single-Ancestry Methods
C+T
LDpred2
EUR PRS-Based Methods
EUR C+T
EUR LDpred2
Existing Multi-Ancestry Methods
Weighted C+T
Weighted LDpred2
PRS-CSx
CT-SLEB
Proposed Multi-Ancestry Method (Alternative Approaches)
MUSS
Weighted MUSS
Proposed Multi-Ancestry Method
MUSSEL

**Supplementary Figure 5: Simulation results showing performance of the PRS constructed by MUSSEL and various existing methods, assuming a fixed per-SNP heritability (0.4) across ancestries under a strong negative selection model for the relationship between SNP effect size and allele frequency but with weaker cross-population (0.6 across all pairs of populations).** The causal SNP proportion (degree of polygenicity) is set to 1.0%, 0.1%, or 0.05% (~192$K$, 19.2$K$, or 9.6$K$ causal SNPs). We generate data for ~19 million common SNPs (MAF≥1%) across the five ancestries but conduct analyses only on the ~2.0 million SNPs in HapMap 3 + MEGA. The discovery GWAS sample size is set to **(a) 15,000 or (b) 45,000** for each non-EUR ancestry, and 100,000 for EUR. A tuning set consisting of 10,000 individuals is used for parameter tuning, as well as training the SL in CT-SLEB and MUSSEL or the linear combination model in weighted C+T, weighted LDpred2, PRS-CSx, and weighted MUSS. The reported $R^2$ values are calculated on an independent testing set of 10,000 individuals for each ancestry group. The corresponding 95% bootstrap CIs are obtained from the same testing set based on 10,000 bootstrap samples using the Bca approach[1] implemented in the R package “boot”.

**(a)**

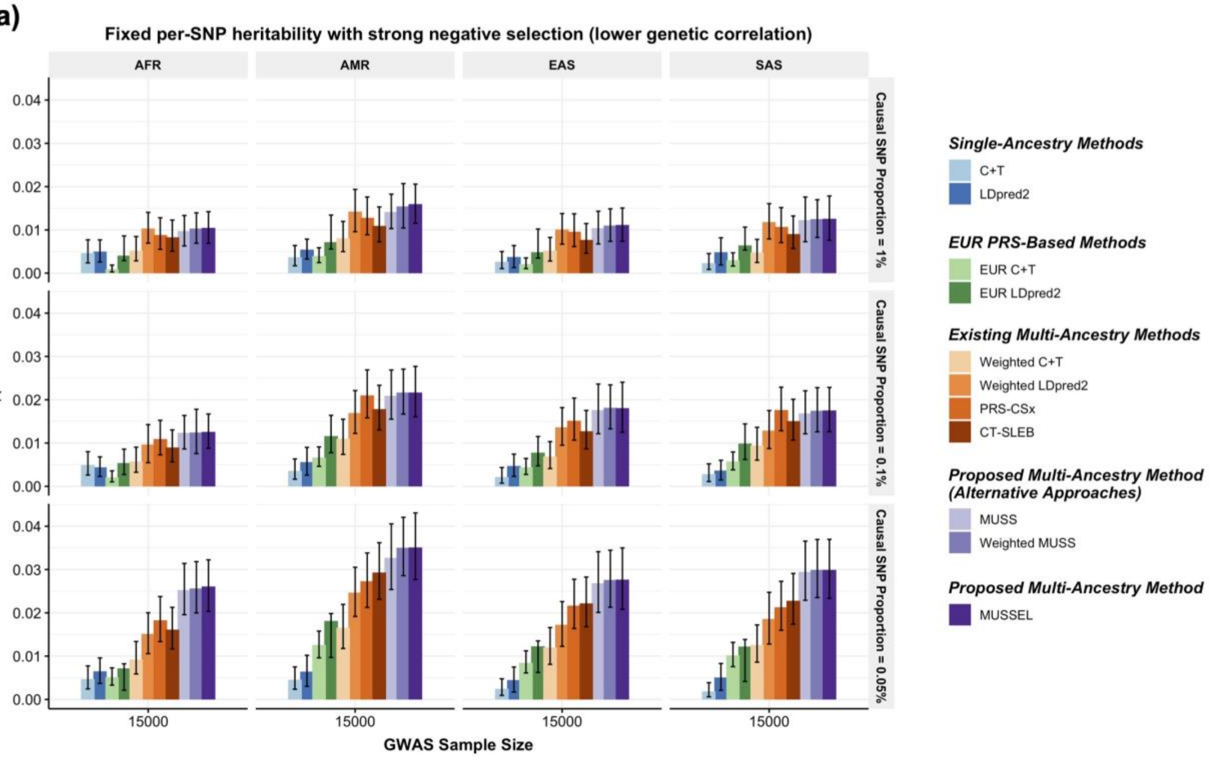

Fixed per-SNP heritability with strong negative selection (lower genetic correlation)
AFR
AMR
EAS
SAS
Causal SNP Proportion = 1%
Causal SNP Proportion = 0.1%
Causal SNP Proportion = 0.05%
$R^2$
0.00
0.01
0.02
0.03
0.04
15000
GWAS Sample Size
Single-Ancestry Methods
C+T
LDpred2
EUR PRS-Based Methods
EUR C+T
EUR LDpred2
Existing Multi-Ancestry Methods
Weighted C+T
Weighted LDpred2
PRS-CSx
CT-SLEB
Proposed Multi-Ancestry Method (Alternative Approaches)
MUSS
Weighted MUSS
Proposed Multi-Ancestry Method
MUSSEL


**(b)**

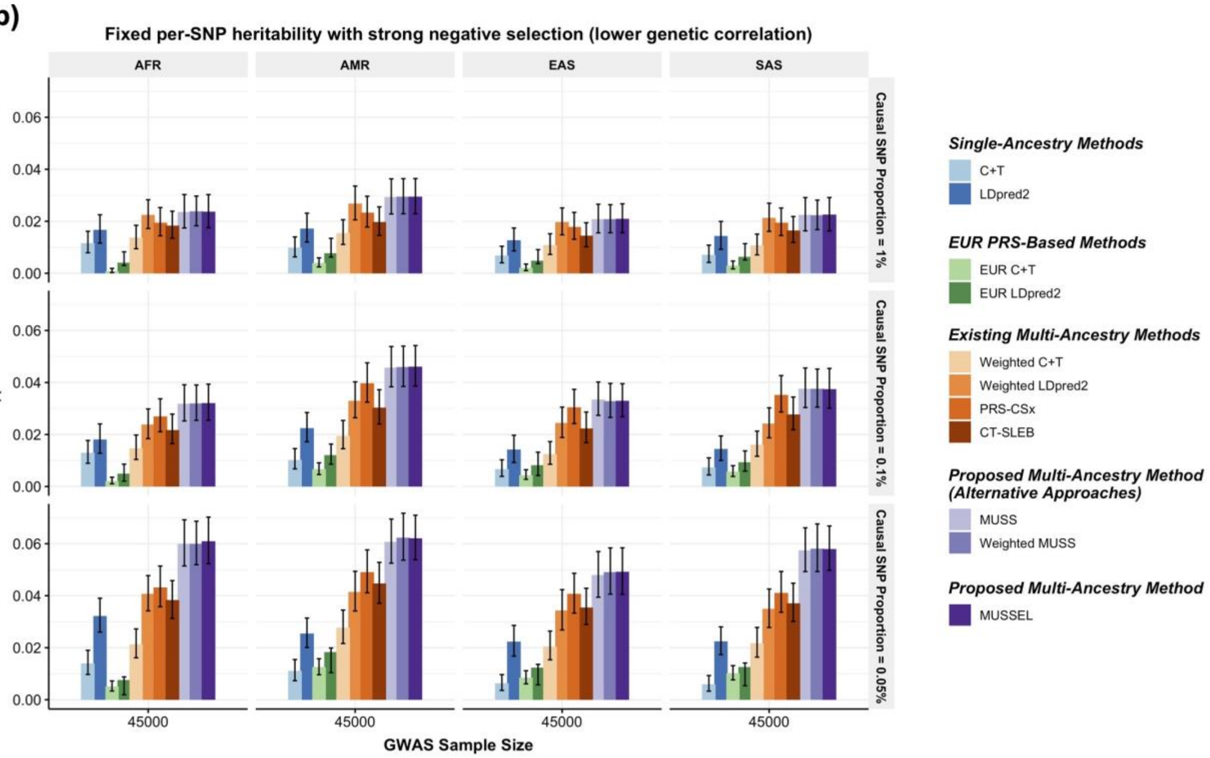

Fixed per-SNP heritability with strong negative selection (lower genetic correlation)
AFR
AMR
EAS
SAS
Causal SNP Proportion = 1%
Causal SNP Proportion = 0.1%
Causal SNP Proportion = 0.05%
$R^2$
0.00
0.02
0.04
0.06
45000
GWAS Sample Size
Single-Ancestry Methods
C+T
LDpred2
EUR PRS-Based Methods
EUR C+T
EUR LDpred2
Existing Multi-Ancestry Methods
Weighted C+T
Weighted LDpred2
PRS-CSx
CT-SLEB
Proposed Multi-Ancestry Method (Alternative Approaches)
MUSS
Weighted MUSS
Proposed Multi-Ancestry Method
MUSSEL

**Supplementary Figure 6: Simulation results showing performance of the PRS constructed by MUSSEL and various existing methods, assuming a fixed per-SNP heritability (0.4) across ancestries under a strong negative selection model for the relationship between SNP effect size and allele frequency but with weaker cross-population (0.6 across all pairs of populations).** The causal SNP proportion (degree of polygenicity) is set to 1.0%, 0.1%, or 0.05% (~192$K$, 19.2$K$, or 9.6$K$ causal SNPs). We generate data for ~19 million common SNPs (MAF≥1%) across the five ancestries but conduct analyses only on the ~2.0 million SNPs in HapMap 3 + MEGA. The discovery GWAS sample size is set to **(a) 80,000 or (b) 100,000** for each non-EUR ancestry, and 100,000 for EUR. A tuning set consisting of 10,000 individuals is used for parameter tuning, as well as training the SL in CT-SLEB and MUSSEL or the linear combination model in weighted C+T, weighted LDpred2, PRS-CSx, and weighted MUSS. The reported $R^2$ values are calculated on an independent testing set of 10,000 individuals for each ancestry group. The corresponding 95% bootstrap CIs are obtained from the same testing set based on 10,000 bootstrap samples using the Bca approach[1] implemented in the R package "boot".

**(a)**

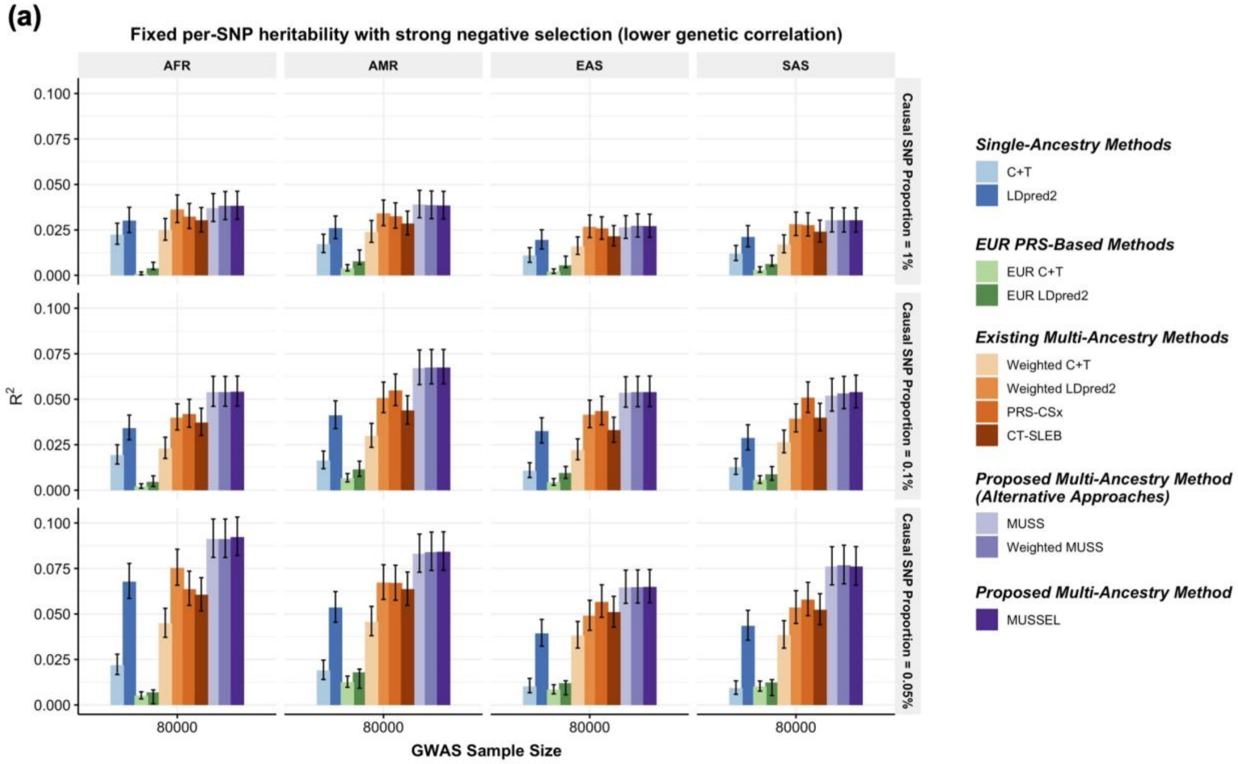

Fixed per-SNP heritability with strong negative selection (lower genetic correlation)
AFR
AMR
EAS
SAS
0.100
0.075
0.050
0.025
0.000
$R^2$
Causal SNP Proportion = 1%
Causal SNP Proportion = 0.1%
Causal SNP Proportion = 0.05%
80000
GWAS Sample Size
Single-Ancestry Methods
C+T
LDpred2
EUR PRS-Based Methods
EUR C+T
EUR LDpred2
Existing Multi-Ancestry Methods
Weighted C+T
Weighted LDpred2
PRS-CSx
CT-SLEB
Proposed Multi-Ancestry Method (Alternative Approaches)
MUSS
Weighted MUSS
Proposed Multi-Ancestry Method
MUSSEL


**(b)**

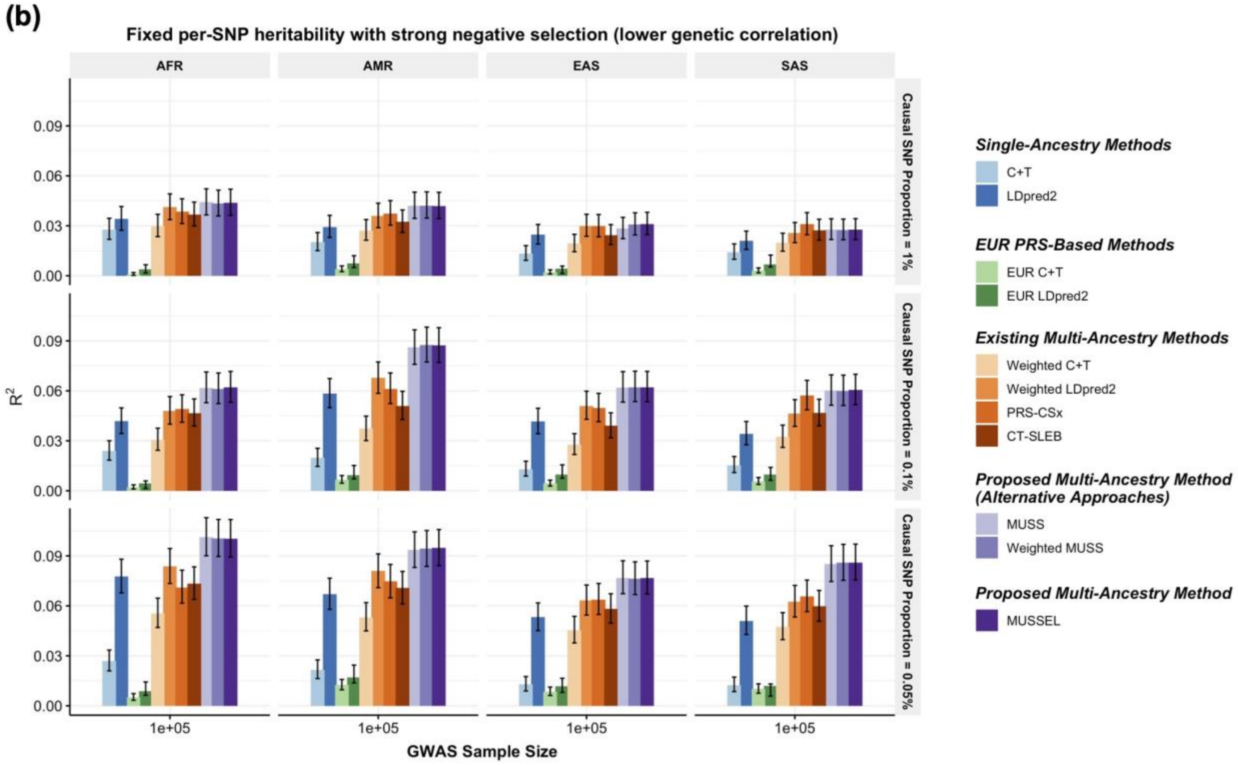

Fixed per-SNP heritability with strong negative selection (lower genetic correlation)
AFR
AMR
EAS
SAS
0.09
0.06
0.03
0.00
$R^2$
Causal SNP Proportion = 1%
Causal SNP Proportion = 0.1%
Causal SNP Proportion = 0.05%
1e+05
GWAS Sample Size
Single-Ancestry Methods
C+T
LDpred2
EUR PRS-Based Methods
EUR C+T
EUR LDpred2
Existing Multi-Ancestry Methods
Weighted C+T
Weighted LDpred2
PRS-CSx
CT-SLEB
Proposed Multi-Ancestry Method (Alternative Approaches)
MUSS
Weighted MUSS
Proposed Multi-Ancestry Method
MUSSEL

**Supplementary Figure 7: Simulation results showing performance of the PRS constructed by MUSSEL and various existing methods, assuming a fixed common SNP heritability (0.4) across ancestries with no negative selection for the relationship between SNP effect size and allele frequency.** The genetic correlation in SNP effect size is set to 0.8 across all pairs of populations. The causal SNP proportion (degree of polygenicity) is set to 1.0%, 0.1%, or 0.05% (~192$K$, 19.2$K$, or 9.6$K$ causal SNPs). We generate data for ~19 million common SNPs (MAF≥1%) across the five ancestries but conduct analyses only on the ~2.0 million SNPs in HapMap 3 + MEGA. The discovery GWAS sample size is set to **(a) 15,000 or (b) 45,000** for each non-EUR ancestry, and 100,000 for EUR. A tuning set consisting of 10,000 individuals is used for parameter tuning, as well as training the SL in CT-SLEB and MUSSEL or the linear combination model in weighted C+T, weighted LDpred2, PRS-CSx, and weighted MUSS. The reported $R^2$ values are calculated on an independent testing set of 10,000 individuals for each ancestry group. The corresponding 95% bootstrap CIs are obtained from the same testing set based on 10,000 bootstrap samples using the Bca approach[1] implemented in the R package “boot”.

(a)

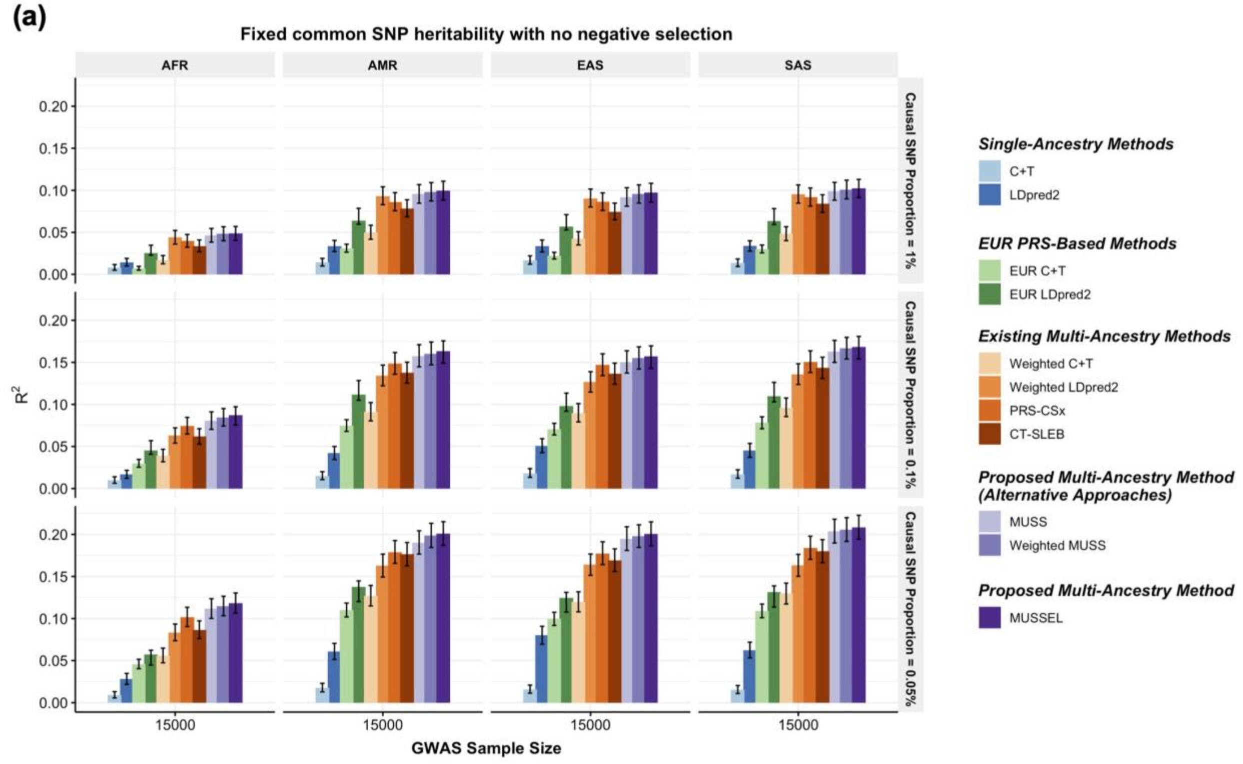

Fixed common SNP heritability with no negative selection
AFR
AMR
EAS
SAS
Causal SNP Proportion = 1%
Causal SNP Proportion = 0.1%
Causal SNP Proportion = 0.05%
$R^2$
0.00
0.05
0.10
0.15
0.20
15000
GWAS Sample Size
Single-Ancestry Methods
C+T
LDpred2
EUR PRS-Based Methods
EUR C+T
EUR LDpred2
Existing Multi-Ancestry Methods
Weighted C+T
Weighted LDpred2
PRS-CSx
CT-SLEB
Proposed Multi-Ancestry Method (Alternative Approaches)
MUSS
Weighted MUSS
Proposed Multi-Ancestry Method
MUSSEL


(b)

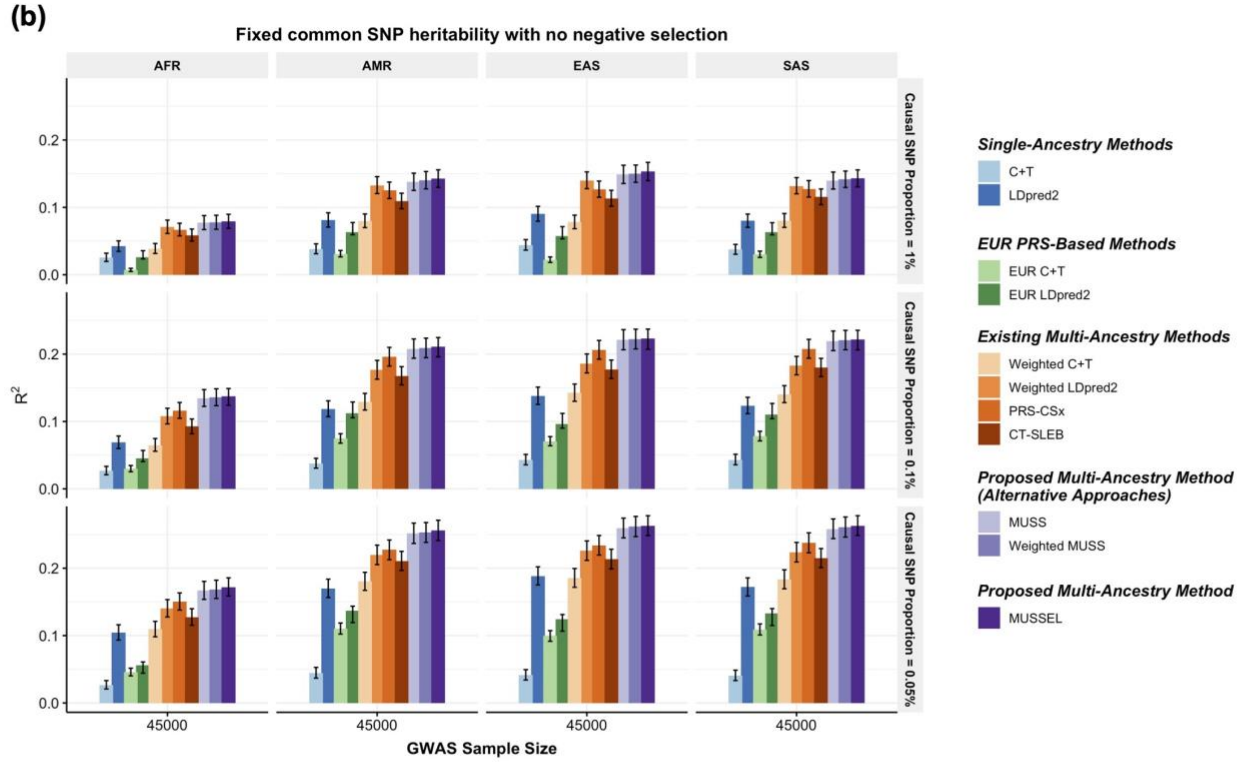

Fixed common SNP heritability with no negative selection
AFR
AMR
EAS
SAS
Causal SNP Proportion = 1%
Causal SNP Proportion = 0.1%
Causal SNP Proportion = 0.05%
$R^2$
0.0
0.1
0.2
45000
GWAS Sample Size
Single-Ancestry Methods
C+T
LDpred2
EUR PRS-Based Methods
EUR C+T
EUR LDpred2
Existing Multi-Ancestry Methods
Weighted C+T
Weighted LDpred2
PRS-CSx
CT-SLEB
Proposed Multi-Ancestry Method (Alternative Approaches)
MUSS
Weighted MUSS
Proposed Multi-Ancestry Method
MUSSEL

**Supplementary Figure 8: Simulation results showing performance of the PRS constructed by MUSSEL and various existing methods, assuming a fixed common SNP heritability (0.4) across ancestries with no negative selection for the relationship between SNP effect size and allele frequency.** The genetic correlation in SNP effect size is set to 0.8 across all pairs of populations. The causal SNP proportion (degree of polygenicity) is set to 1.0%, 0.1%, or 0.05% (~192$K$, 19.2$K$, or 9.6$K$ causal SNPs). We generate data for ~19 million common SNPs (MAF≥1%) across the five ancestries but conduct analyses only on the ~2.0 million SNPs in HapMap 3 + MEGA. The discovery GWAS sample size is set to **(a) 80,000 or (b) 100,000** for each non-EUR ancestry, and 100,000 for EUR. A tuning set consisting of 10,000 individuals is used for parameter tuning, as well as training the SL in CT-SLEB and MUSSEL or the linear combination model in weighted C+T, weighted LDpred2, PRS-CSx, and weighted MUSS. The reported $R^2$ values are calculated on an independent testing set of 10,000 individuals for each ancestry group. The corresponding 95% bootstrap CIs are obtained from the same testing set based on 10,000 bootstrap samples using the Bca approach[1] implemented in the R package “boot”.

(a)

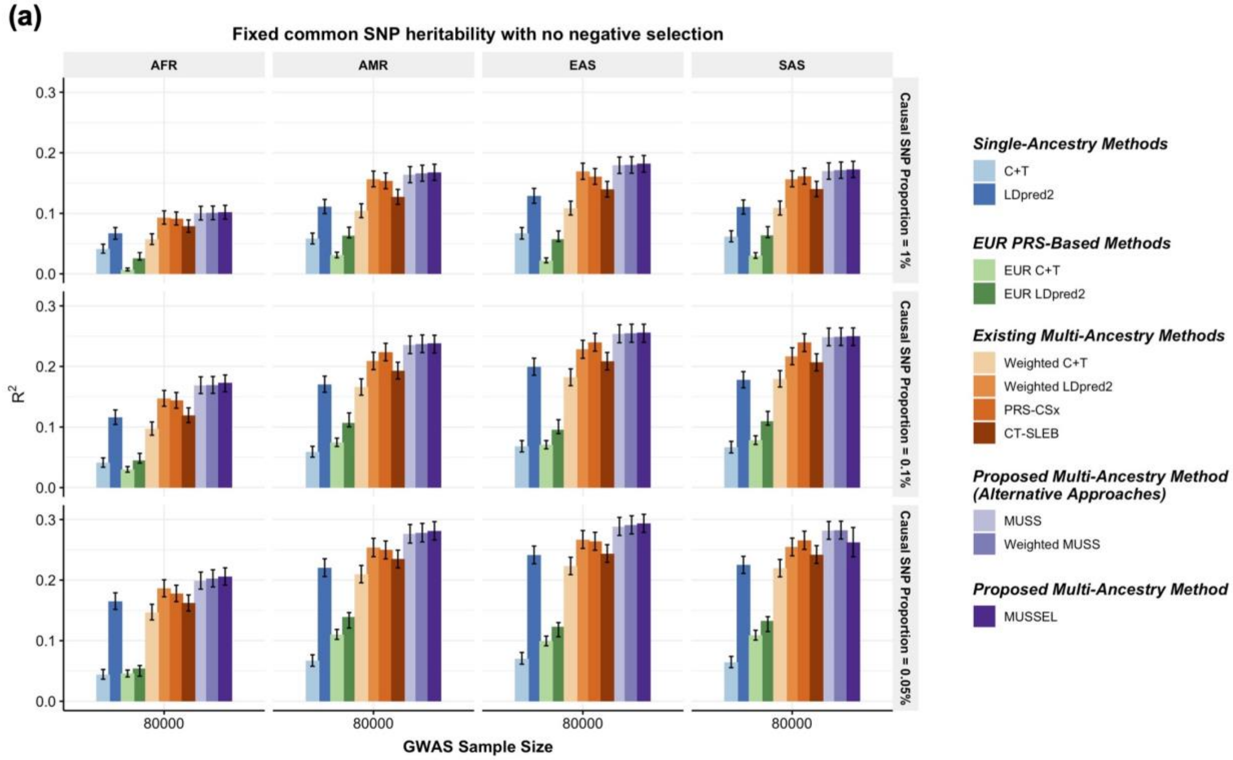
Fixed common SNP heritability with no negative selection
AFR
AMR
EAS
SAS
Causal SNP Proportion = 1%
Causal SNP Proportion = 0.1%
Causal SNP Proportion = 0.05%
$R^2$
0.0
0.1
0.2
0.3
80000
GWAS Sample Size
Single-Ancestry Methods
C+T
LDpred2
EUR PRS-Based Methods
EUR C+T
EUR LDpred2
Existing Multi-Ancestry Methods
Weighted C+T
Weighted LDpred2
PRS-CSx
CT-SLEB
Proposed Multi-Ancestry Method (Alternative Approaches)
MUSS
Weighted MUSS
Proposed Multi-Ancestry Method
MUSSEL

(b)

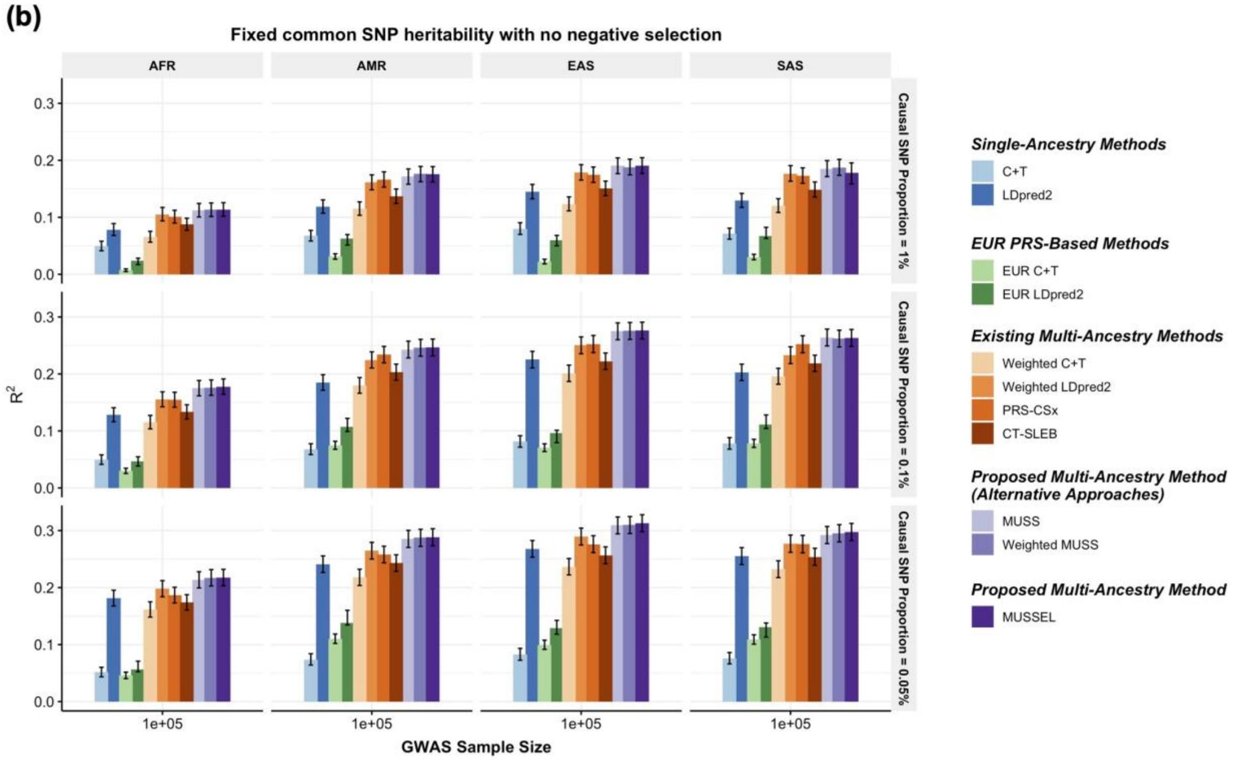
Fixed common SNP heritability with no negative selection
AFR
AMR
EAS
SAS
Causal SNP Proportion = 1%
Causal SNP Proportion = 0.1%
Causal SNP Proportion = 0.05%
$R^2$
0.0
0.1
0.2
0.3
1e+05
GWAS Sample Size
Single-Ancestry Methods
C+T
LDpred2
EUR PRS-Based Methods
EUR C+T
EUR LDpred2
Existing Multi-Ancestry Methods
Weighted C+T
Weighted LDpred2
PRS-CSx
CT-SLEB
Proposed Multi-Ancestry Method (Alternative Approaches)
MUSS
Weighted MUSS
Proposed Multi-Ancestry Method
MUSSEL

**Supplementary Figure 9: Simulation results showing performance of the PRS constructed by MUSSEL and various existing methods, assuming a fixed common SNP heritability (0.4) across ancestries under a mild negative selection model for the relationship between SNP effect size and allele frequency.** The genetic correlation in SNP effect size is set to 0.8 across all pairs of populations. The causal SNP proportion (degree of polygenicity) is set to 1.0%, 0.1%, or 0.05% (~192$K$, 19.2$K$, or 9.6$K$ causal SNPs). We generate data for ~19 million common SNPs (MAF≥1%) across the five ancestries but conduct analyses only on the ~2.0 million SNPs in HapMap 3 + MEGA. The discovery GWAS sample size is set to **(a) 15,000 or (b) 45,000** for each non-EUR ancestry, and 100,000 for EUR. A tuning set consisting of 10,000 individuals is used for parameter tuning, as well as training the SL in CT-SLEB and MUSSEL or the linear combination model in weighted C+T, weighted LDpred2, PRS-CSx, and weighted MUSS. The reported $R^2$ values are calculated on an independent testing set of 10,000 individuals for each ancestry group. The corresponding 95% bootstrap CIs are obtained from the same testing set based on 10,000 bootstrap samples using the Bca approach[1] implemented in the R package "boot".

(a)

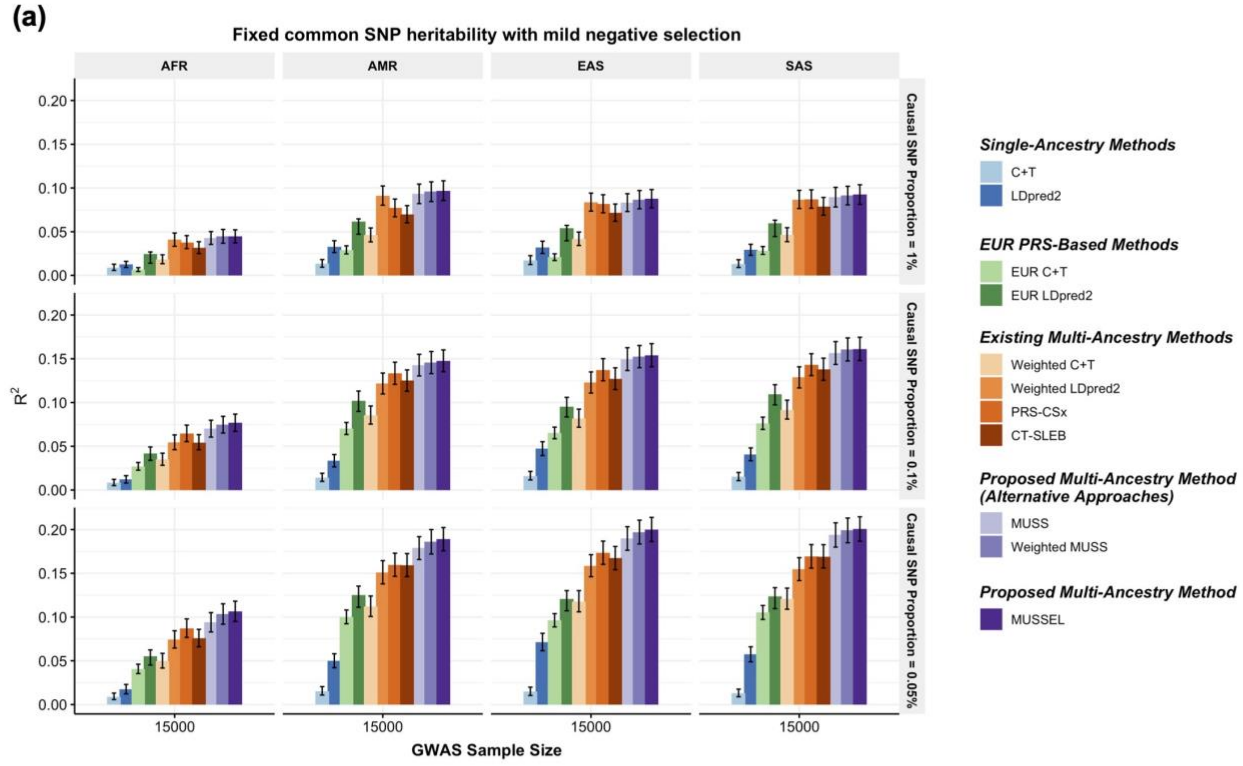
Fixed common SNP heritability with mild negative selection
AFR
AMR
EAS
SAS
Causal SNP Proportion = 1%
Causal SNP Proportion = 0.1%
Causal SNP Proportion = 0.05%
$R^2$
0.00
0.05
0.10
0.15
0.20
15000
GWAS Sample Size
Single-Ancestry Methods
C+T
LDpred2
EUR PRS-Based Methods
EUR C+T
EUR LDpred2
Existing Multi-Ancestry Methods
Weighted C+T
Weighted LDpred2
PRS-CSx
CT-SLEB
Proposed Multi-Ancestry Method (Alternative Approaches)
MUSS
Weighted MUSS
Proposed Multi-Ancestry Method
MUSSEL

(b)

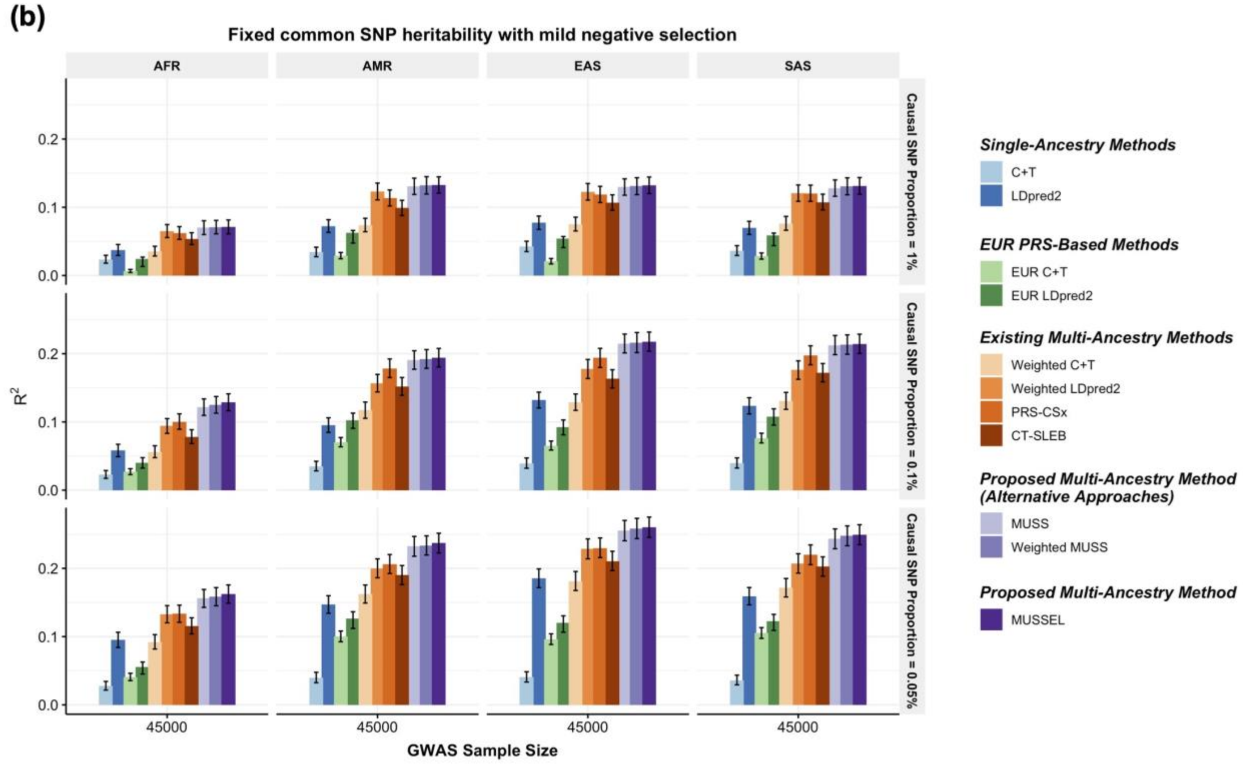
Fixed common SNP heritability with mild negative selection
AFR
AMR
EAS
SAS
Causal SNP Proportion = 1%
Causal SNP Proportion = 0.1%
Causal SNP Proportion = 0.05%
$R^2$
0.0
0.1
0.2
45000
GWAS Sample Size
Single-Ancestry Methods
C+T
LDpred2
EUR PRS-Based Methods
EUR C+T
EUR LDpred2
Existing Multi-Ancestry Methods
Weighted C+T
Weighted LDpred2
PRS-CSx
CT-SLEB
Proposed Multi-Ancestry Method (Alternative Approaches)
MUSS
Weighted MUSS
Proposed Multi-Ancestry Method
MUSSEL

**Supplementary Figure 10: Simulation results showing performance of the PRS constructed by MUSSEL and various existing methods, assuming a fixed common SNP heritability (0.4) across ancestries under a mild negative selection model for the relationship between SNP effect size and allele frequency.** The genetic correlation in SNP effect size is set to 0.8 across all pairs of populations. The causal SNP proportion (degree of polygenicity) is set to 1.0%, 0.1%, or 0.05% (~192$K$, 19.2$K$, or 9.6$K$ causal SNPs). We generate data for ~19 million common SNPs (MAF≥1%) across the five ancestries but conduct analyses only on the ~2.0 million SNPs in HapMap 3 + MEGA. The discovery GWAS sample size is set to **(a) 80,000 or (b) 100,000** for each non-EUR ancestry, and 100,000 for EUR. A tuning set consisting of 10,000 individuals is used for parameter tuning, as well as training the SL in CT-SLEB and MUSSEL or the linear combination model in weighted LDpred2, PRS-CSx, and weighted MUSS. The reported $R^2$ values are calculated on an independent testing set of 10,000 individuals for each ancestry group. The corresponding 95% bootstrap CIs are obtained from the same testing set based on 10,000 bootstrap samples using the Bca approach[1] implemented in the R package “boot”.

**(a)**

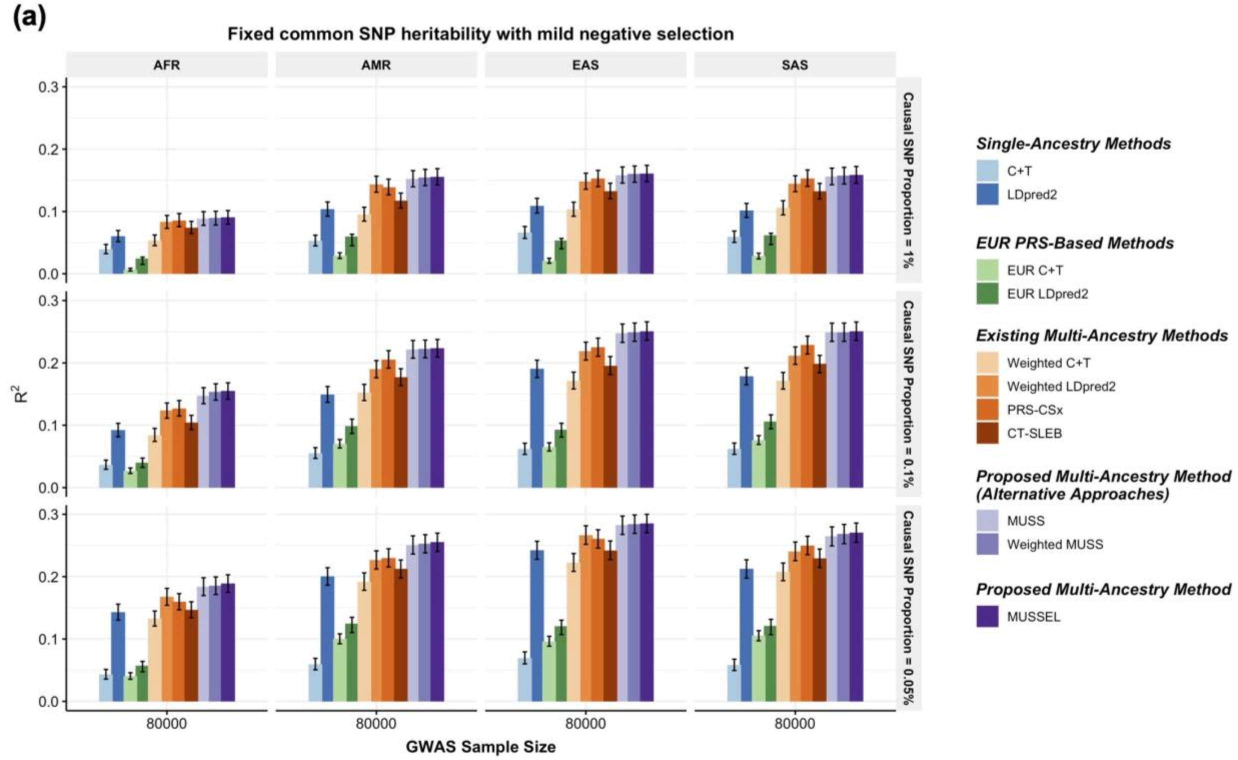

Fixed common SNP heritability with mild negative selection
AFR
AMR
EAS
SAS
Causal SNP Proportion = 1%
Causal SNP Proportion = 0.1%
Causal SNP Proportion = 0.05%
$R^2$
0.0
0.1
0.2
0.3
80000
GWAS Sample Size
Single-Ancestry Methods
C+T
LDpred2
EUR PRS-Based Methods
EUR C+T
EUR LDpred2
Existing Multi-Ancestry Methods
Weighted C+T
Weighted LDpred2
PRS-CSx
CT-SLEB
Proposed Multi-Ancestry Method (Alternative Approaches)
MUSS
Weighted MUSS
Proposed Multi-Ancestry Method
MUSSEL


**(b)**

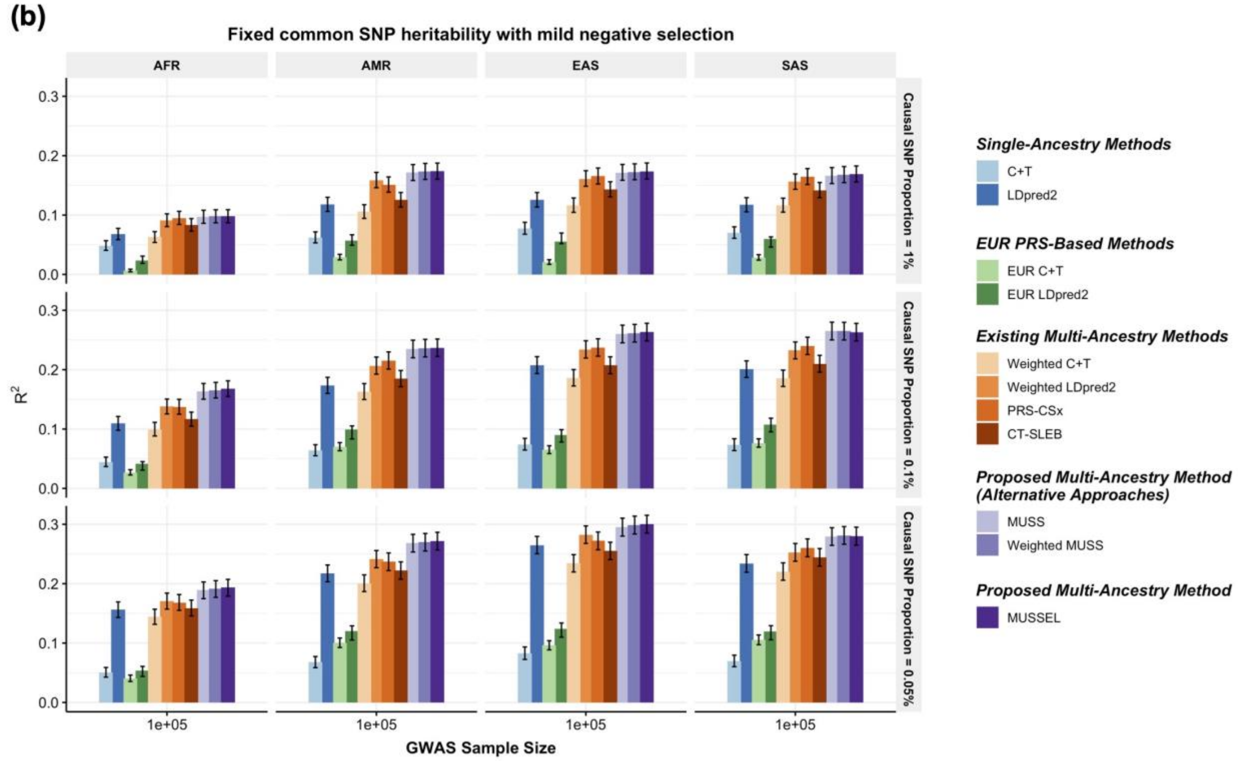

Fixed common SNP heritability with mild negative selection
AFR
AMR
EAS
SAS
Causal SNP Proportion = 1%
Causal SNP Proportion = 0.1%
Causal SNP Proportion = 0.05%
$R^2$
0.0
0.1
0.2
0.3
1e+05
GWAS Sample Size
Single-Ancestry Methods
C+T
LDpred2
EUR PRS-Based Methods
EUR C+T
EUR LDpred2
Existing Multi-Ancestry Methods
Weighted C+T
Weighted LDpred2
PRS-CSx
CT-SLEB
Proposed Multi-Ancestry Method (Alternative Approaches)
MUSS
Weighted MUSS
Proposed Multi-Ancestry Method
MUSSEL

**Supplementary Figure 11: Simulation results with 20% ancestry mis-specification in the LD reference sample, assuming a fixed common SNP heritability (0.4) across ancestries under a strong negative selection model for the relationship between SNP effect size and allele frequency.** The LD matrix for each ancestry group is estimated based on a slightly mis-specified LD reference sample that contains 800 individuals from the same ancestry group and 50 individuals from each of the other four ancestry groups, totaling 200 individuals with ancestry mismatch. The genetic correlation in SNP effect size is set to 0.8 across all pairs of populations. The causal SNP proportion (degree of polygenicity) is set to 1.0%, 0.1%, or 0.05% (~192$K$, 19.2$K$, or 9.6$K$ causal SNPs). We generate data for ~19 million common SNPs (MAF≥1%) across the five ancestry groups but conduct analyses only on the ~2.0 million SNPs in HapMap 3 + MEGA. The discovery GWAS sample size is set to **(a) 15,000 or (b) 45,000** for each non-EUR ancestry, and 100,000 for EUR. A tuning set consisting of 10,000 individuals is used for parameter tuning, as well as training the SL in CT-SLEB and MUSSEL or the linear combination model in weighted C+T, weighted LDpred2, PRS-CSx, and weighted MUSS. The reported $R^2$ values are calculated on an independent testing set of 10,000 individuals for each ancestry group.

**(a)**

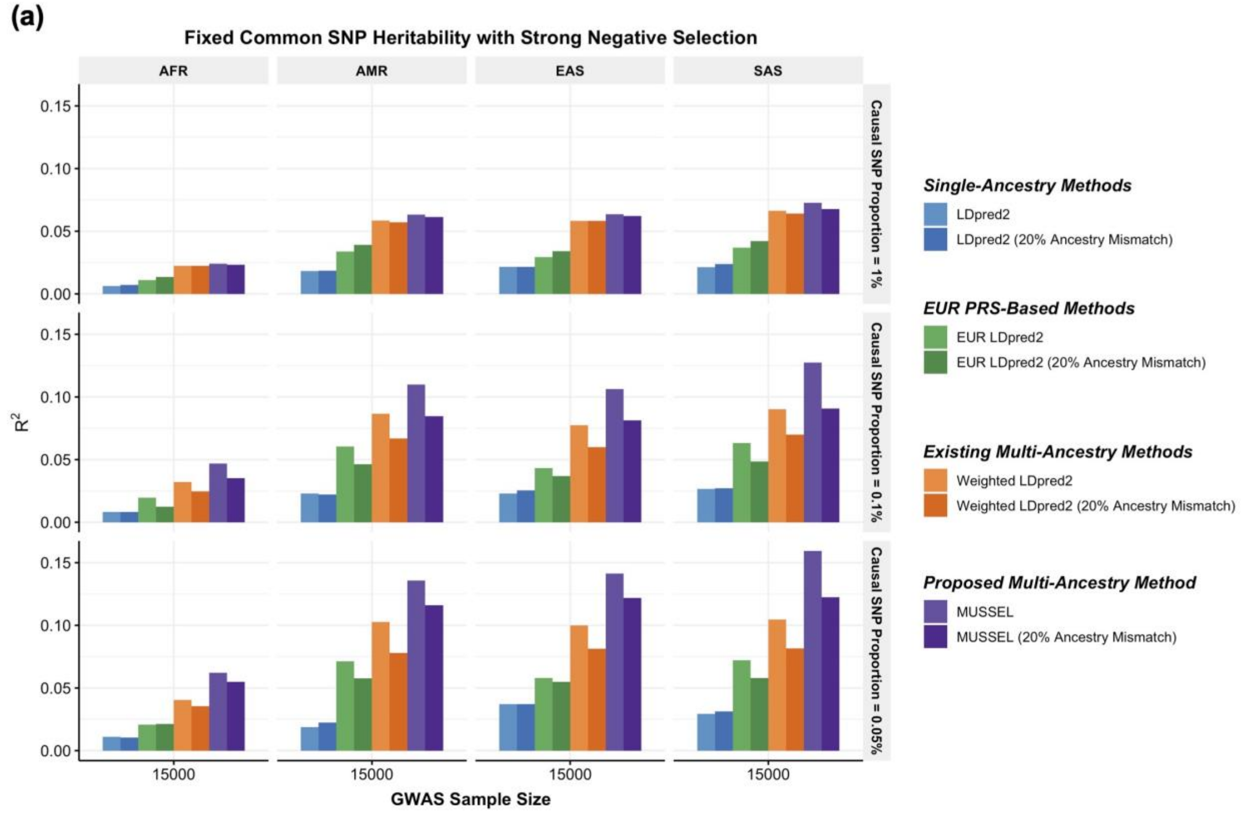

Fixed Common SNP Heritability with Strong Negative Selection
AFR
AMR
EAS
SAS
Causal SNP Proportion = 1%
Causal SNP Proportion = 0.1%
Causal SNP Proportion = 0.05%
$R^2$
0.15
0.10
0.05
0.00
15000
GWAS Sample Size
Single-Ancestry Methods
LDpred2
LDpred2 (20% Ancestry Mismatch)
EUR PRS-Based Methods
EUR LDpred2
EUR LDpred2 (20% Ancestry Mismatch)
Existing Multi-Ancestry Methods
Weighted LDpred2
Weighted LDpred2 (20% Ancestry Mismatch)
Proposed Multi-Ancestry Method
MUSSEL
MUSSEL (20% Ancestry Mismatch)


**(b)**

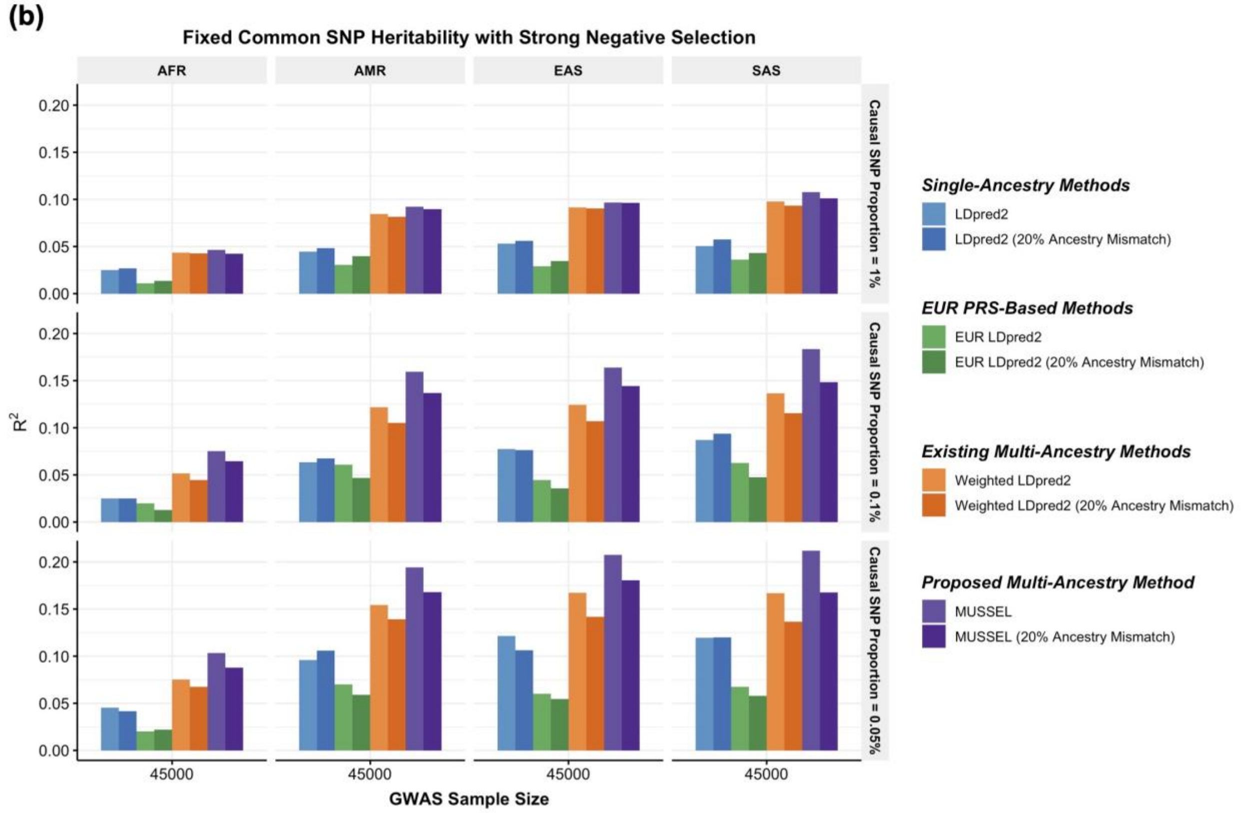

Fixed Common SNP Heritability with Strong Negative Selection
AFR
AMR
EAS
SAS
Causal SNP Proportion = 1%
Causal SNP Proportion = 0.1%
Causal SNP Proportion = 0.05%
$R^2$
0.20
0.15
0.10
0.05
0.00
45000
GWAS Sample Size
Single-Ancestry Methods
LDpred2
LDpred2 (20% Ancestry Mismatch)
EUR PRS-Based Methods
EUR LDpred2
EUR LDpred2 (20% Ancestry Mismatch)
Existing Multi-Ancestry Methods
Weighted LDpred2
Weighted LDpred2 (20% Ancestry Mismatch)
Proposed Multi-Ancestry Method
MUSSEL
MUSSEL (20% Ancestry Mismatch)

**Supplementary Figure 12: Simulation results with 20% ancestry mis-specification in the LD reference sample, assuming a fixed common SNP heritability (0.4) across ancestries under a strong negative selection model for the relationship between SNP effect size and allele frequency.** The LD matrix for each ancestry group is estimated based on a slightly mis-specified LD reference sample that contains 800 individuals from the same ancestry group and 50 individuals from each of the other four ancestry groups, totaling 200 individuals with ancestry mismatch. The genetic correlation in SNP effect size is set to 0.8 across all pairs of populations. The causal SNP proportion (degree of polygenicity) is set to 1.0%, 0.1%, or 0.05% (~192$K$, 19.2$K$, or 9.6$K$ causal SNPs). We generate data for ~19 million common SNPs (MAF≥1%) across the five ancestry groups but conduct analyses only on the ~2.0 million SNPs in HapMap 3 + MEGA. The discovery GWAS sample size is set to **(a) 80,000 or (b) 100,000** for each non-EUR ancestry, and 100,000 for EUR. A tuning set consisting of 10,000 individuals is used for parameter tuning, as well as training the SL in CT-SLEB and MUSSEL or the linear combination model in weighted C+T, weighted LDpred2, PRS-CSx, and weighted MUSS. The reported $R^2$ values are calculated on an independent testing set of 10,000 individuals for each ancestry group.

(a)

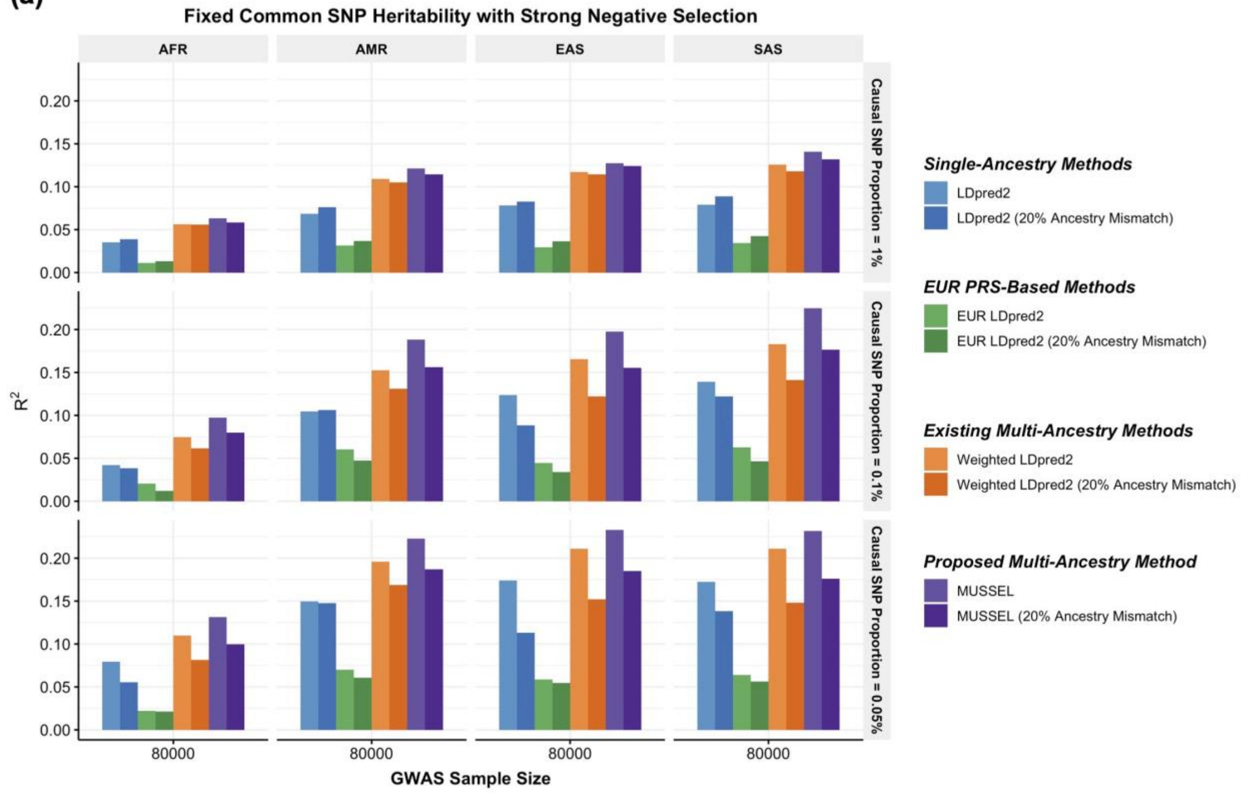

Fixed Common SNP Heritability with Strong Negative Selection
AFR
AMR
EAS
SAS
Causal SNP Proportion = 1%
Causal SNP Proportion = 0.1%
Causal SNP Proportion = 0.05%
R2
0.00
0.05
0.10
0.15
0.20
80000
GWAS Sample Size
Single-Ancestry Methods
LDpred2
LDpred2 (20% Ancestry Mismatch)
EUR PRS-Based Methods
EUR LDpred2
EUR LDpred2 (20% Ancestry Mismatch)
Existing Multi-Ancestry Methods
Weighted LDpred2
Weighted LDpred2 (20% Ancestry Mismatch)
Proposed Multi-Ancestry Method
MUSSEL
MUSSEL (20% Ancestry Mismatch)


(b)

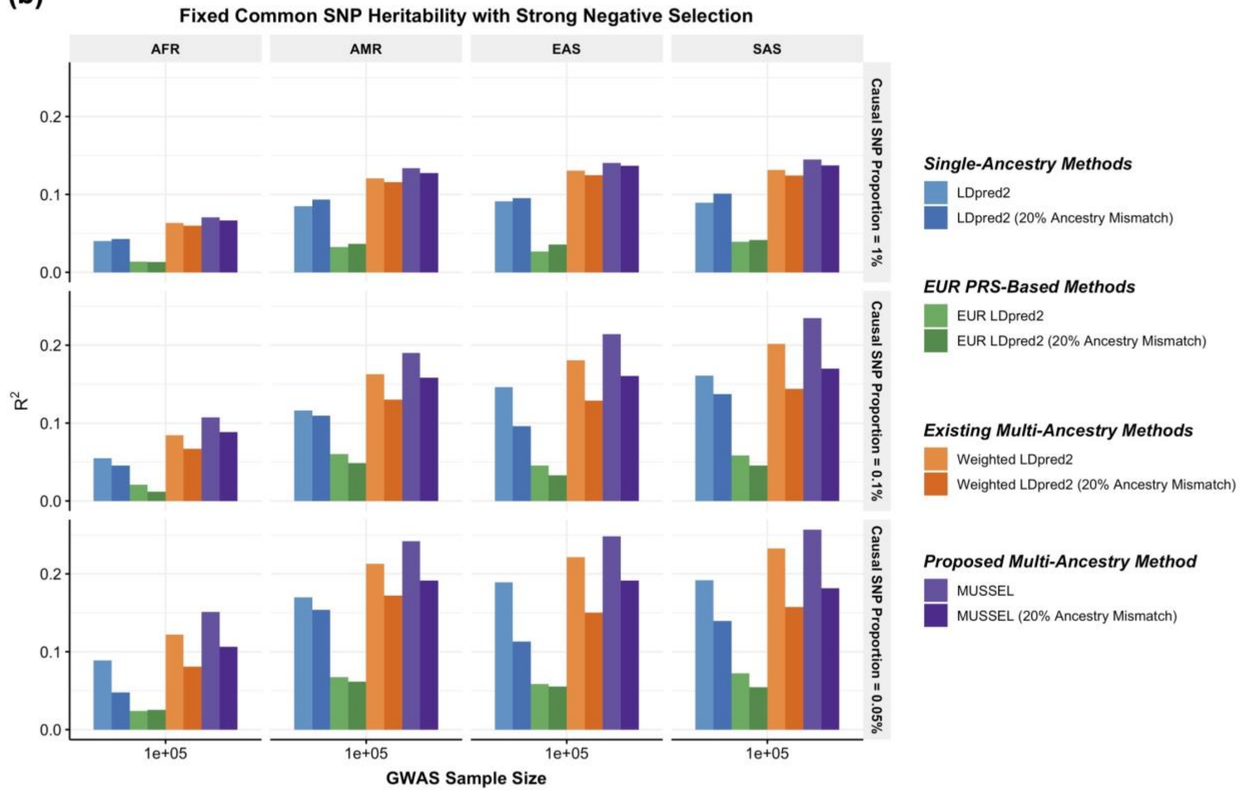

Fixed Common SNP Heritability with Strong Negative Selection
AFR
AMR
EAS
SAS
Causal SNP Proportion = 1%
Causal SNP Proportion = 0.1%
Causal SNP Proportion = 0.05%
R2
0.0
0.1
0.2
1e+05
GWAS Sample Size
Single-Ancestry Methods
LDpred2
LDpred2 (20% Ancestry Mismatch)
EUR PRS-Based Methods
EUR LDpred2
EUR LDpred2 (20% Ancestry Mismatch)
Existing Multi-Ancestry Methods
Weighted LDpred2
Weighted LDpred2 (20% Ancestry Mismatch)
Proposed Multi-Ancestry Method
MUSSEL
MUSSEL (20% Ancestry Mismatch)

**Supplementary Figure 13: Prediction $R^2$ with 95% bootstrap CIs on validation individuals of AFR (N=2,015–3,428), EAS (N=2,316-4,647), and AMR ancestries (N=3,479-4,397) in PAGE based on discovery GWAS from PAGE (AFR $N_{GWAS}$=7,775 – 13,699, AMR $N_{GWAS}$=13,894 – 17,558), BBJ (EAS $N_{GWAS}$=70,657 – 158,284), and UKBB (EUR $N_{GWAS}$=315,133 – 355,983).** We used genotype data from 1000 Genomes Project (498 EUR, 659 AFR, 347 AMR, 503 EAS, 487 SAS) as the LD reference dataset. All methods were evaluated on the ~2.0 million SNPs that are available in HapMap 3 + MEGA, except for PRS-CSx which is evaluated based on the HapMap 3 SNPs only, as implemented in their software. Ancestry- and trait-specific GWAS sample sizes, number of SNPs included, and validation sample sizes are summarized in Supplementary Table 3.1. A random half of the validation individuals is used as the tuning set to tune model parameters, as well as train the SL in CT-SLEB and MUSSEL or the linear combination model in weighted C+T, weighted LDpred2, PRS-CSx, and weighted MUSS. The other half of the validation set is used as the testing set to report $R^2$ values for PRS on each ancestry, after adjusting for whether or not the sample is from BioMe and the top 10 genetic principal components for BMI, and additionally the age at lipid measurement and sex. The 95% bootstrap CIs of the estimated $R^2$ are obtained from the testing set based on 10,000 bootstrap samples using the Bca approach[1] implemented in the R package “boot”. Detailed 95% bootstrap CIs are reported in Supplementary Table 9.

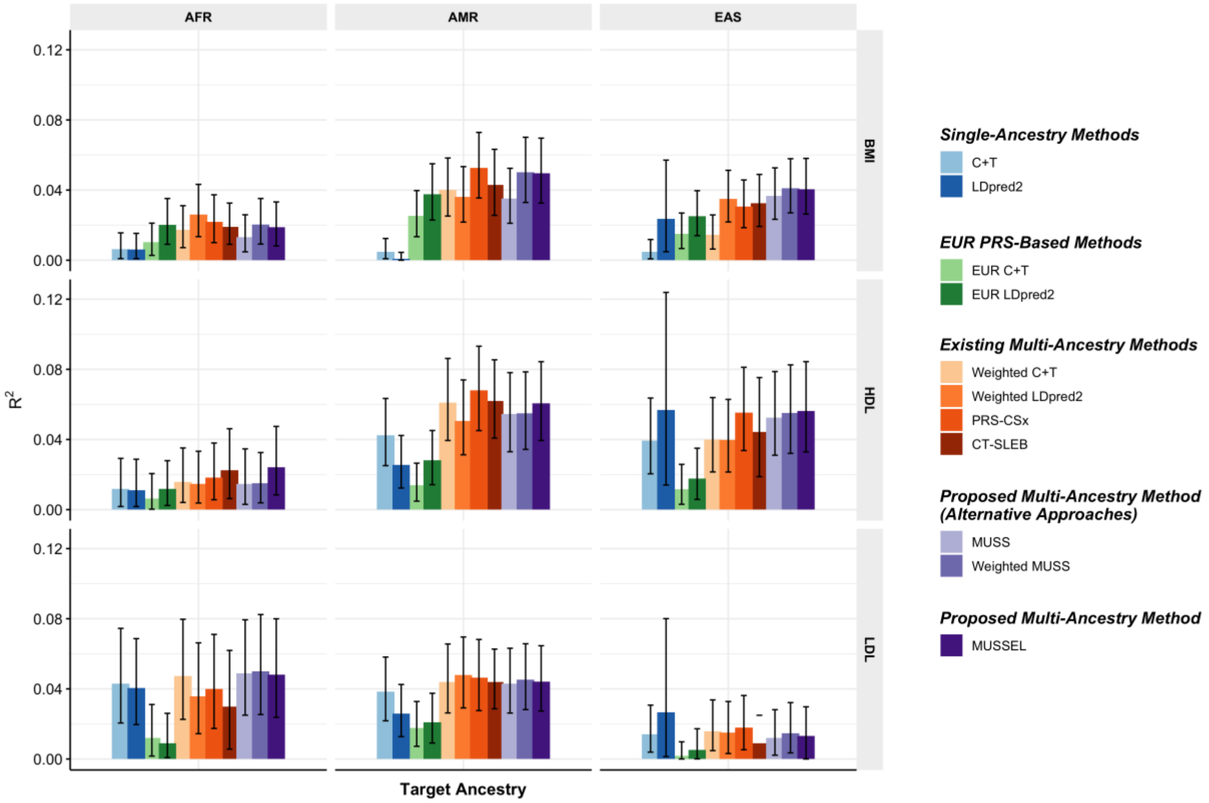

**Supplementary Figure 14: Prediction $R^2$ with 95% bootstrap CIs on UKBB validation individuals of EUR (17,457 – 19,030), AFR (7,954 – 8,598), EAS (1,752 – 1,921), and SAS (9,385 – 10,288) origin based on discovery GWAS from GLGC on EUR ($N_{GWAS}$ =842,660 – 930,671), AFR or admixed AFR ($N_{GWAS}$ =87,760 – 92,555), Hispanic/Latino ($N_{GWAS}$ =46,040 – 49,582), EAS ($N_{GWAS}$ =82,587 – 146,492), and SAS ($N_{GWAS}$ =33,658 – 34,135). EUR ($N_{GWAS}$ =842,660 – 930,671), AFR or admixed AFR ($N_{GWAS}$ =87,760 – 92,555), Hispanic/Latino ($N_{GWAS}$ =46,040 – 49,582), EAS ($N_{GWAS}$ =82,587 – 146,492), and SAS ($N_{GWAS}$ =33,658 – 34,135).** The LD reference data is either (a) 1000 Genomes Project (498 EUR, 659 AFR, 347 AMR, 503 EAS, 487 SAS), or (b) UKBB data (PRS-CSx: default UKBB LD reference data which overlap with our testing samples including 375,120 EUR, 7,507 AFR, 687 AMR, 2,181 EAS, and 8,412 SAS; all other methods: UKBB tuning samples including 10,000 EUR, 4,585 AFR, 1,010 EAS, and 5,427 SAS). The ancestry of UKBB individuals were determined by a genetic ancestry prediction approach (Supplementary Notes). Due to the low prediction accuracy of genetic component analysis and extremely small validation sample size of UKBB AMR, prediction $R^2$ on UKBB AMR is unreliable and thus is not reported here. All methods were evaluated on the ~2.0 million SNPs that are available in HapMap 3 + MEGA, except for PRS-CSx which is evaluated based on the HapMap 3 SNPs only, as implemented in their software. Ancestry- and trait-specific GWAS sample sizes, number of SNPs included, and validation sample sizes are summarized in Supplementary Table 4.1. A random half of the validation individuals is used as the tuning set to tune model parameters, as well as train the SL in CT-SLEB and MUSSEL or the linear combination model in weighted LDpred2, PRS-CSx, and weighted MUSS. The other half of the validation set is used as the testing set to report $R^2$ values for each ancestry. The 95% bootstrap CIs of the estimated $R^2$ are obtained from the testing set based on 10,000 bootstrap samples using the Bca approach[1] implemented in the R package "boot". Detailed 95% bootstrap CIs are reported in Supplementary Table 9. In (b), PRS-CSx and other methods do not have a fair comparison because the UKBB LD reference data provided by the PRS-CSx software ($UKBB_{PRS\text{-}CSx}$) is much larger than that for other methods, and thus the $R^2$ of PRS-CSx PRS may be inflated due to a big overlap between $UKBB_{PRS\text{-}CSx}$ and the UKBB testing sample.

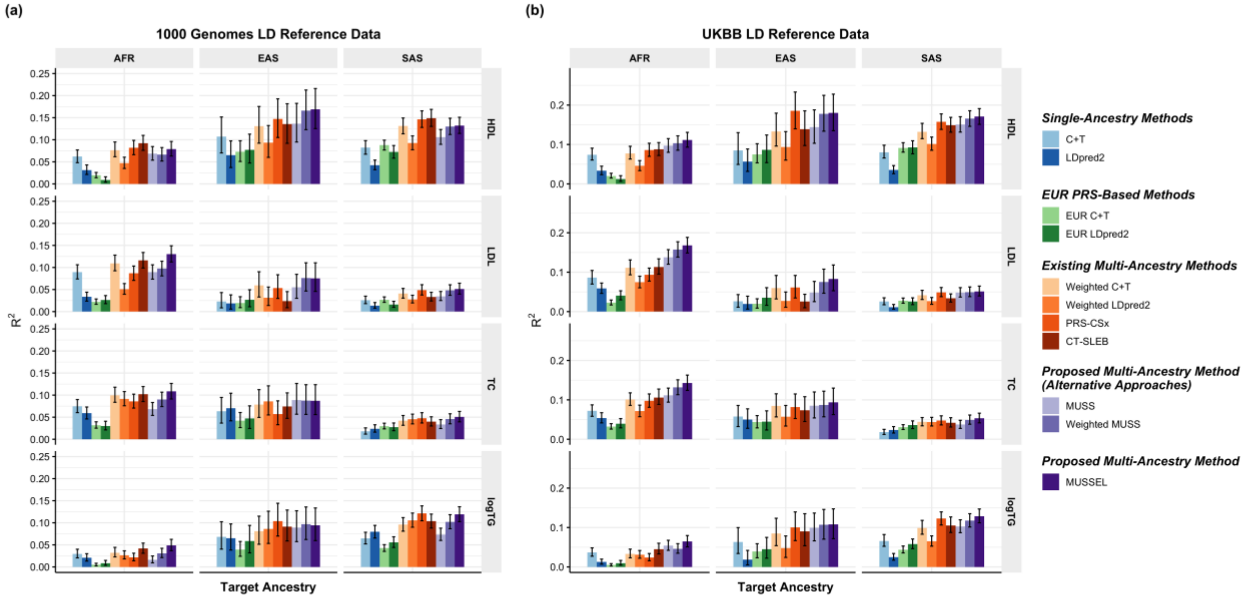

**Supplementary Figure 15: Prediction $R^2$ with 95% bootstrap CIs on UKBB validation individuals of AFR (N=9,169) origin based on discovery GWAS from GLGC on EUR ($N_{GWAS}$ =48,223 – 48,326), AFR ($N_{GWAS}$ =21,511 – 21,547), and Hispanic/Latino ($N_{GWAS}$ =15,362 – 15,411).** The LD reference data is either **(a)** 1000 Genomes Project (498 EUR, 659 AFR, 347 AMR, 503 EAS, 487 SAS), or **(b)** UKBB data (PRS-CSx: default UKBB LD reference data which overlap with our testing samples including 375,120 EUR, 7,507 AFR, 687 AMR, 2,181 EAS, and 8,412 SAS; all other methods: UKBB tuning samples including 10,000 EUR, 4,585 AFR, 1,010 EAS, and 5,427 SAS). The ancestry of UKBB individuals were determined by a genetic ancestry prediction approach (Supplementary Notes). Due to the low prediction accuracy of genetic component analysis and extremely small validation sample size of UKBB AMR, prediction $R^2$ on UKBB AMR is unreliable and thus is not reported here. All methods were evaluated on the ~2.0 million SNPs that are available in HapMap3 + MEGA, except for PRS-CSx which is evaluated based on the HapMap 3 SNPs only, as implemented in their software. Ancestry- and trait-specific sample sizes of GWAS, number of SNPs included, and validation sample sizes are summarized in Supplementary Table 5.1. A random half of the validation individuals is used as the tuning set to tune model parameters, as well as train the SL in CT-SLEB and MUSSEL or the linear combination model in weighted LDpred2, PRS-CSx, and weighted MUSS. The other half of the validation set is used as the testing set to report $R^2$ values for each ancestry, after adjusting for age, sex, and the top 10 genetic principal components. The 95% bootstrap CIs of the estimated $R^2$ are obtained from the testing set based on 10,000 bootstrap samples using the Bca approach[1] implemented in the R package "boot". Detailed 95% bootstrap CIs are reported in Supplementary Table 9. In (b), PRS-CSx and other methods do not have a fair comparison because the UKBB LD reference data provided by the PRS-CSx software ($UKBB_{PRS\text{-}CSx}$) is much larger than that for other methods, and thus the $R^2$ of PRS-CSx may be inflated due to a big overlap between $UKBB_{PRS\text{-}CSx}$ and the UKBB testing sample.

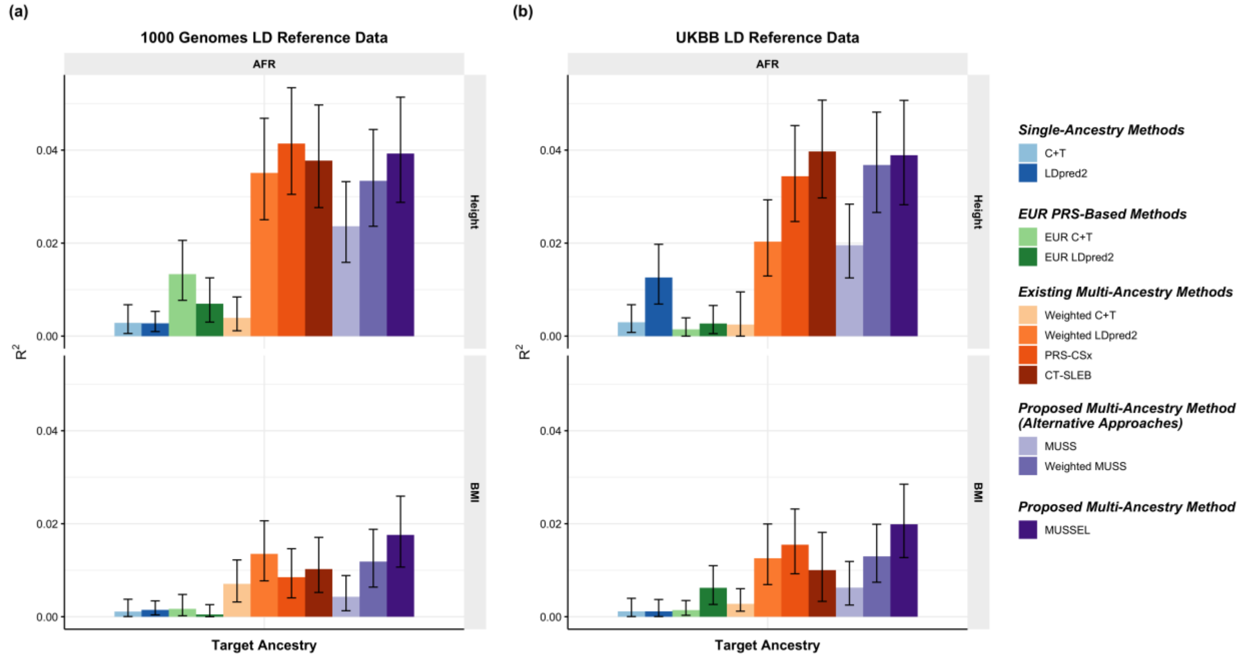

**Supplementary Figure 16:** Manhattan plot and QQ plot[1] based on the GWAS summary-level association statistics from PAGE for BMI in four populations: European, Admixed African or African, Hispanic, and East Asian.

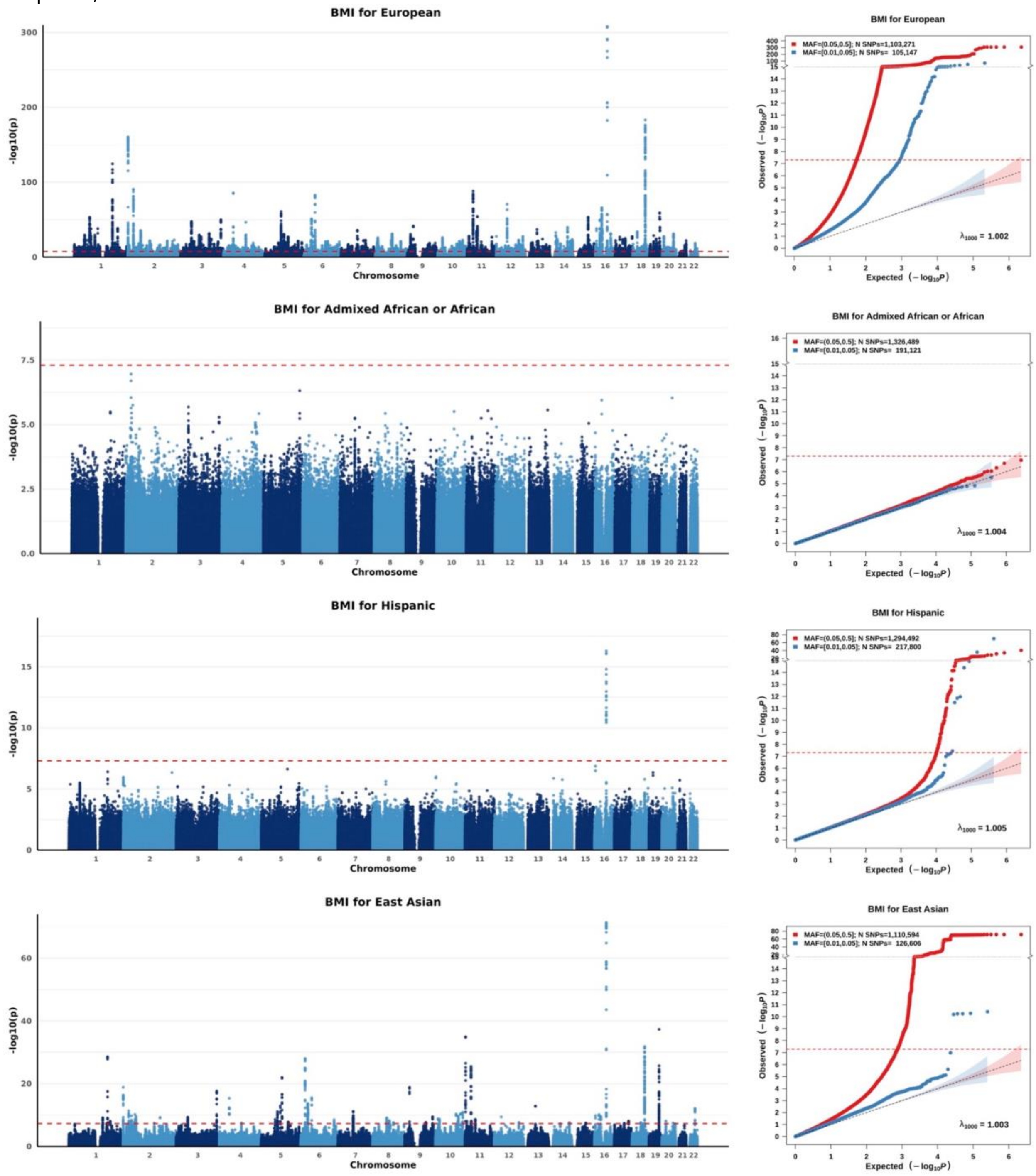


**1** For continuous traits, $\lambda_{1000}$ scales the genomic inflation factor $\lambda$ to a study with 1000 subjects using $\lambda_{1000} = 1 + 1000(\lambda - 1)/N$, where N is the total sample size. For binary traits, $\lambda_{1000}$ scales $\lambda$ to a study with 1000 cases and 1000 controls using $\lambda_{1000} = 1 + 1000(\lambda - 1)(\frac{1}{N_{case}} + \frac{1}{N_{control}})$.

**Supplementary Figure 17:** Manhattan plot and QQ plot[1] based on the GWAS summary-level association statistics from PAGE for high-density lipoprotein (HDL) in four populations: European, Admixed African or African, Hispanic, and East Asian.

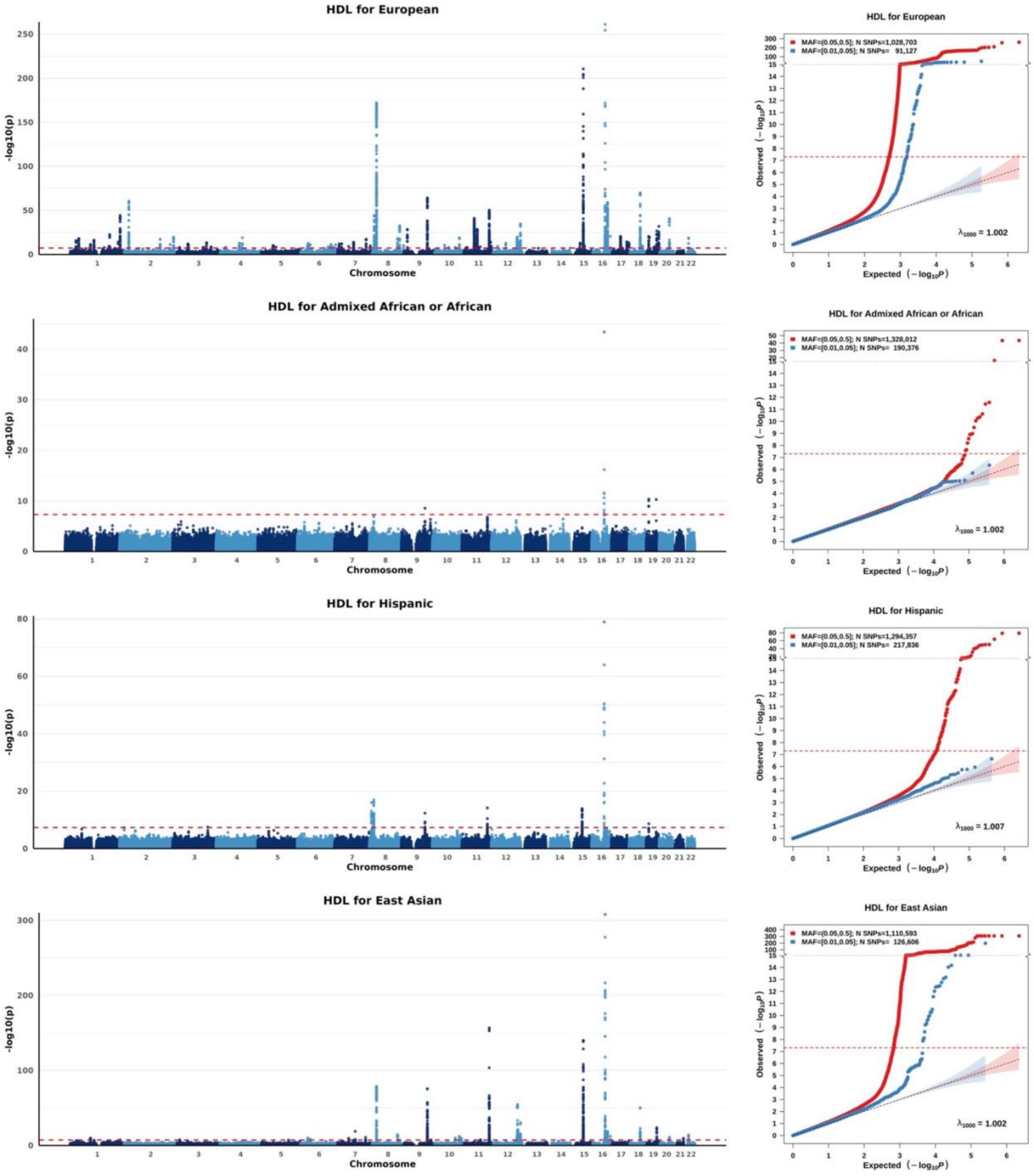


[1] For continuous traits, $\lambda_{1000}$ scales the genomic inflation factor $\lambda$ to a study with 1000 subjects using $\lambda_{1000} = 1 + 1000(\lambda - 1)/N$, where N is the total sample size. For binary traits, $\lambda_{1000}$ scales $\lambda$ to a study with 1000 cases and 1000 controls using $\lambda_{1000} = 1 + 1000(\lambda - 1)(\frac{1}{N_{case}} + \frac{1}{N_{control}})$.

**Supplementary Figure 18:** Manhattan plot and QQ plot[1] based on the GWAS summary-level association statistics from PAGE for low-density lipoprotein (LDL) in four populations: European, Admixed African or African, Hispanic, and East Asian.

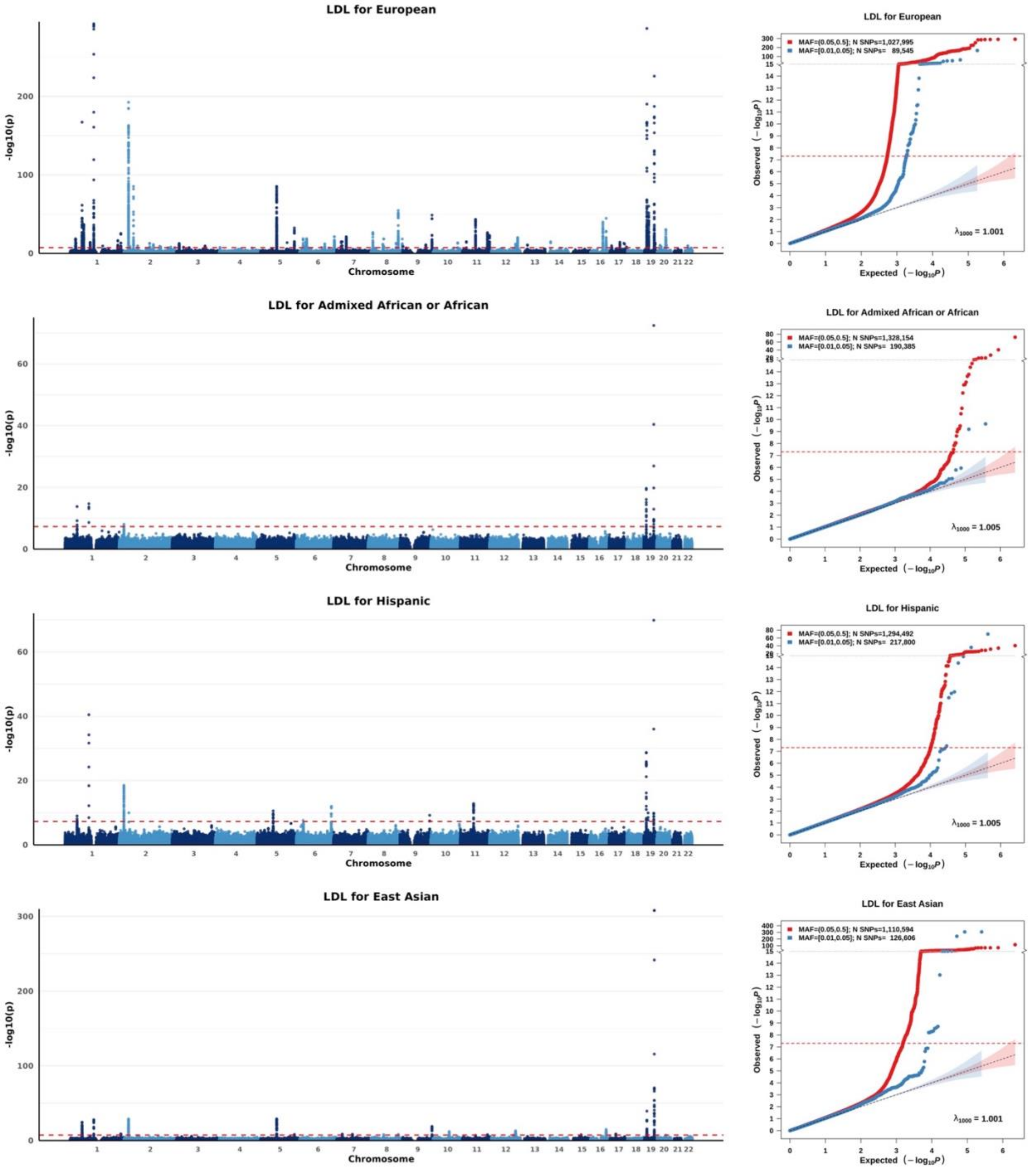


[1] For continuous traits, $\lambda_{1000}$ scales the genomic inflation factor $\lambda$ to a study with 1000 subjects using $\lambda_{1000} = 1 + 1000(\lambda - 1)/N$, where N is the total sample size. For binary traits, $\lambda_{1000}$ scales $\lambda$ to a study with 1000 cases and 1000 controls using $\lambda_{1000} = 1 + 1000(\lambda - 1)(\frac{1}{N_{case}} + \frac{1}{N_{control}})$.

### Supplementary Notes

#### Detailed simulation setup

We investigated the performance of MUSSEL and a series of existing methods under various simulated scenarios of genetic architecture for phenotype and GWAS sample sizes across ancestries. This large-scale, multi-ancestry simulated dataset including 600,000 individuals across EUR, AFR, AMR, EAS, and SAS origins has recently been released by our group[1]. Specifically, the genotype data was simulated using HAPGEN2 (version 2.1.2)[2] based on the genotype data of 2,504 unrelated individuals from Phase 3 1000 Genomes Project (503 EUR, 661 AFR, 347 AMR, 504 EAS, and 489 SAS)[3]. We have checked and confirmed the consistency between the LD pattern in the original 1000 Genomes reference data and the LD pattern in our simulated data{Zhang, 2022 #360}. Approximately 19.2 million common biallelic SNPs with MAF≥0.01 in at least one ancestry group were included. For phenotype data, genetic architectures were simulated by first selecting a random set of 1.0%, 0.1%, or 0.05% SNPs across the whole genome to be causal, that is approximately 192$K$, 19.2$K$, or 9.6$K$ causal SNPs among 19.2 million SNPs. Under a spike and slab structure, the nonzero standardized effect sizes for the causal SNPs were then generated under various negative selection models according to a function of allele frequency, $\beta_{kj}^{(J)} \propto \{q_{kj}(1-q_{kj})\}^{\alpha}$: (1) strong negative selection: $\alpha = 0$, (2) mild negative selection: $\alpha = 0.75$, or (3) no negative selection, $\alpha = 1$. The genetic correlation was set to $\rho = 0.8$ or 0.6 between all pairs of ancestries. Specifically, we first generated $\nu_{kj} \sim N(0, H_k^2/m_k)$ for SNPs only existing in ancestry $k$, with $cov(\nu_{kj}, \nu_{k'j}) = \rho H_k H_{k'}/m_k m_{k'}$ for SNPs shared between ancestries $k$ and $k'$, where $H_k^2$ and $m_k$ denote the total heritability and the number of causal SNPs, respectively, in ancestry $k$. To control the total heritability at the predefined level $H_k^2$s, we set the standardized SNP effect sizes to $\beta_{kj}^{(J)} = \{q_{kj}(1-$

$q_{kj})\}^{\alpha}\nu_{kj}\sqrt{H_k^2/\sum_{j=1}^{m_k}[\{q_{kj}(1-q_{kj})\}^{\alpha}\nu_{kj}]^2}$. Two heritability settings were considered: (1) a constant common SNP heritability 0.4 across all ancestries, and (2) a total heritability of 0.4 across all 19.2 million SNPs with a constant per-SNP heritability across ancestries, which leads to a common SNP heritability proportional to the number of common SNPs in the corresponding ancestry.

We simulated 120,000 individuals for each ancestry. For EUR, $N_{GWAS}$=100,000 individuals were included in the discovery GWAS, while the remaining 20,000 individuals were evenly split into a tuning set for parameter tuning and a testing set to report prediction $R^2$ of the methods. For each non-EUR ancestry, $N_{GWAS}$ individuals were included in the discovery GWAS, while two separate sets, each including 10,000 individuals, were selected randomly from the remaining (120,000 – $N_{GWAS}$) individuals to construct tuning and testing dataset. Although currently the non-EUR GWAS sample sizes are typically a lot smaller than EUR GWAS sample sizes, they are expected to continue growing, as there is an increasing emphasis on health equity. To mimic such real-world scenarios, we set non-EUR GWAS sample sizes to $N_{GWAS}$ = 15,000, 45,000, 80,000, or 100,000, that gradually increase and eventually reach a similar level to the EUR GWAS sample size (100,000). For each ancestry group, the genotype data of 1000 randomly selected individuals in the discovery GWAS were used to estimate the ancestry-specific LD.

**Predicted genetic ancestry for non-EUR individuals in UKBB**

We compute genetic ancestry for all UKBB individuals that are not self-reported Whites. To balance between samples of different ancestry groups, we also include 8,000 unrelated self-reported Whites to form the set of UKBB individuals for genetic ancestry prediction. We use

2,504 unrelated individuals from 1000 Genomes Project, including 498 EUR, 659 AFR, 347 AMR, 503 EAS, and 487 SAS individuals to form the reference data for genetic ancestry prediction. We first compute the top 20 genetic principal components (PCs) for all UKBB and 1000 Genomes individuals together using PLINK 2.0 command --pca 20 allele-wts[5]. We then train a random forest classifier with 1,500 trees using the R package “randomForest”[6] based on the genetic PCs of the 1000 Genomes individuals with their true labels being provided by gnomAD[7] that can be used to capture enough ancestral information. Finally, we apply the trained random forest classifier to predict the genetic ancestry of UKBB individuals based on their genetic PCs.

References


1. DiCiccio TJ, Efron B. Bootstrap Confidence Intervals. Statistical Science 1996;11:189-212.
2. Zhang H, Zhan J, Jin J, et al. A new Method for Multi-ancestry Polygenic Prediction Improves Performance across Diverse Populations. bioRxiv 2022:2022.03.24.485519.
3. Su Z, Marchini J, Donnelly P. HAPGEN2: simulation of multiple disease SNPs. Bioinformatics 2011;27:2304-5.
4. Genomes Project C, Auton A, Brooks LD, et al. A global reference for human genetic variation. Nature 2015;526:68-74.
5. Shaun Purcell CC. PLINK 2.0. URL: www.cog-genomics.org/plink/2.0/.
6. Liaw A, Wiener M. Classification and regression by randomForest. R news 2002;2:18-22.
7. Karczewski KJ, Francioli LC, Tiao G, et al. The mutational constraint spectrum quantified from variation in 141,456 humans. Nature 2020;581:434-43.
